# Catalytic domain plasticity of MKK7 reveals structural mechanisms of allosteric activation and new targeting opportunities

**DOI:** 10.1101/2020.06.11.145995

**Authors:** Martin Schröder, Li Tan, Jinhua Wang, Yanke Liang, Nathanael S. Gray, Stefan Knapp, Apirat Chaikuad

**Author notes:** Authors for correspondence: Stefan Knapp;, and Apirat Chaikuad;. Lead contact: Apirat Chaikuad;.

## Abstract

MKK7 (MEK7) is a key regulator of the JNK stress signaling pathway and targeting MKK7 has been proposed as a chemotherapeutic strategy. Detailed understanding of the MKK7 structure and factors that impact its activity is therefore of critical importance. Here, we present a comprehensive set of MKK7 crystal structures revealing insights into catalytic domain plasticity and the role of the N-terminal regulatory helix, conserved in all MAP2Ks, mediating kinase activation. Crystal structures harboring this regulatory helix revealed typical structural features of active kinase, providing exclusively a first model of the MAP2K active state. A small molecule screening campaign yielded multiple scaffolds, including type-II irreversible inhibitors a binding mode that has not been reported previously. We also observed an unprecedented allosteric pocket located in the N-terminal lobe for the approved drug ibrutinib. Collectively, our structural and functional data expand and provide alternative targeting strategies for this important MAP2K kinase.

## Introduction

Dual specific mitogen-activated protein kinase kinase 7 (MKK7), also known as MEK7 or MAP2K7, is a member of mitogen-activated kinase kinase (MAP2K) subfamily, and a key activator of c-Jun N-terminal kinase (JNK) signaling, a pathway that regulates primarily stress and inflammatory responses^*1*^. MKK7 shares this signaling role with the related MAP2K MKK4 as both kinases are able to phosphorylate and activate JNKs. However, in contrast to other MAP2K members, MKK7 harbors three docking domains (D domain) at its N-terminus, which cooperatively and selectively interact with JNKs enabling efficient phosphorylation of the TxY motif threonine that is essential for regulating JNK activity^*2-4*^. The activation of the MKK7 kinase domain itself requires phosphorylation of serine and threonine residues within the SKAKT motif^*5*^. MKK7 auto-inhibition is controlled by several factors, including the N-terminal regulatory helix^*6, 7*^. However, molecular details of the transition into the catalytically active state remained unknown for MKK7 as well as for other MAP2Ks.

Many studies have demonstrated that MKK7 and JNK signaling play crucial roles in the development of a variety of human diseases, including inflammatory diseases^*8, 9*^, Alzheimer^*10, 11*^ as well as cancer^*12-15*^. Several lines of evidence have linked elevated MKK7 levels to the development of metastasis and progression of cancer^*16, 17*^. The aberrantly activated JNK cascade may also contribute to the resistance of tumors to anti-cancer drugs including 5-fluorouracil, DNA-damaging agents and the kinase inhibitor sorafenib^*18-20*^. Furthermore, JNK signaling has been recently implicated in Zika virus^*21*^ and Hepatitis B^*22*^ infections. These data make a compelling case for targeting MKK7 as a therapeutic strategy for these diverse diseases. However, several reports have proposed also tumor suppressor functions of MKK7 ^*23-26*^, suggesting that the roles of MMK7 vary dramatically depending on stress stimuli, cell types, and disease mechanisms in different contexts^*12, 27*^. A highly selective inhibitor of MKK7 would represent an important tool to address the role of MKK7 in the above mentioned pathologies.

Previous efforts to develop potent and selective inhibitors targeting JNK pathway have mostly focused on direct targeting of JNK family members, exemplified by several recent potent inhibitors including CC-930^*28*^, JNK-IN-8^*29*^ and LN2332 (compound 7)^*30*^. However, these inhibitors lack isoform selectivity and have been associated with cellular toxicity potentially limiting chemotherapeutic benefit^*31*^. To complement these efforts and to overcome some of these challenges, targeting MKK4 and MKK7 has been suggested as an alternative strategy for inhibiting JNK signaling. Inhibition of these MAP2Ks may provide benefits by maintaining some basal activity of JNKs and reducing potential toxicity. Given that MKK4 is additionally involved in p38 pathway^*32*^, it would be expected that targeting MKK4 may lead to more pleiotropic effects in comparison to targeting MKK7 which is a JNK specific activator.

Previous structural reports have suggested that accessibility of MKK7 ATP binding site might be constrained by an auto-inhibition mechanism^*6, 7*^. However, the high degree of plasticity of the kinase in particular in its inactive state suggests possibilities for the accommodation of small molecule binders^*7, 33*^. In addition, the presence of accessible cysteine residues within the MKK7 ATP binding pocket opens also an opportunity for irreversible targeting strategy^*34*^, supported by previous observations of covalent binding the natural product 5Z-7-oxzeaenol and promiscuous aminopyrimidine SM1-71 with the kinase domain^*35-37*^. In addition, previous studies have demonstrated that ibrutinib which is an BTK inhibitor can interact covalently with MKK7 in cells ^*37, 38*^. The development of ATP competitive kinase inhibitors often follows two main design strategies: Type-I inhibitors target the active state and are anchored to the kinase hinge region whereas type-II inhibitors may also interact with the kinase hinge but they target an inactive state characterized by an outward flipped conformation of the conserved tripeptide motif DFG (DFG-out). To date, irreversible targeting using type-I inhibitors has exclusively been exploited for this kinase, leading to covalent indazole-based derivatives^*39*^ and 4-amino-pyrazolopyrimidine-based inhibitors^*33, 40*^. However, despite nanomolar *in vitro* inhibitory potencies, the cellular activities of these inhibitors are still limited to the micromolar activity range, highlighting the need for additional inhibitor scaffolds with improved cellular activity and further optimization.

An often used strategy to covalent inhibitor development involves the identification of potent reversible inhibitors and subsequent installation of a suitable electrophilic warhead. Thus, a better understanding of the structural plasticity of MKK7 ATP binding pocket as well as identification of suitable reversible inhibitor with diverse binding modes will undoubtedly facilitate covalent targeting approach. To address these challenges, we determined multiple crystal structured of MKK7, which revealed molecular mechanisms of MKK7 activation, a long standing question for the MAP2K family in general, as well as identified alternative strategies such as allosteric pockets that could be utilized for type-III targeting strategies. Interestingly, we found that MKK7 comprises an N-terminal regulatory helix, resembling that of MEK1/2, which acts allosterically by facilitating a transition from an inactive, auto-inhibitory state to an active conformation. In addition, our inhibitor screening revealed that MKK7 can be targeted by both type-I and type-II inhibitors with reversible and irreversible modes of action. We observed also the presence of an unexpected, additional binding site that may be used for the development of allosteric inhibitors using ibrutinib as a starting point. Thus, the structural data presented here offer multiple new targeting strategies for future development of highly selective MKK7 inhibitors and provide insight in the regulation of this important MAP2K.

## Results

### Structure of the MKK7 kinase domain revealed an auto-inhibitory conformation

To provide structural insights into the kinase domain of MKK7, we first determined the crystal structure of wild-type ΔN115-MKK7 in its apo, non-phosphorylated form (Figure 1A and Supplementary table 1). This structure revealed an inactive state with several catalytic motifs displaying catalytically-incompetent conformations, including an “out” conformation of the DFG motif and the absence of the canonical salt bridge of the β3 “VIAK” motif lysine (K165) with the αC helix glutamate. Strikingly, both β1 and β2 strands were distorted, resulting also in distortion of the P-loop that consequently protruded into the ATP binding site limiting access to the co-factor. To investigate if this inactive conformation might be a structural feature of the non-phosphorylated form of the kinase, we crystallized the phosphorylation-mimetic mutant harboring S287D and T291D substitutions (MKK7, DD). Despite slightly different P-loop conformation, the mutant, however, still assumed an inactive state characterized by occlusion of the ATP site resembling the conformation observed in the wild-type structure (Figure 1B), suggesting that these mutations were not sufficient to induce an active conformation. In agreement with previous studies ^*7*^, the persistently inactive conformations observed in MKK7 apo-structures suggest additional auto-regulation mechanism that are essential for controlling the kinase catalytic function.

**Figure 1:**
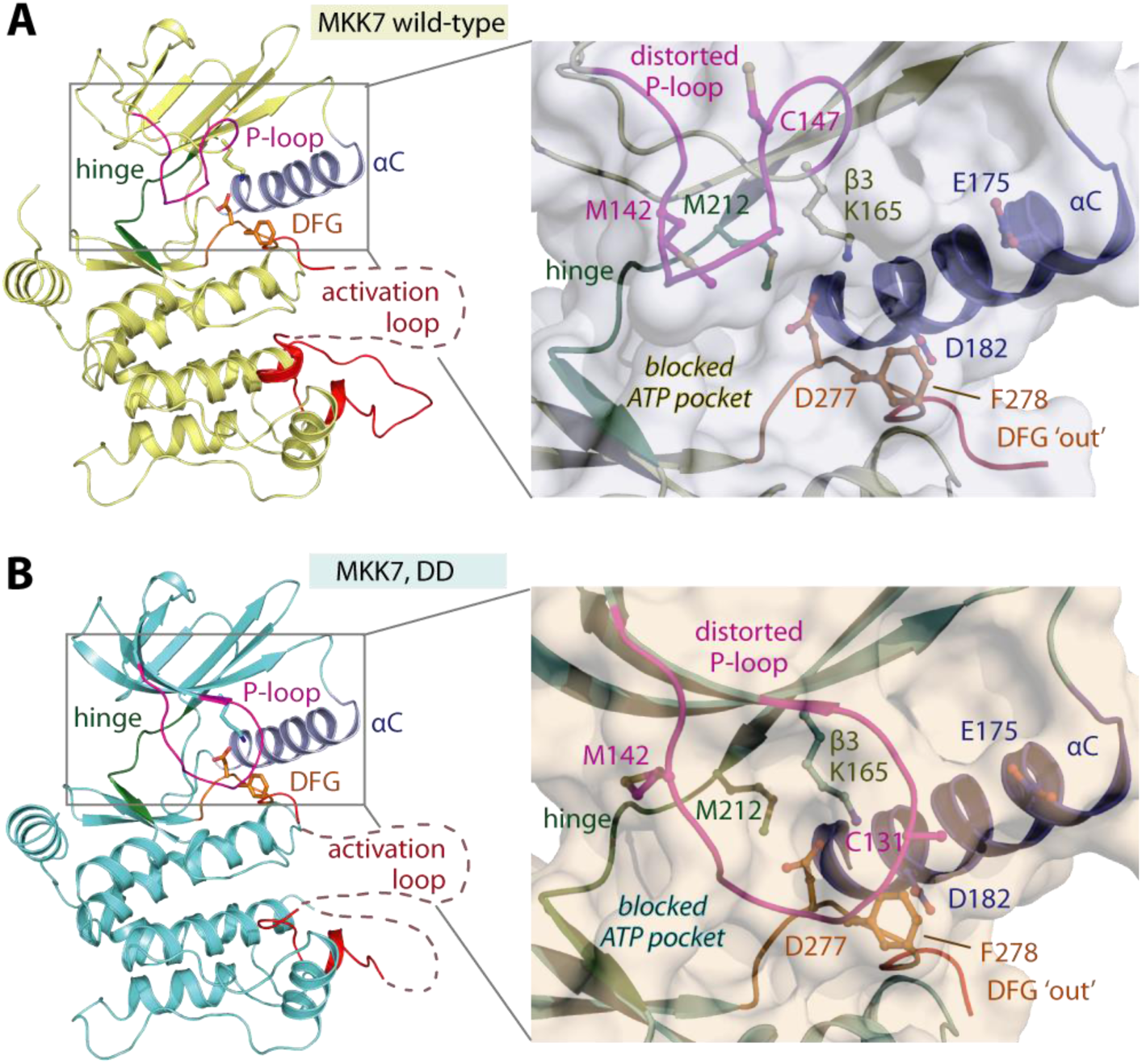
Inactive, auto-inhibitory conformations of the kinase domain of MKK7. Crystal structures of wild-type ΔN115-MKK7 (**A**; MKK7 wild-type) and the corresponding S287D/T291D double mutant (**B**; MKK7, DD) reveal an inactive, catalytically-incompetent state, suggested by an “out” conformation of the DFG motif, the lack of VIAK/αC salt bridge, and a distorted P-loop that occludes the ATP pocket as shown by surface-filled representation in the right panels.

### The role of the N-terminal regulatory helix for an active state of MKK7

High flexibility of the inactive MKK7 kinase domain prompted us to investigate the molecular mechanism for its activation, a long-standing question not only relevant for MKK7 but for the entire MAP2K subfamily. Since the introduction of two phosphorylation-mimetic mutations was insufficient to induce an active state in the crystal structure, we postulated a potential contribution of other regulatory elements that could trigger kinase activation. Previous studies have demonstrated that MEK1 harbors the N-terminal helix preceding the kinase domain, a regulatory element that controls kinase activity ^*41, 42*^. Consistent with previous prediction ^*43*^, comparative sequence analyses suggested that despite slight variations for example in the αC glutamate position, MAP2K members shared highly similar domain architecture, prompting us to postulate existence of the N-terminal helix in MKK7 with a similar regulatory role to that in the counterpart MEK1/2 (Figure 2A and Supplementary Figure S1)^*41*^.

**Figure 2:**
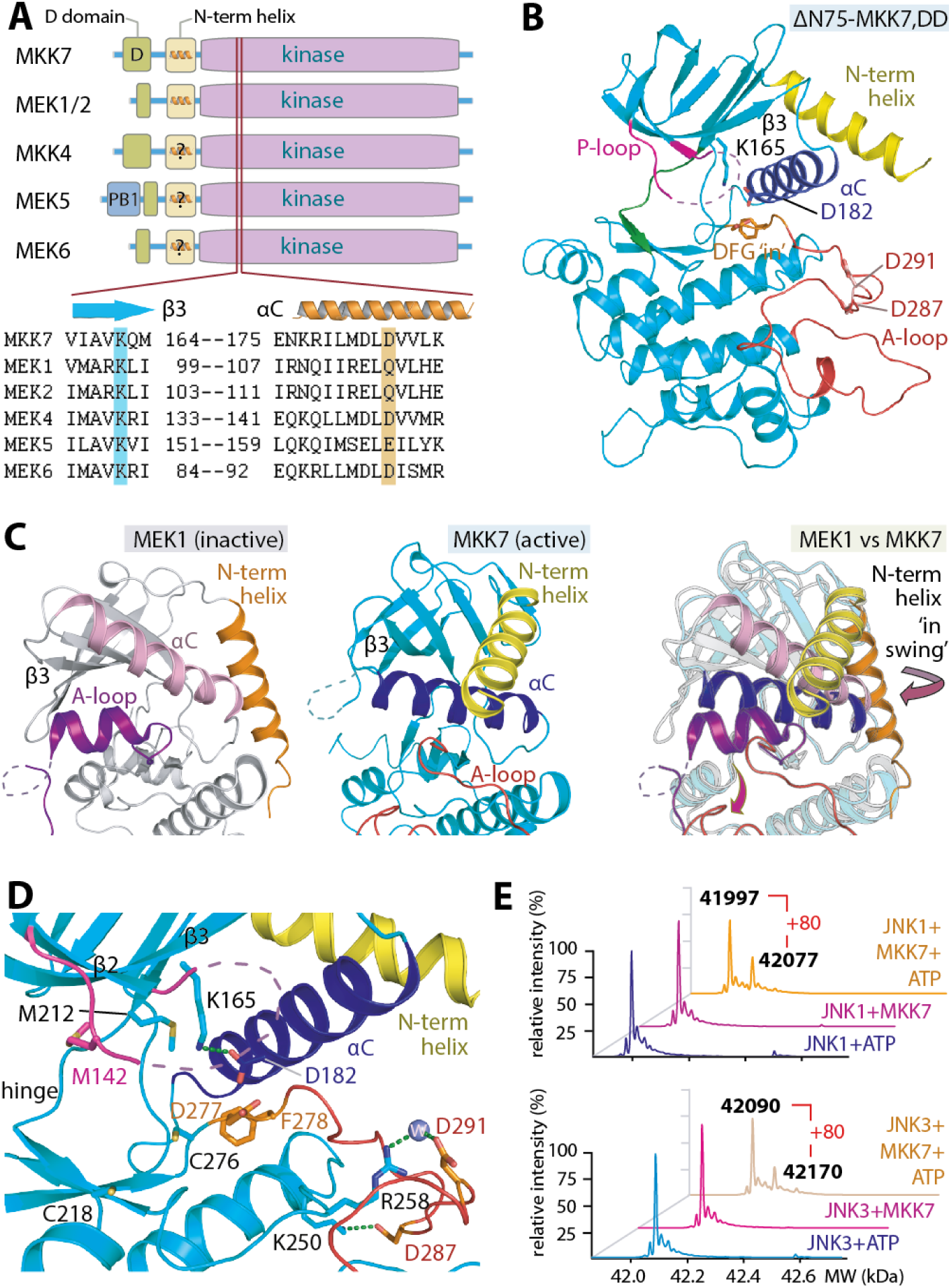
Active conformation of MKK7 induced by the N-terminal helix. **A**) Domain architecture comparison among the MAP2K subfamily members reveals similarly an extensive linker, which is predicted based on MEK1 to form an N-terminal regulatory helix (N-term helix), in between the C-terminus kinase domain and the N-terminus MAPK docking domain. Sequence comparison demonstrates slight variation, either glutamate of aspartate, at the catalytic αC residue. **B**) Crystal structure of ΔN75-MKK7 that harbors two phosphorylation-mimetic S287D/T291D mutations (ΔN60-MKK7, DD) demonstrates the proximity of the N-term helix to the αC. **C**) Structural comparison between inactive MEK1 (pdb id: 3eqc) and active ΔN75-MKK7, DD demonstrates distinct positions and trajectories of their N-term helix. **D**) Closed up details of the catalytic features in MKK7 potentially induced by the N-terminal helix and the S287D and T291D mutations, including the β3 Lys165-αC Asp182 salt bridge, an ‘in’ DFG as well as an ordering of the activation loop (A-loop). **E**) An increase in 80 Dalton mass indicates phosphorylation of JNK1 and JNK3 by catalytically-active S287D/T291D full-length MKK7.

To investigate the presence of the N-terminal regulatory helix in MKK7 as well as its role in kinase activation, we attempted to crystallize three different MKK7 with extended N-terminal regions, full-length, ΔN20 and ΔN75, all of which had S287 and T291 mutated to aspartic acid, a mutation known to mimic phosphorylation^*44, 45*^. Only ΔN75, S287D/T291D-MKK7(ΔN75-MKK7, DD) led to successful crystallization, from which the structure gratifyingly revealed that the kinase adopted an active conformation with the extended N-terminal region displaying the expected helical structure (Figure 2B). In comparison, the position and trajectory of the N-terminal regulatory helix in the active MKK7 that packed against the αC in the front of the kinase juxtaposed that of the counterpart helix in the inactive MEK1, which is located on the back of the hinge (Figure 2C). Although we observe no significant movement of the αC when compared with the inactive apo state, the N-terminal helix did trigger several structural rearrangements (Figure 2D). These included an ‘inward’ swing of the αC Asp182 towards the ATP binding pocket, which enables formation of the canonical salt bridge with Lys175, as well as creates a steric effect on Asp277 resulting in structural rearrangement of the DFG motif into an ‘in’ active state (Supplementary Figure S2). Another indication that this structure represented an active conformation was a complete structuring of the activation segment, assisted by direct and water-mediated interactions between the two phosphorylation mimetic Asp287 and Asp191 to Lys250 and the HRD Arg258. However, the P-loop remained disordered, which was likely due to the absence of the ATP co-factor. The active state of the N-terminally-extended MKK7 observed in the crystal structure was in agreement with catalytic competency of the S287D/T291D full-length kinase, which contains the MAPK docking domain (D-domain), for JNK1 and JNK3 phosphorylation (Figure 2E).

### Inhibitor screening identified diverse hits for MKK7

Considering the flexible nature of MKK7 kinase domain in its inactive state which exhibits restricted access to its ATP binding pocket, we examined whether the kinase could accommodate diverse types of inhibitors. We screened our in-house library of 360 compounds against MKK7 using a melting thermal shift (ΔTm) assay^*46*^. These efforts identified nine inhibitors that resulted in ΔTm of more than 5 °C indicating potent binding (Figure 3A and 3B and Supplementary table 2). The most potent hits comprised a wide variety of compounds, such as type-I inhibitors (ibrutinib, OTSSP167 and CPT1-70-1; ΔTm ∼ 11-13 °C) and several trifluoromethyl-benzene-based type-II inhibitors (Figure 3B and 3C). The highest thermal shift of 24.7 °C was observed by a type-II compound, TL10-105. Despite much lower ΔTm, several type-I inhibitors such as K00007 and ASC69 were also interesting hits due to their distinct hinge binding motifs. Overall, the screening data suggested that highly flexible ATP binding pocket of MKK7 can be targeted by diverse inhibitor scaffolds and binding modes.

**Figure 3:**
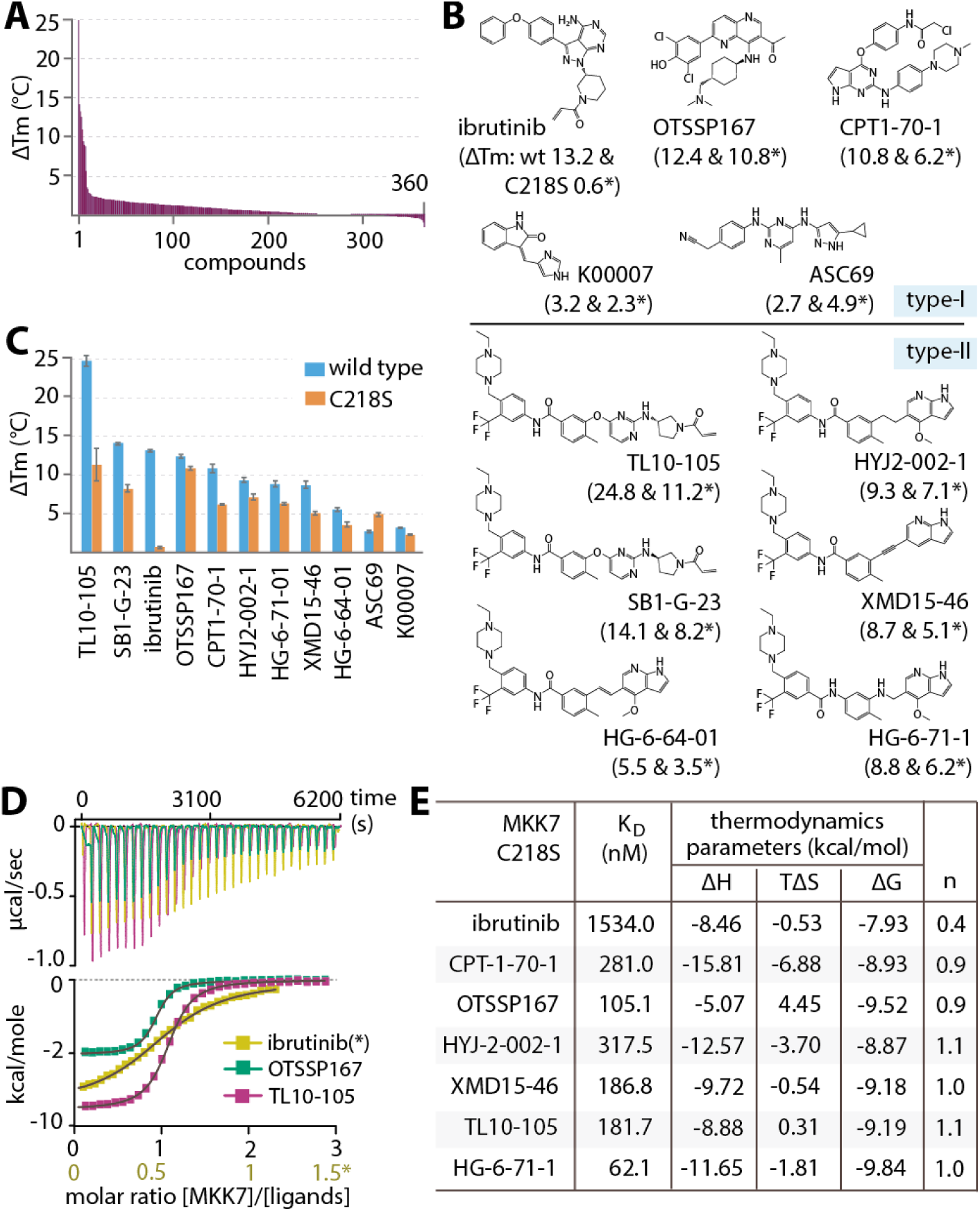
Inhibitor screening for MKK7. **A**) Thermal shift profile of wild-type MKK7 kinase domain against 360 compounds (see Supplementary table 2 for full results). **B**) Chemical structures of the top nine hits as well as two additional weak binder K00007 and ASC69. Thermal shifts (ΔTm) of the inhibitors measured for wild-type kinase and C218S mutant, indicated as * for the latter, are shown in brackets. **C**) Plots of ΔTm values of the selected compounds for the wild type kinase and C218S mutant. **D**) Examples of ITC data for the binding of selected inhibitors in the C218S mutant. Shown are the raw isotherms (top) and the integrated heat of binding with the independent binding fitting (bottom). Note the different scale on the molar ratio for ibrutinib as indicated by *. Due to the similar binding affinity of ibrutinib to the ATP and allosteric site, the binding isotherm has been fitted to a single site model. The derived affinity data should therefore be considered apparent binding data. **E**) Summary of ITC binding constants and thermodynamic parameters for the interactions between the inhibitors and the C218S mutant. Note the stoichiometry of 0.4 for ibrutinib was likely due to its additional interaction with an allosteric binding site (see text for details).

Further analyses of the chemical structures suggested that several compounds including ibrutinib, CPT1-70-1, TL10-105 and SB1-G-23 may bind to the kinase domain by an irreversible binding mechanism. To provide evidence for covalent interactions, we examined C218S MKK7 mutant and showed that, although binding of non-covalent inhibitors remained the same, ΔTm for covalent inhibitors decreased significantly (Figure 3C). Comparable thermal shifts for almost all inhibitors for the mutant suggested similar reversible binding affinities, which was confirmed by isothermal calorimetry (ITC) K_D_ values of 62-318 nM (Figure 3D and F). However, ibrutinib displayed unusual behavior, with ΔT_m_ of 0.6 °C and an ITC K_D_ of ∼1.5 μM for the C218S mutant, suggesting that interaction with the wild type kinase is largely driven by irreversible binding with little contribution of its ATP-mimetic moiety.

### Potencies of the identified inhibitors in enzyme kinetic and cell-based assays

We next performed an activity-based assay to assess inhibitory potencies of nine selected inhibitors for S287D/T291D full-length MKK7. Unexpectedly, when the compounds and ATP were added at the same time only CPT1-70-1 and OTSSP167 exhibited IC_50_s in sub-micromolar range (0.69 and 0.26 μM, respectively) (Figure 4A and 4B). Given that pre-incubation time may affect inhibitor potency^*33*^, we conducted the assays with a 30-minute pre-incubation period. This resulted in significantly lower IC_50_ values for all compounds, except K00007 (Figure 4A and 4B). Different degrees of potency changes corresponded to different modes of inhibition, with reversible compounds, such as type-I OTSSP167 and ASC69 and type-II HG-6-71-01, displaying a modest decrease (2.7-4.3-fold), and covalent inhibitors, including type-I ibrutinib and CPT1-70-1 and type-II TL10-105 and SB1-G-23, exhibiting a significantly larger IC_50_ shift (8-15 fold). Overall, ibrutinib, CPT1-70-1, OTSSP167 and TL10-105 were the most potent inhibitors with their apparent IC_50_s in the 0.06-0.16 μM range.

**Figure 4:**
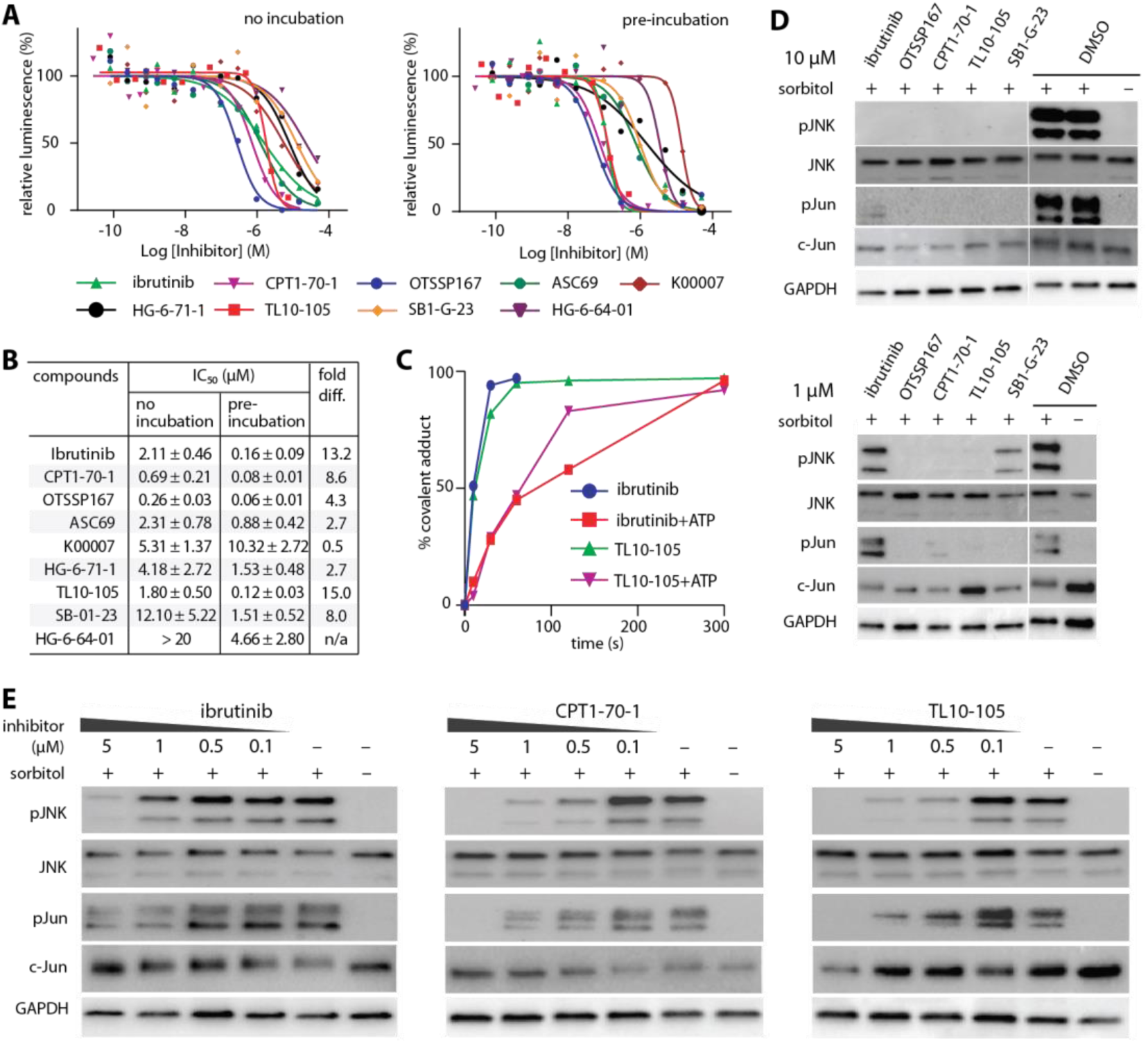
*In vitro* potencies of selected inhibitors for MKK7. **A**) Dose-response plots for inhibition of S287D/T291D full-length MKK7 with different inhibitors from two different experimental settings; *left* for no inhibitor pre-incubation, and *right* for 30-minute pre-incubation. **B**) Summary of average IC_50_ values for both assays calculated from triplicates. The fold differences in the IC_50_ values (fold diff.) between two experiment settings were indicated. **C**) Plots of the percentage of adduct between S287D/T291D full-length MKK7 and type-I ibrutinib or type-II TL10-105 measured by mass spectrometry over the measured time scale. The reactions were performed in two different settings, with or without the supplementation of 400 μM ATP as indicated. **D)** Effects of MKK7 inhibitors on downregulation of phosphorylated JNK and c-Jun levels in sorbitol-induced THP-1 cells. **E)** Three MKK7 covalent inhibitor type-I ibrutinib, type-I CPT1-70-1 and type-II TL10-105 exhibit dose-response inhibition on JNK signaling in sorbitol-induced THP-1 cells, albeit with different potencies, demonstrated by the decrease of phosphorylated JNK and c-Jun.

The results from the enzymatic assays together with the crystal structures suggested that ATP competition as well as energy barriers between MKK7 conformations could present the determinant factors for inhibitor binding. To assess this, we used liquid-chromatography-coupled mass spectrometry (LC-MS) to analyze the kinetics of adduct formation between MKK7 and two covalent inhibitors, type-I ibrutinib and type-II TL10-105, which required different structural rearrangement due to their different modes of binding. We observed that covalent adduct formation was completed faster in the absence of ATP in both cases, while in the presence of ATP TL10-105 slightly out-performed ibrutinib (Figure 4C). These results suggest that despite high plasticity of MKK7, which allows for diverse inhibitor binding, fine-tuning of the ATP-mimetic moieties might present a key for achieving high potencies, especially for irreversible inhibitors.

We next assessed the inhibitory potencies of five selected inhibitors on JNK pathway in sorbitol-treated THP-1 cells. While all inhibitors showed complete inhibition of JNK and c-Jun phosphorylation at 10 µM, some phosphorylation was observed at 1 µM concentration with ibrutinib and SB1-G-23 demonstrating weaker inhibitory potencies compared to OTSSP167, CPT1-70-1 and TL10-105 (Figure 4D). We then chose to further profile the cellular potencies of three covalent inhibitors: the type-I inhibitor ibrutinib and CPT1-70-1 as well as the type-II inhibitorTL10-105 in dose response titrations. Although all three compounds exhibited inhibition of JNK and c-Jun phosphorylation in a dose-dependent manner, ibrutinib activity was significantly weaker compared to CPT1-70-1 and TL10-105, which ablated the level of phosphorylated JNKs and c-Jun significantly at nanomolar inhibitor concentration (Figure 4E). Nevertheless, the observed inhibitory profiles could also be influenced by off-targets as seen from kinome wide selectivity profiles for these inhibitors (see supplementary table 3).

### Different modes of inhibitor binding in MKK7

To provide structural insights into inhibitor binding, we determined the co-crystal structures of MKK7 with the identified hit compounds, including ibrutinib, CPT1-70-1, ASC69 and K00007 (Figure 5A-D). The electron density unveiled a clear covalent bond between ibrutinib and CPT1-70-1 with Cys218, which was observed to be oxidized in the non-covalent complex with ASC69 and K00007 (Supplementary figure S3). Despite exerting similarly type-I binding mode, detailed contacts with the hinge varied among the inhibitors due to their different chemical moieties. Two hydrogen bonds were observed for ibrutinib 4-aminopyrazolopyrimidine and K00007 oxindole, while CPT1-70-1 and ASC69 formed three interactions through their 2-aminopyrolopyrimidine and aminopyrazole scaffolds, respectively. Structural comparison revealed that the binding modes of ibrutinib and K00007 were reminiscent to that of their related analogue 4-amino-pyrazolopyrimidine-based^*33, 40*^ and indazole-based^*39*^ MKK7 inhibitors, respectively (Supplementary figure S4).

**Figure 5:**
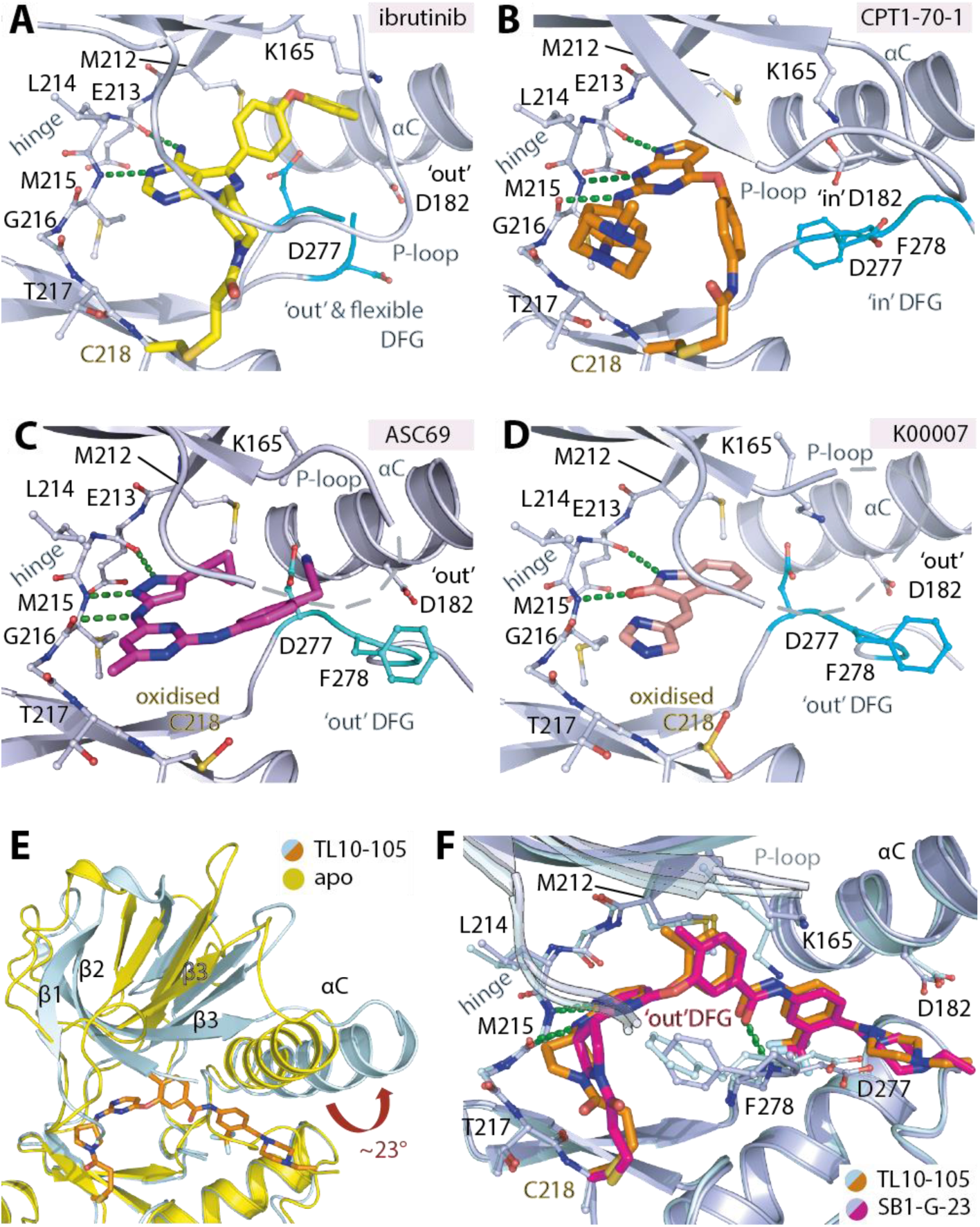
Crystal structures of the complexes between MKK7 and inhibitors. Detailed interactions of ibrutinib (**A**), CPT1-70-1 (**B**), ASC69 (**C**) and K00007 (**D**) within MKK7. All inhibitors are shown in stick representation with green, dashed line indicating hydrogen bonds. **E**) Superimposition of the MKK7-TL10-105 complex and the apo structure reveals structural alterations of the N-terminal lobe, in essence the αC, necessary for the type-II inhibitor binding. **F**) Structural comparison between the TL10-105- and SB1-G-23-MKK7 complexes. Both inhibitors form covalent adduct with Cys218 and share highly similar binding modes, with an exception noted for different configurations of the inhibitor pyrrolidine moieties and the DFG Phe278.

In addition to different details of the observed interaction, comparative structural analyses demonstrated structural alterations within the kinase upon binding to diverse type-I compounds. CPT1-70-1 induced an ‘in’ conformation of the DFG motif and Asp182, while the other three type-I inhibitors bound with an ‘out’ DFG conformation. Of particular note was the binding mode of ibrutinib where the phenoxyphenyl moiety of the inhibitor protruded into a hydrophobic back pocket in proximity to the αC, thus destabilizing the DFG motif and resulting in two distinct conformations. Another prominent structural difference was the position of the P-loop, which assumed a range of different conformations: disordered in the ASC69 and K00007 complexes; distorted in the ibrutinib complex; and structured in the CPT1-70-1 complex. These inhibitor-bound structures further support the high plasticity of the kinase ATP binding pocket.

We also determined the structures of ΔN115-MKK7 bound to TL10-105 and its R-enantiomer SB1-G-23, two type-II inhibitors. Both inhibitors formed a covalent adduct with Cys218 in a similar manner to that observed for type-I ibrutinib and CPT1-70-1 (Figure 5E-F). In comparison to the apo structure, accommodation of these type-II compounds required significant structural alteration, exemplified by a ∼23° ‘out’ swing of the αC to create the hydrophobic pocket necessary for the trifluoromethyl-benzene moieties (Figure 5E). Although TL10-105 and SB1-G-23 displayed a highly similar binding mode, we observed different configurations of the pyrrolidine ring, which may explain their different potencies (Figure 5F).

### Ibrutinib exhibits additional binding at an allosteric site

Unexpectedly, our analysis of ibrutinib-MKK7 structure revealed not only a covalently bound inhibitor in the active site, but some extra electron density within the N-lobe. We were able to model an additional molecule of ibrutinib into that electron density, suggesting a 2:1 ibrutinib:MKK7 binding stoichiometry (Figure 6 and Supplementary Figure S3), and in agreement with the ITC measurements (Figure 3E). The presence of the inhibitor in this allosteric site was rather unexpected, and was unlikely to be influenced by crystal packing since this region of the kinase was highly solvent exposed. In addition, a second crystal structure where ibrutinib bound only to the allosteric site in MKK7 with an empty ATP binding pocket suggested that such allosteric interaction was independent of and not induced by the presence of ligands in the ATP binding site. Comparison with the apo structure indicated that this allosteric pocket was induced by ibrutinib binding. Significant structural alterations for the pocket formation included an ‘out’ swing of Phe202, an outward movement of the helical turn preceding the β1 that harbors Ile133 as well as a small alteration of Trp151 side chain (Figure 6A). Nevertheless, direct contact between ibrutinib and the kinase at this allosteric site was limited to only a potential π-stacking with the β5 Phe209 (Figure 6B). To assess if the allosteric binding of the inhibitor would occur in solution, we performed ITC experiment using the pre-formed ibrutinib-covalent-adduct MKK7 of which the ATP binding pocket was blocked. Indeed, the change of titration heat demonstrated ibrutinib binding to this MKK7 adduct with the affinities measured to be ∼12.6 μM (Figure 6C). Taken together, we reveal the existence of previously unobserved allosteric ligand binding site in MKK7.

**Figure 6:**
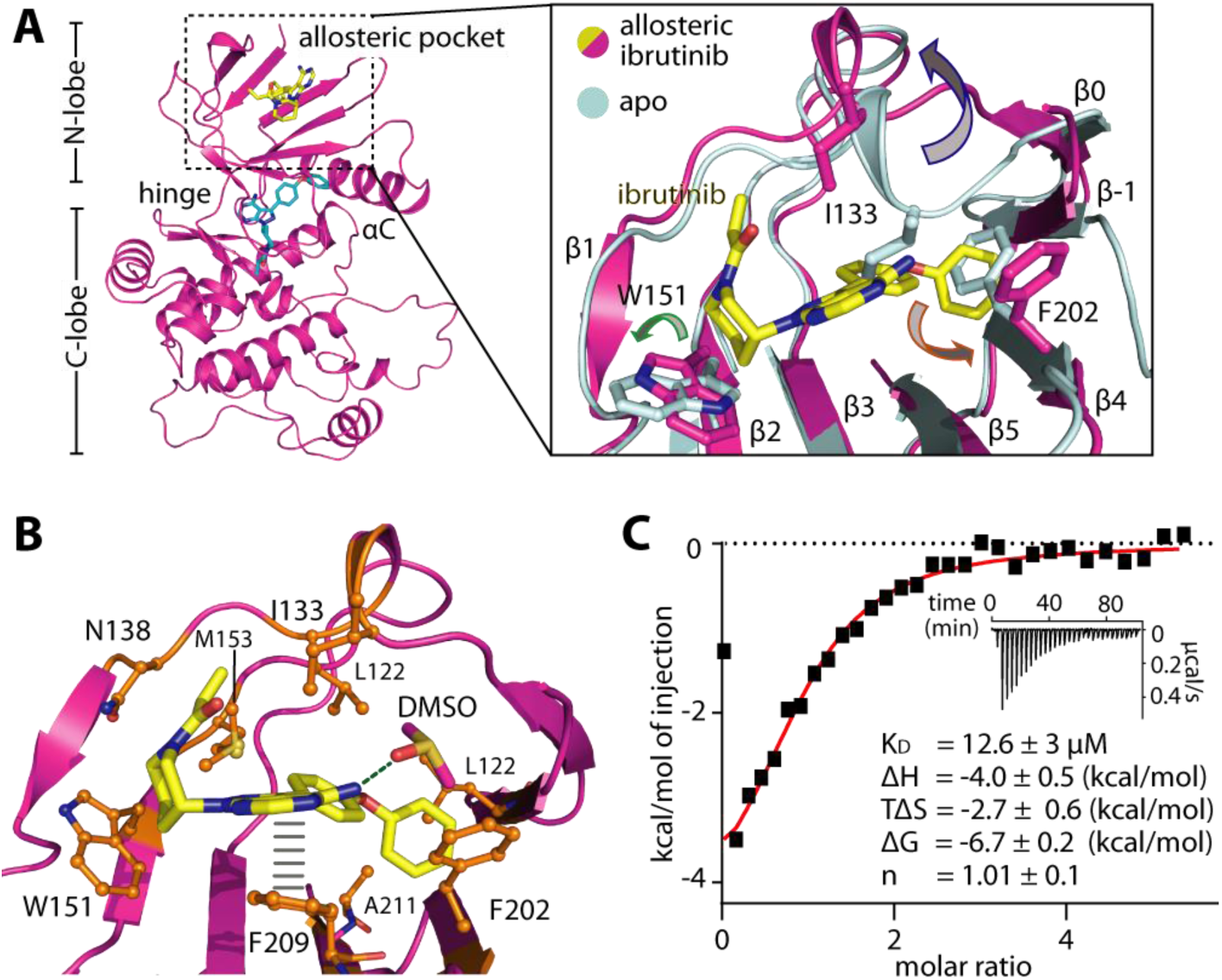
Allosteric binding of ibrutinib. **A**) Crystal structure demonstrates the binding of the second ibrutinib (yellow stick) at the allosteric pocket located on top of the N-terminal lobe of MKK7. Inset shows structural superimposition with the apo-structure, revealing the required structural rearrangements for the pocket formation (indicated by arrows; see text for details). **B**) The interactions between ibrutinib and the kinase at the allosteric site. Grey bars indicate π-stacking between Phe209 and the bound ibrutinib, which makes also a hydrogen bond to a DMSO molecule bound in vicinity. **C**) ITC binding data for the allosteric interaction between ibrutinib and the pre-formed ibrutinib-adduct MKK7. Shown are the normalized heat of binding with the single-site fitting (red line), and the raw titration isotherms (inset). The thermodynamic parameters and equilibrium binding constants (K_D_) are the average from triplicates.

## Discussion

Second-tier kinase MKK7 is a key mediator of the JNK signaling response, and represents a potential drug target for selective inhibition of JNK signaling. Here, we present a comprehensive biochemical and structural characterization of a diverse set of MKK7 inhibitors that includes both non-covalent and covalent binders, as well as both type I and type II inhibitors. Overall, despite the role of auto-inhibition in MKK7 regulation, our study demonstrates that this kinase possesses a plastic ATP-binding site that can accommodate a range of scaffolds. Moreover, accessibility of a reactive cysteine suggests additional opportunities for targeting this kinase. In addition to a range of type-I previously reported compounds and scaffolds that include 5Z-7-oxozeaenol^*36*^, indazole-based derivatives^*39*^, 4-amino-pyrazolopyrimidine-based inhibitors^*33, 40*^ and aminopyrimidine SM1-71^*35*^, we now expand the MKK7 inhibitor pool to include covalent type-II inhibitors. More broadly, TL10-105 and SB1-G-23, two compounds we identified and characterized here represent, to our knowledge, unique examples of type-II covalent kinase inhibitors. Another unexpected observation that we make in this study stems from the analysis of ibrutinib binding to MKK7. As seen in the crystal structure of MKK7-ibrutinib complex, and further corroborated by our ITC measurement, ibrutinib bound to MKK7 in 2:1 stoichiometry, with the second molecule of ibrutinib bound within the newly discovered allosteric pocket at the N-terminal lobe of MKK7. Although the biological function of this allosteric pocket remains to be determined, previous studies have proposed that this N-terminal lobe region is involved in mediating protein-protein interactions in the context of MKK7^*6*^, as well as other binding partners, including GADD45β^*47, 48*^. While the molecular details of the GADD45β binding site on the upper kinase still needs to be described, targeting the MKK7-GADD45β interaction, for example with the peptide-mimetic inhibitor DTP3, has been proposed as an alternative chemotherapeutic strategy for treatment of some cancer types with elevated GADD45β levels such as multiple myelomas^*26*^. Also for the tyrosine kinase ABL, a second binding site of the FDA approved inhibitor gleevec that comprises the myristate binding pocket has been described ^*49*^, which resulted in a new generation of kinases inhibitors that are now tested clinically ^*50*^. Collectively, the range of structural information described in this work together with the discovery of MKK7 allosteric pocket, and the initial description of type-II covalent kinase inhibitors offers an important resource for future medicinal chemistry efforts. In addition, the two most potent type-I CPT1-70-1 and type-II TL10-105 with significant cellular inhibitory effects on JNK signaling pathway may offer an alternative chemical starting point for MKK7 inhibitor development in addition to the previous indazole-and aminopyrazolopyrimidine -based inhibitors.

Mechanistically, our crystal structures provide not only detailed molecular models into how MKK7 engages different scaffolds, but offer insights into MKK7 auto-inhibition and activation as well. Previous studies have reported diverse inactive configurations as auto-inhibitory mechanisms for MKK7, including various distorted conformations of the P-loop^*6, 7*^ as well as the hinge Leu214 in an unprecedented ‘in’ conformation^*33*^. Here, we demonstrate that MKK7 can assume an inactive ‘out’ DFG conformation, thus displaying an additional mechanism for occlusion of the ATP site and auto-inhibition^*7*^.

Molecular mechanisms underlying kinase activation has been a long standing question for MAP2K subfamily. Here, our structure of active MKK7 provides an exclusive insight into molecular process of kinase activation, revealing an essential role of the N-terminal regulatory helix. This regulatory helix has previously been described in MEK1^*41*^, and is likely to be present in other MAP2Ks^*43*^. We speculate that activation segment phosphorylation together with correct positioning of the regulatory helix in proximity of the αC triggers simultaneous structural rearrangements, leading to catalytic salt bridge formation as well as formation of the substrate binding site. In this scenario, the effects of the N-terminal regulatory helix would be allosteric. Our structural insights were consistent with the recent molecular dynamic studies, demonstrating that the regulatory helix which is located further away in an inactive state could undergo conformational changes and form complex network interactions with the αC and activation segment necessary for stabilization and activation of MEK1^*41, 42*^. Although it remains to be formally confirmed, we expect that similar mechanism based on the allosteric effect of the N-terminal regulatory helix might be shared across the MAP2K subfamily.

## Significance

Among the MAP2K subfamily members, MKK7 a poorly studied kinases despite being a key regulator of the JNK stress signaling pathway presents opportunities for translational exploitation. Here we present a comprehensive set of crystal structures unveiling a number of unique structural features, which significantly expand our mechanistic understanding of this kinase, as well as the MAP2K family in general. An intriguing finding was the allosteric role of the N-terminal regulatory helix which is conserved in all MAP2K kinases. Our structural data showed that this helix triggers transformation into the active state providing a first model of the activation mechanism in MAP2Ks and the active conformation of these key drug targets. We identified a diversity of potent inhibitors that significantly expand the number of chemical scaffolds that can be utilized targeting MKK7. Importantly, diverse co-crystal structures proposed new targeting strategies and provided insight into the conformational plasticity of the MKK7 catalytic domain. For instance, we present for the first time high resolution structures of irreversible type-II inhibitors offering a template for future exploitation of this binding mode. Additionally, we discovered an unprecedented allosteric pocket located in the N-terminal lobe that can be targeted by the approved drug ibrutinib. Collectively, our structural and functional data revealed mechanistic insights into the activation mechanism of MAP2Ks, and provide new targeting strategies for this important kinase.

## Acknowledgement

M.S., A.C. and S.K. are grateful for support by the SGC, a registered charity (number 1097737) that receives funds from AbbVie, Bayer Pharma AG, Boehringer Ingelheim, Canada Foundation for Innovation, Eshelman Institute for Innovation, Genome Canada, Innovative Medicines Initiative (EU/EFPIA), Janssen, Merck KGaA, Germany, MSD, Novartis Pharma AG, Ontario Ministry of Economic Development and Innovation, Pfizer, São Paulo Research Foundation-FAPESP, Takeda, Wellcome [106169/ZZ14/Z]. We are grateful for support by the Collaborative Sonderforschungsbereich 1177 Autophagy (SFB1177, to M.S., A.C. and S.K.) at Frankfurt University, as well as the German Cancer Consortium (DKTK to SK). The authors thank staffs at Diamond Light Source for their support during crystallographic X-ray diffraction data collection. We thank Dr. Milka Kostic (Twitter: @MilkaKostic) for the assistance in critical reading and editing the manuscript.

## Author Contributions

A.C. designed the research. A.C., S.K. and N.S.G. supervised the research. A.C. performed structural study. M.S. performed biochemical and biophysical study. L.T., J.W. and Y.L. contributed and synthesized compounds. A.C. drafted the manuscript, which has been edited and approved by all authors.

## Method details

### Compounds

Sources and chemical syntheses of inhibitors are described in Supplementary information.

### Protein expression and purification

All MKK7 constructs, including i) wild-type ΔN115-MKK7 (aa 116-421), ii) C218S ΔN115-MKK7, iii) S287D/T291D ΔN115-MKK7, iv) S287D/T291D ΔN75-MKK7 (aa 76-421), and v) S287D/T291D full-length MKK7 (1-421), were sub-cloned into pNIC28-Bsa4 and were expressed as N-terminal His_6_-tagged fusions in *E. coli*. Recombinant proteins were purified using Ni^2+^-affinity chromatography, and the histidine tags were subsequently cleaved using TEV protease. The cleaved proteins were passed through Ni^2+^ beads, and further purified by size exclusion chromatography. MKK7 proteins were stored in 20 mM HEPES, pH 7.5, 150-200 mM NaCl and 0.5 mM TCEP.

### Crystallization and structure determination

Crystallization was performed using MKK7 proteins at ∼8-11 mg/ml and sitting drop vapor diffusion method at 4 or 20 °C with the conditions described in Supplementary table 1. Successful structural determination of different complexes was obtained using different MKK7 proteins as outlined in Supplementary table 1. For type-I inhibitor complexes, soaking of the inhibitors into wild-type ΔN115-MKK7 apo crystals was performed. For type-II inhibitor complexes, the S287D/T291D ΔN115-MKK7 protein was incubated with the inhibitors prior to co-crystallization. All crystals were cryo-protected with mother liquor supplemented with 20-25% ethylene glycol before flash cooled in liquid nitrogen. Diffraction data collected at Diamond Light Source were processed with XDS ^*51*^ or iMosflm ^*52*^, and scaled using Scala from CCP4 suite^*53*^. Initial structure solution was obtained by molecular replacement using Phaser^*54*^ and published MKK7 coordinates (pdb id: 2dyl). Manual model rebuilding was performed in COOT^*55*^, alternated with structure refinement in REFMAC ^*56*^. Geometry of the final structures was verified using Molprobity ^*57*^. Data collection and refinement statistics are summarized in Supplementary table 1.

### Thermal shift assays

Recombinant MKK7 at 2 μM in 10 mM HEPES, pH 7.5 and 500 mM NaCl was mixed with 10 μM inhibitors, and the reaction was incubated briefly at room temperature for ∼10 mins. SyPRO orange dye (Invitrogen) was subsequently added, and the fluorescence signals corresponding to temperature-dependent protein unfolding were measured using a Real-Time PCR Mx3005p machine (Stratagene). The melting temperature calculation was performed as described previously^*46*^.

### Isothermal calorimetry

All isothermal calorimetry (ITC) experiments were performed on NanoITC (TA instrument) at 25 °C. For binding studies of ATP-competitive inhibitors, C218S ΔN115-MKK7 at ∼0.1 mM in 20 mM HEPES, pH 7.5, 150 mM NaCl, 0.5 mM TCEP and 5% glycerol was titrated into the reaction cell containing the inhibitors at ∼0.01 mM. For ibrutinib allosteric binding study, the covalent adduct of wild-type ΔN115-MKK7 with ibrutinib was firstly prepared by incubating the protein with two-fold molar excess of Ibrutinib prior to purification by size exclusion chromatography. Completion of the covalent formation was verified by mass spectrometry, and the resulting MKK7 kinase domain covalently pre-bound to ibrutinib at ∼0.32 mM was titrated into ibrutinib at 0.02 mM. The integrated heat of titration was calculated and fitted to a single, independent binding model using the software provided by the manufacture. The thermodynamic parameters (ΔH and TΔS), equilibrium association and dissociation constants (Ka and K_D_), and stoichiometry (n) were calculated.

### Activity-based inhibition assay

Activity of S287D/T291D full-length MKK7 was measured using ADP-Glo assay (Promega). JNK3 covalently pre-bound to JNK-specific covalent inhibitor^*30*^ for blocking its ATP binding site was used as a substrate. MKK7 at 25 nM and JNK3 at 30 µM were mixed in the reaction buffer containing 40 mM tris, pH 7.5, 20 mM MgCl2, 60 mM NaCl and 1% Glycerol. Inhibitors were then added into the mixture using an ECHO acoustic dispenser. Two sets of experiment were conducted. For the ‘no incubation’ set, ATP at final 0.1 mM concentration was added immediately, while for the ‘pre-incubation’ experiment the MKK7-JNK3-inhibitor mixtures were pre-incubated at 4 °C for 30 mins prior to ATP addition. The reaction was stopped at 10 mins after ATP addition, and the luminescence signals were analyzed according to the manufacture’s protocols using BMG PheraStar. IC_50_ values of inhibitors were calculated using Prism software.

### Mass spectrometry analyses for covalent adduct formation

S287D/T291D full-length MKK7 at 10 µM was incubated with compounds at two-fold molar excess in buffer containing 20 mM HEPES pH 7.5, 200 mM NaCl, 0.5 mM TCEP and 5% Glycerol. For ATP competition experiment, 20 mM MgCl_2_ and 0.4 mM ATP were added. The reactions were stopped at different time points by mixing with a solution of 1% acetic acid and 200 mM DTT. The samples were purified on C18 stage tips using protocol described previously^*58*^ prior to intact mass analyses using LC-TOF mass spectrometer (Agilent).

### Western blot analyses of inhibitor cellular potencies

THP-1 cells (ATCC, TIB-202) were cultured at 37°C with 5% CO_2_ in RPMI 1640 (Invitrogen) supplemented with 10% heat inactivated FBS (Invitrogen) and Gibco(tm) penicillin-streptavidin antibiotic (invitrogen). Cells were seeded at 600,000 cells/ml and treated for 2 hours with inhibitors or DMSO followed by an addition of 0.6 M sorbitol (Sigma-Aldrich) for 20 minutes. Cells were harvested by centrifugation at 100xg for 3 minutes, and washed with ice cold PBS and subsequently lysed in buffer containing 50 mM Tris, pH 8.0, 150 mM NaCl, 1% NP40 (Roche) and a phosphatase inhibitor mixture containing 20 mM NaF, 2 mM Na3VO4, 2 mM β-phosphoglycerol. Samples were analyzed by SDS/PAGE under reducing conditions, followed by Western blotting using phospho-JNK, JNK, phosphor-c-Jun and c-Jun antibodies (Cell Signaling Technology).

## Supporting Information

The supporting information includes figures, tables, crystallographic data and supplementary methods for chemical synthesis.

## Notes

The coordinates and structure factors of all complexes have been deposited to the protein data bank under accession codes 6YFZ, 6YG0, 6YG1, 6YG2, 6YG3, 6YG4, 6YG5, 6YG6, and 6YG7.

## Abbreviations

MKK7 and MAP2K7: mitogen-activated protein kinase kinase 7
JNK: c-Jun N-terminal kinase
MKK4 and MAP2K7: mitogen-activated protein kinase kinase 4
MAPK: mitogen-activated protein kinase
MAP2K: mitogen-activated protein kinase kinase
MEK1: mitogen-activated protein kinase kinase 1
MEK2: mitogen-activated protein kinase kinase 2

## References

[1] Davis, R. J. (2000) Signal transduction by the JNK group of MAP kinases, Cell 103, 239–252.

[2] Kragelj, J., Palencia, A., Nanao, M. H., Maurin, D., Bouvignies, G., Blackledge, M., and Jensen, M. R. (2015) Structure and dynamics of the MKK7-JNK signaling complex, Proc Natl Acad Sci U S A 112, 3409–3414.

[3] Ho, D. T., Bardwell, A. J., Grewal, S., Iverson, C., and Bardwell, L. (2006) Interacting JNK-docking sites in MKK7 promote binding and activation of JNK mitogen-activated protein kinases, J Biol Chem 281, 13169–13179.

[4] Fleming, Y., Armstrong, C. G., Morrice, N., Paterson, A., Goedert, M., and Cohen, P. (2000) Synergistic activation of stress-activated protein kinase 1/c-Jun N-terminal kinase (SAPK1/JNK) isoforms by mitogen-activated protein kinase kinase 4 (MKK4) and MKK7, Biochem J 352 Pt 1, 145–154.

[5] Takekawa, M., Tatebayashi, K., and Saito, H. (2005) Conserved docking site is essential for activation of mammalian MAP kinase kinases by specific MAP kinase kinase kinases, Mol Cell 18, 295–306.

[6] Kinoshita, T., Hashimoto, T., Sogabe, Y., Fukada, H., Matsumoto, T., and Sawa, M. (2017) High-resolution structure discloses the potential for allosteric regulation of mitogen-activated protein kinase kinase 7, Biochem Biophys Res Commun 493, 313–317.

[7] Sogabe, Y., Hashimoto, T., Matsumoto, T., Kirii, Y., Sawa, M., and Kinoshita, T. (2016) A crucial role of Cys218 in configuring an unprecedented auto-inhibition form of MAP2K7, Biochem Biophys Res Commun 473, 476–481.

[8] Han, Z., Boyle, D. L., Chang, L., Bennett, B., Karin, M., Yang, L., Manning, A. M., and Firestein, G. S. (2001) c-Jun N-terminal kinase is required for metalloproteinase expression and joint destruction in inflammatory arthritis, J Clin Invest 108, 73–81.

[9] Roy, P. K., Rashid, F., Bragg, J., and Ibdah, J. A. (2008) Role of the JNK signal transduction pathway in inflammatory bowel disease, World J Gastroenterol 14, 200–202.

[10] Huang, Y. A., Zhou, B., Wernig, M., and Sudhof, T. C. (2017) ApoE2, ApoE3, and ApoE4 Differentially Stimulate APP Transcription and Abeta Secretion, Cell 168, 427–441 e421.

[11] Mazzitelli, S., Xu, P., Ferrer, I., Davis, R. J., and Tournier, C. (2011) The loss of c-Jun N-terminal protein kinase activity prevents the amyloidogenic cleavage of amyloid precursor protein and the formation of amyloid plaques in vivo, J Neurosci 31, 16969–16976.

[12] Wagner, E. F., and Nebreda, A. R. (2009) Signal integration by JNK and p38 MAPK pathways in cancer development, Nat Rev Cancer 9, 537–549.

[13] Cellurale, C., Sabio, G., Kennedy, N. J., Das, M., Barlow, M., Sandy, P., Jacks, T., and Davis, R. J. (2011) Requirement of c-Jun NH(2)-terminal kinase for Ras-initiated tumor formation, Mol Cell Biol 31, 1565–1576.

[14] Das, M., Garlick, D. S., Greiner, D. L., and Davis, R. J. (2011) The role of JNK in the development of hepatocellular carcinoma, Genes Dev 25, 634–645.

[15] Chen, N., Nomura, M., She, Q. B., Ma, W. Y., Bode, A. M., Wang, L., Flavell, R. A., and Dong, Z. (2001) Suppression of skin tumorigenesis in c-Jun NH(2)-terminal kinase-2-deficient mice, Cancer Res 61, 3908–3912.

[16] Sakai, H., Sato, A., Aihara, Y., Ikarashi, Y., Midorikawa, Y., Kracht, M., Nakagama, H., and Okamoto, K. (2014) MKK7 mediates miR-493-dependent suppression of liver metastasis of colon cancer cells, Cancer Sci 105, 425–430.

[17] Lotan, T. L., Lyon, M., Huo, D., Taxy, J. B., Brendler, C., Foster, B. A., Stadler, W., and Rinker-Schaeffer, C. W. (2007) Up-regulation of MKK4, MKK6 and MKK7 during prostate cancer progression: an important role for SAPK signalling in prostatic neoplasia, J Pathol 212, 386–394.

[18] Sui, X., Kong, N., Wang, X., Fang, Y., Hu, X., Xu, Y., Chen, W., Wang, K., Li, D., Jin, W., Lou, F., Zheng, Y., Hu, H., Gong, L., Zhou, X., Pan, H., and Han, W. (2014) JNK confers 5-fluorouracil resistance in p53-deficient and mutant p53-expressing colon cancer cells by inducing survival autophagy, Sci Rep 4, 4694.

[19] Nguyen, T. V., Sleiman, M., Moriarty, T., Herrick, W. G., and Peyton, S. R. (2014) Sorafenib resistance and JNK signaling in carcinoma during extracellular matrix stiffening, Biomaterials 35, 5749–5759.

[20] Hayakawa, J., Depatie, C., Ohmichi, M., and Mercola, D. (2003) The activation of c-Jun NH2-terminal kinase (JNK) by DNA-damaging agents serves to promote drug resistance via activating transcription factor 2 (ATF2)-dependent enhanced DNA repair, J Biol Chem 278, 20582–20592.

[21] Xu, H., Cheng, M., Chi, X., Liu, X., Zhou, J., Lin, T., and Yang, W. (2019) High-throughput screening identifies mixed lineage kinase 3 as a key host regulatory factor in Zika virus infection, J Virol.

[22] He, P., Zhang, B., Liu, D., Bian, X., Li, D., Wang, Y., Sun, G., and Zhou, G. (2016) Hepatitis B Virus X Protein Modulates Apoptosis in NRK-52E Cells and Activates Fas/FasL Through the MLK3-MKK7-JNK3 Signaling Pathway, Cell Physiol Biochem 39, 1433–1443.

[23] Hubner, A., Mulholland, D. J., Standen, C. L., Karasarides, M., Cavanagh-Kyros, J., Barrett, T., Chi, H., Greiner, D. L., Tournier, C., Sawyers, C. L., Flavell, R. A., Wu, H., and Davis, R. J. (2012) JNK and PTEN cooperatively control the development of invasive adenocarcinoma of the prostate, Proc Natl Acad Sci U S A 109, 12046–12051.

[24] Schramek, D., Kotsinas, A., Meixner, A., Wada, T., Elling, U., Pospisilik, J. A., Neely, G. G., Zwick, R. H., Sigl, V., Forni, G., Serrano, M., Gorgoulis, V. G., and Penninger, J. M. (2011) The stress kinase MKK7 couples oncogenic stress to p53 stability and tumor suppression, Nat Genet 43, 212–219.

[25] Chen, P., O’Neal, J. F., Ebelt, N. D., Cantrell, M. A., Mitra, S., Nasrazadani, A., Vandenbroek, T. L., Heasley, L. E., and Van Den Berg, C. L. (2010) Jnk2 effects on tumor development, genetic instability and replicative stress in an oncogene-driven mouse mammary tumor model, PLoS One 5, e10443.

[26] Tornatore, L., Sandomenico, A., Raimondo, D., Low, C., Rocci, A., Tralau-Stewart, C., Capece, D., D’Andrea, D., Bua, M., Boyle, E., van Duin, M., Zoppoli, P., Jaxa-Chamiec, A., Thotakura, A. K., Dyson, J., Walker, B. A., Leonardi, A., Chambery, A., Driessen, C., Sonneveld, P., Morgan, G., Palumbo, A., Tramontano, A., Rahemtulla, A., Ruvo, M., and Franzoso, G. (2014) Cancer-selective targeting of the NF-kappaB survival pathway with GADD45beta/MKK7 inhibitors, Cancer Cell 26, 495–508.

[27] Asaoka, Y., and Nishina, H. (2010) Diverse physiological functions of MKK4 and MKK7 during early embryogenesis, J Biochem 148, 393–401.

[28] Plantevin Krenitsky, V., Nadolny, L., Delgado, M., Ayala, L., Clareen, S. S., Hilgraf, R., Albers, R., Hegde, S., D’Sidocky, N., Sapienza, J., Wright, J., McCarrick, M., Bahmanyar, S., Chamberlain, P., Delker, S. L., Muir, J., Giegel, D., Xu, L., Celeridad, M., Lachowitzer, J., Bennett, B., Moghaddam, M., Khatsenko, O., Katz, J., Fan, R., Bai, A., Tang, Y., Shirley, M. A., Benish, B., Bodine, T., Blease, K., Raymon, H., Cathers, B. E., and Satoh, Y. (2012) Discovery of CC-930, an orally active anti-fibrotic JNK inhibitor, Bioorg Med Chem Lett 22, 1433–1438.

[29] Zhang, T., Inesta-Vaquera, F., Niepel, M., Zhang, J., Ficarro, S. B., Machleidt, T., Xie, T., Marto, J. A., Kim, N., Sim, T., Laughlin, J. D., Park, H., LoGrasso, P. V., Patricelli, M., Nomanbhoy, T. K., Sorger, P. K., Alessi, D. R., and Gray, N. S. (2012) Discovery of potent and selective covalent inhibitors of JNK, Chem Biol 19, 140–154.

[30] Muth, F., El-Gokha, A., Ansideri, F., Eitel, M., Doring, E., Sievers-Engler, A., Lange, A., Boeckler, F. M., Lammerhofer, M., Koch, P., and Laufer, S. A. (2017) Tri- and Tetrasubstituted Pyridinylimidazoles as Covalent Inhibitors of c-Jun N-Terminal Kinase 3, J Med Chem 60, 594–607.

[31] Messoussi, A., Feneyrolles, C., Bros, A., Deroide, A., Dayde-Cazals, B., Cheve, G., Van Hijfte, N., Fauvel, B., Bougrin, K., and Yasri, A. (2014) Recent progress in the design, study, and development of c-Jun N-terminal kinase inhibitors as anticancer agents, Chem Biol 21, 1433–1443.

[32] Wang, J., Chen, L., Ko, C. I., Zhang, L., Puga, A., and Xia, Y. (2012) Distinct signaling properties of mitogen-activated protein kinase kinases 4 (MKK4) and 7 (MKK7) in embryonic stem cell (ESC) differentiation, J Biol Chem 287, 2787–2797.

[33] Wolle, P., Engel, J., Smith, S., Goebel, L., Hennes, E., Lategahn, J., and Rauh, D. (2019) Characterization of Covalent Pyrazolopyrimidine-MKK7 Complexes and a Report on a Unique DFG-in/Leu-in Conformation of Mitogen-Activated Protein Kinase Kinase 7 (MKK7), J Med Chem 62, 5541–5546.

[34] Chaikuad, A., Koch, P., Laufer, S. A., and Knapp, S. (2018) The Cysteinome of Protein Kinases as a Target in Drug Development, Angew Chem Int Ed Engl 57, 4372–4385.

[35] Rao, S., Gurbani, D., Du, G., Everley, R. A., Browne, C. M., Chaikuad, A., Tan, L., Schroder, M., Gondi, S., Ficarro, S. B., Sim, T., Kim, N. D., Berberich, M. J., Knapp, S., Marto, J. A., Westover, K. D., Sorger, P. K., and Gray, N. S. (2019) Leveraging Compound Promiscuity to Identify Targetable Cysteines within the Kinome, Cell Chem Biol 26, 818–829 e819.

[36] Sogabe, Y., Matsumoto, T., Hashimoto, T., Kirii, Y., Sawa, M., and Kinoshita, T. (2015) 5Z-7-Oxozeaenol covalently binds to MAP2K7 at Cys218 in an unprecedented manner, Bioorg Med Chem Lett 25, 593–596.

[37] Bradshaw, J. M., McFarland, J. M., Paavilainen, V. O., Bisconte, A., Tam, D., Phan, V. T., Romanov, S., Finkle, D., Shu, J., Patel, V., Ton, T., Li, X., Loughhead, D. G., Nunn, P. A., Karr, D. E., Gerritsen, M. E., Funk, J. O., Owens, T. D., Verner, E., Brameld, K. A., Hill, R. J., Goldstein, D. M., and Taunton, J. (2015) Prolonged and tunable residence time using reversible covalent kinase inhibitors, Nat Chem Biol 11, 525–531.

[38] Lanning, B. R., Whitby, L. R., Dix, M. M., Douhan, J., Gilbert, A. M., Hett, E. C., Johnson, T. O., Joslyn, C., Kath, J. C., Niessen, S., Roberts, L. R., Schnute, M. E., Wang, C., Hulce, J. J., Wei, B., Whiteley, L. O., Hayward, M. M., and Cravatt, B. F. (2014) A road map to evaluate the proteome-wide selectivity of covalent kinase inhibitors, Nat Chem Biol 10, 760–767.

[39] Shraga, A., Olshvang, E., Davidzohn, N., Khoshkenar, P., Germain, N., Shurrush, K., Carvalho, S., Avram, L., Albeck, S., Unger, T., Lefker, B., Subramanyam, C., Hudkins, R. L., Mitchell, A., Shulman, Z., Kinoshita, T., and London, N. (2019) Covalent Docking Identifies a Potent and Selective MKK7 Inhibitor, Cell Chem Biol 26, 98–108 e105.

[40] Wolle, P., Hardick, J., Cronin, S. J. F., Engel, J., Baumann, M., Lategahn, J., Penninger, J. M., and Rauh, D. (2019) Targeting the MKK7-JNK (Mitogen-Activated Protein Kinase Kinase 7-c-Jun N-Terminal Kinase) Pathway with Covalent Inhibitors, J Med Chem 62, 2843–2848.

[41] Fischmann, T. O., Smith, C. K., Mayhood, T. W., Myers, J. E., Reichert, P., Mannarino, A., Carr, D., Zhu, H., Wong, J., Yang, R. S., Le, H. V., and Madison, V. S. (2009) Crystal structures of MEK1 binary and ternary complexes with nucleotides and inhibitors, Biochemistry 48, 2661–2674.

[42] Ordan, M., Pallara, C., Maik-Rachline, G., Hanoch, T., Gervasio, F. L., Glaser, F., Fernandez-Recio, J., and Seger, R. (2018) Intrinsically active MEK variants are differentially regulated by proteinases and phosphatases, Sci Rep 8, 11830.

[43] Akinleye, A., Furqan, M., Mukhi, N., Ravella, P., and Liu, D. (2013) MEK and the inhibitors: from bench to bedside, J Hematol Oncol 6, 27.

[44] Pearlman, S. M., Serber, Z., and Ferrell, J. E., Jr. (2011) A mechanism for the evolution of phosphorylation sites, Cell 147, 934–946.

[45] Dissmeyer, N., and Schnittger, A. (2011) Use of phospho-site substitutions to analyze the biological relevance of phosphorylation events in regulatory networks, Methods Mol Biol 779, 93–138.

[46] Fedorov, O., Niesen, F. H., and Knapp, S. (2012) Kinase inhibitor selectivity profiling using differential scanning fluorimetry, Methods Mol Biol 795, 109–118.

[47] Rega, C., Russo, R., Foca, A., Sandomenico, A., Iaccarino, E., Raimondo, D., Milanetti, E., Tornatore, L., Franzoso, G., Pedone, P. V., Ruvo, M., and Chambery, A. (2018) Probing the interaction interface of the GADD45beta/MKK7 and MKK7/DTP3 complexes by chemical cross-linking mass spectrometry, Int J Biol Macromol 114, 114–123.

[48] Papa, S., Monti, S. M., Vitale, R. M., Bubici, C., Jayawardena, S., Alvarez, K., De Smaele, E., Dathan, N., Pedone, C., Ruvo, M., and Franzoso, G. (2007) Insights into the structural basis of the GADD45beta-mediated inactivation of the JNK kinase, MKK7/JNKK2, J Biol Chem 282, 19029–19041.

[49] Salah, E., Ugochukwu, E., Barr, A. J., von Delft, F., Knapp, S., and Elkins, J. M. (2011) Crystal structures of ABL-related gene (ABL2) in complex with imatinib, tozasertib (VX-680), and a type I inhibitor of the triazole carbothioamide class, J Med Chem 54, 2359–2367.

[50] Zhang, J., Adrian, F. J., Jahnke, W., Cowan-Jacob, S. W., Li, A. G., Iacob, R. E., Sim, T., Powers, J., Dierks, C., Sun, F., Guo, G. R., Ding, Q., Okram, B., Choi, Y., Wojciechowski, A., Deng, X., Liu, G., Fendrich, G., Strauss, A., Vajpai, N., Grzesiek, S., Tuntland, T., Liu, Y., Bursulaya, B., Azam, M., Manley, P. W., Engen, J. R., Daley, G. Q., Warmuth, M., and Gray, N. S. (2010) Targeting Bcr-Abl by combining allosteric with ATP-binding-site inhibitors, Nature 463, 501–506.

[51] Kabsch, W. (2010) Xds, Acta Crystallogr D Biol Crystallogr 66, 125–132.

[52] Powell, H. R., Johnson, O., and Leslie, A. G. (2013) Autoindexing diffraction images with iMosflm, Acta Crystallogr D Biol Crystallogr 69, 1195–1203.

[53] Winn, M. D., Ballard, C. C., Cowtan, K. D., Dodson, E. J., Emsley, P., Evans, P. R., Keegan, R. M., Krissinel, E. B., Leslie, A. G., McCoy, A., McNicholas, S. J., Murshudov, G. N., Pannu, N. S., Potterton, E. A., Powell, H. R., Read, R. J., Vagin, A., and Wilson, K. S. (2011) Overview of the CCP4 suite and current developments, Acta Crystallogr D Biol Crystallogr 67, 235–242.

[54] McCoy, A. J., Grosse-Kunstleve, R. W., Adams, P. D., Winn, M. D., Storoni, L. C., and Read, R. J. (2007) Phaser crystallographic software, J Appl Crystallogr 40, 658–674.

[55] Emsley, P. (2017) Tools for ligand validation in Coot, Acta Crystallogr D Struct Biol 73, 203–210.

[56] Winn, M. D., Murshudov, G. N., and Papiz, M. Z. (2003) Macromolecular TLS refinement in REFMAC at moderate resolutions, Methods Enzymol 374, 300–321.

[57] Williams, C. J., Headd, J. J., Moriarty, N. W., Prisant, M. G., Videau, L. L., Deis, L. N., Verma, V., Keedy, D. A., Hintze, B. J., Chen, V. B., Jain, S., Lewis, S. M., Arendall, W. B., 3rd, Snoeyink, J., Adams, P. D., Lovell, S. C., Richardson, J. S., and Richardson, D. C. (2018) MolProbity: More and better reference data for improved all-atom structure validation, Protein Sci 27, 293–315.

[58] Rappsilber, J., Ishihama, Y., and Mann, M. (2003) Stop and go extraction tips for matrix-assisted laser desorption/ionization, nanoelectrospray, and LC/MS sample pretreatment in proteomics, Anal Chem 75, 663–670.

