## Supplementary information for "Catalytic domain plasticity of MKK7 reveals structural mechanisms of allosteric activation and new targeting opportunities"

```
MKK7      -----------MAA-------------------------------SSLEQKLSRLEA-KL   17
MEK1      ------------------------------------------------------------   0
MEK2      ------------------------------------------------------------   0
MEK4      ------------------------------------------------------------   0
MEK5      MLWLALGPFPAMENQVLVIRIKIPNSGAVDWTVHSGPQLLFRDVLDVIGQVLPEATTTAF   60
MEK6      ------------------------------------------------------------   0

MKK7      KQENREARRRIDLNLDISPQRPRPIIVITLSPAPAPSQRAALQLPLANDGGSRSPSSESS   77
MEK1      ------------------------------MPK-------------------KKPTP--I   9
MEK2      -----------------------------MLAR-------------------RKPVLPAL   12
MEK4      ------------------------------MAAPSPSGGGGS--GGGSGSGTPGPVGSPA   28
MEK5      EYE-DEDGDRITVRSDEEMKAMLSYYYSTVME-----------------QQVNGQLIEPL   102
MEK6      ------------------------------------------------------------   0

MKK7      PQHPTPPA----RPRHMLGLPSTLFTPR----------SMESIEI----------D-QKL   112
MEK1      QLNP-APD-------------------------GSAVNGTSSAETNLEALQKKLEELELD   43
MEK2      TINPTIAE-------------------------GPSPTSEGASEANLVDLQKKLEELELD   47
MEK4      PGHPAVSSMQGKRKALKLNFANPPFKSTARFTLNPNPTGVQNPHI----------E-RLR   77
MEK5      QIFPRACKPPGERNIHGLKVNTRA-----------GPSQHSSPAV----------SDSLP   141
MEK6      -----MSQSKGKKRNPGLKIPKEAFEQ---------------PQT----------S-STP   29

MKK7      -QEIMKQTGYLTIGGQRYQAEINDLENLGEMGSGTCGQVWKMRFRKTGHVIAVKQMRRSG   171
MEK1      EQQRKRLEAFLTQKQKVGELKDDDFEKISELGAGNGGVVFKVSHKPSGLVMARKLIHLEI   103
MEK2      EQQKKRLEAFLTQKAKVGELKDDDFERISELGAGNGGVVTKVQHRPSGLIMARKLIHLEI   107
MEK4      THSIESSGKLKISPEQHWDFTAEDLKDLGEIGRGAYGSVNKMVHKPSGQIMAVKRIRSTV   137
MEK5      SNSLKKSSAELKKILANGQMNEQDIRYRDTLGHGNGGTVYKAYHVPSGKILAVKVILLDI   201
MEK6      PRDLDSKACI-SIGNQNFEVKADDLEPIMELGRGAYGVVEKMRHVPSGQIMAVKRIRATV   88

MKK7      NKEENKRILMDLDVVLKSHDCPYIVQCFGTFITNTDVFIAMELMGTCAEKLKKR----MQ   227
MEK1      KPAIRNQIIRELQVLHE-CNSPYIVGFYGAFYSDGEISICMEHMDGGSLDQVLK----KA   158
MEK2      KPAIRNQIIRELQVLHE-CNSPYIVGFYGAFYSDGEISICMEHMDGGSLDQVLK----EA   162
MEK4      DEKEQKQLLMDLDVVMRSSDCPYIVQFYGALFREGDCWICMELMSTSFDKFYKYVYSVLD   197
MEK5      TLELQKQIMSELEILYK-CDSSYIIGFYGAFFVENRISICTEFMDGGSLDVYR-------   253
MEK6      NSQEQKRLLMDLDISMRTVDCPFTVTFYGALFREGDVWICMELMDTSLDKFYKQVIDK-G   147

MKK7      GPIPERILGKMTVAIVKALYYLKEKHGVIHRDVKPSNILLDERGQIKLCDFGISGRLVDS   287
MEK1      GRIPEQILGKVSIAVIKGLTYLREKHKIMHRDVKPSNILVNSRGEIKLCDFGVSGQLIDS   218
MEK2      KRIPEEILGKVSIAVLRGLAYLREKHQIMHRDVKPSNILVNSRGEIKLCDFGVSGQLIDS   222
MEK4      DVIPEEILGKITLATVKALNHLKENLKIIHRDIKPSNILLDRSGNIKLCDFGISGQLVDS   257
MEK5      -KMPEHVLGRIAVAVVKGLTYLW-SLKILHRDVKPSNMLVNTRGQVKLCDFGVSTQLVNS   311
MEK6      QTIPEDILGKIAVSIVKALEHLHSKLSVIHRDVKPSNVLINALGQVKMCDFGISGYLVDS   207

MKK7      KAKTRSAGCAAYMAPERIDPPDPTKPDYDIRADVWSLGISLVELATGQFPYKNCKT----   343
MEK1      MANSF-VGTRSYMSPERLQG-----THYSVQSDIWSMGLSLVEMAVGRYPIPPPDAKELE   272
MEK2      MANSF-VGTRSYMAPERLQG-----THYSVQSDIWSMGLSLVELAVGRYPIPPPDAKELE   276
MEK4      IAKTRDAGCRPYMAPERIDPSA-SRQGYDVRSDVWSLGITLYELATGRFPYPKWNS----   312
MEK5      IAKTY-VGTNAYMAPERISG-----EQYGIHSDVWSLGISFMELALGRFPYPQIQKN---   362
MEK6      VAKTIDAGCKPYMAPERINPEL-NQKGYSVKSDIWSLGITMIELAILRFPYDSWGT----   262

MKK7      -----------------------------------------DFEVLTKVLQEEPPLLPGH   362
MEK1      LMFGCQV----EGDAAETPPRPRTPGRPLSSYGMDSRPPMAIFELLDYIVNEPPPKLPS-   327
MEK2      AIFGRPVVDGEEGEPHSISPRPRPPGRPVSGHGMDSRPAMAIFELLDYIVNEPPPKLPN-   335
MEK4      -----------------------------------------VFDQLTQVVKGDPPQLSNS   331
MEK5      ------------------------------------QGSLMPLQLLQCIVDEDSPVLPV-   385
MEK6      -----------------------------------------PFQQLKQVVEEPSPQLPA-   280

MKK7      --MGFSGDFQSFVKDCLTKDHRKRPKYNKLLEHSFIKRYETLE-VDVASWFKDVMAKTES   419
MEK1      --GVFSLEFQDFVNKCLIKNPAERADLKQLMVHAFIKRSDAEE-VDFAGWLCSTIGLNQP   384
MEK2      --GVFTPDFQEFVNKCLIKNPAERADLKMLTNHTFIKRSEVEE-VDFAGWLCKTLRLNQP   392
MEK4      EEREFSPSFINFVNLCLTKDESKRPKYKELLKHPFILMYEERA-VEVACYVCKILDQMPA   390
MEK5      --GEFSEPFVHFITQCMRKQPKERPAPEELMGHPFIVQFNDGNAAVVSMWVCRALEERRS   443
MEK6      --DKFSAEFVDFTSQCLKKNSKERPTYPELMQHPFFTLHESKG-TDVASFVKLILGD---   334

MKK7      -PRTSGVLSQPHLPFFR 435
MEK1      STPTHAAGV-------- 393
MEK2      GTPTRTAV--------- 400
MEK4      -TPSSPMYVD------- 399
MEK5      -QQGPP----------- 448
MEK6      ----------------- 334
```

**Supplementary figure S1.** Sequence alignment of MAP2K family members. Green boxes indicate predicted MAPK docking domains, and pink boxes indicate the starting point of the C-terminal kinase domains. The N-terminal regulatory helices of MEK1 and MKK7 evident from the crystal structures are highlighted in yellow boxes.

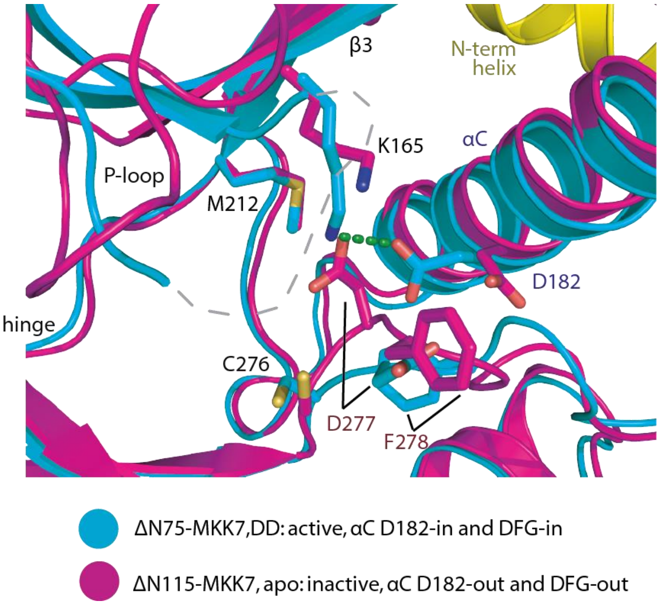


**Supplementary figure S2.** Structural superimposition of the inactive, apo kinase domain and active ΔN75-MKK7 harboring S287D and T291D mutations reveals potential steric clashes upon an in-swing of the αC Asp182 in the active form to the DFG-out Asp277 in the apo state.

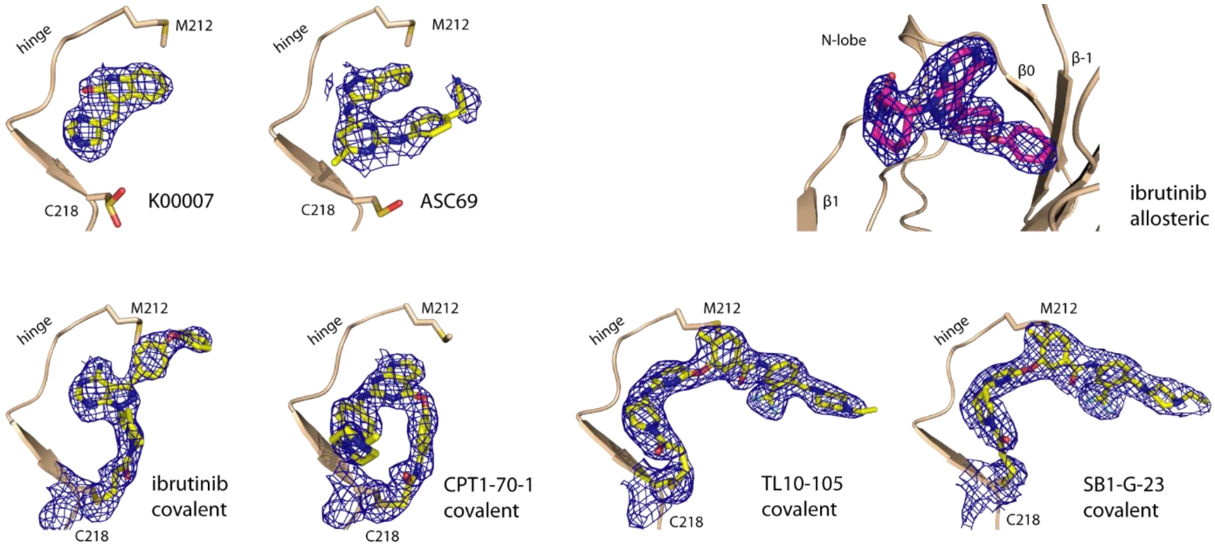


**Supplementary figure S3.** Omitted $|2F_O|-|Fc|$ electron density maps contoured at 1σ for the bound ligands and Cys218 in the case of covalent binding.

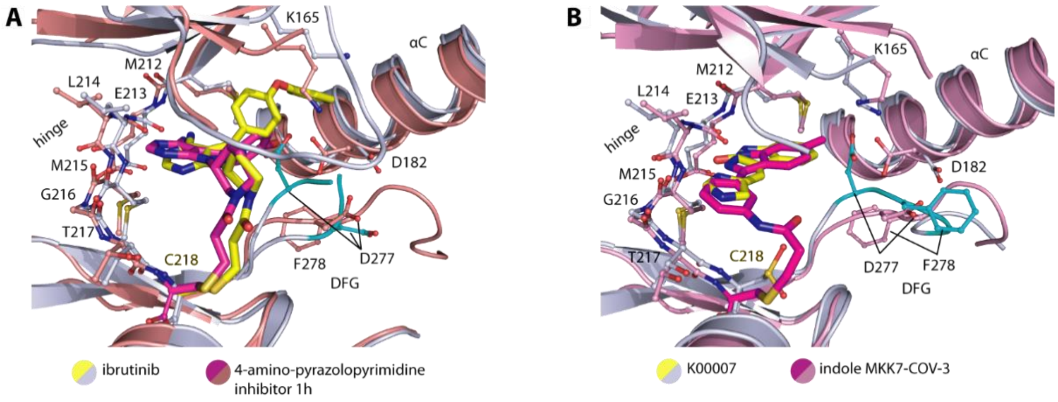


**Supplementary figure S4.** Comparison of inhibitor interactions in MKK7 between ibrutinib and 4-amino-pyrazolopyrimidine 1h (A) and K0007 and indole MKK7-COV-3 (B) reveals similar binding modes of the hinge interacting moieties in the pairs.

**Supplementary table 1.** Data collection and refinement statistics for MKK7 structures.

| **Complex** | **Apo ΔN115-MKK7, wild type** | **Apo ΔN115-MKK7, DD** | **ΔN75-MKK7, DD** | **ΔN115-MKK7, wild type: ibrutinib** | **ΔN115-MKK7, wild type: ibrutinib (allosteric only)** |
|---|---|---|---|---|---|
| **PDB accession code** | **6YFZ** | **6YG0** | **6YG1** | **6YG2** | **6YZ4** |
| **Beamline** | Diamond, i04 | Diamond, i02 | Diamond, i02 | Diamond, i04 | Diamond, i04 |
| ***Data Collection*** | | | | | |
| Resolution[a] (Å) | 30.41-1.90 (2.00-1.90) | 36.19-2.00 (2.11-2.00) | 64.83-2.22 (2.34-2.22) | 19.55-2.00 (2.11-2.00) | 25.58-1.70 (1.79-1.70) |
| Spacegroup | $P\ 2_12_12_1$ | $P\ 2_12_12_1$ | $C\ 2$ | $P\ 2_12_12_1$ | $P\ 2_12_12_1$ |
| Cell dimensions | a=57.5, b=74.5, c=81.8 Å α=β=γ= 90.0° | a=60.4, b=69.8, c=84.6 Å α=β=γ= 90.0° | a=127.2, b=67.9, c=142.8 Å α= γ= 90.0°, β=114.8° | a=53.1, b=75.0, c=86.7 Å α=β=γ= 90.0° | a=53.5, b=74.7, c=87.2 Å α=β=γ= 90.0° |
| No. unique reflections[a] | 28,220 (4,047) | 24,828 (3,550) | 54,756 (7,970) | 24,057 (3,473) | 39,163 (5,630) |
| Completeness[a] (%) | 99.6 (99.6) | 100.0 (100.0) | 99.8 (99.8) | 99.9 (100.0) | 99.9 (99.9) |
| I/σI[a] | 9.2 (2.0) | 13.8 (2.0) | 8.2 (2.0) | 12.1 (2.2) | 13.1 (2.3) |
| $R_{merge}$[a] | 0.103 (0.843) | 0.066 (0.952) | 0.090 (0.661) | 0.106 (0.901) | 0.085 (0.781) |
| CC (1/2) | 0.996 (0.621) | 0.997 (0.693) | 0.996 (0.675) | 0.997 (0.734) | 0.999 (0.552) |
| Redundancy[a] | 6.1 (6.1) | 6.7 (6.8) | 3.7 (3.5) | 7.3 (7.4) | 6.5 (6.5) |
| ***Refinement*** | | | | | |
| No. atoms in refinement (P/L/O)[b] | 2,386/ -/ 233 | 2,229/ -/ 89 | 7,344/ -/ 353 | 2,333/ 66/ 230 | 2,332/ 33/ 362 |
| B factor (P/L/O)[b] (Å²) | 38/ -/ 43 | 61/ -/60 | 64/ -/55 | 38/ 37/44 | 28/ 24/ 38 |
| $R_{fact}$ (%) | 19.2 | 19.4 | 20.4 | 17.9 | 16.9 |
| $R_{free}$ (%) | 23.7 | 24.5 | 23.2 | 23.0 | 20.6 |
| rms deviation bond[c] (Å) | 0.016 | 0.015 | 0.014 | 0.015 | 0.016 |
| rms deviation angle[c] (°) | 1.7 | 1.4 | 1.4 | 1.5 | 1.6 |
| ***Molprobity Ramachandran*** | | | | | |
| Favour (%) | 96.23 | 98.18 | 98.05 | 98.58 | 97.18 |
| Outlier (%) | 0 | 0 | 0.32 | 0 | 0 |
| Crystallization condition | 19% PEG3350, 0.15 M ammonium acetate, 0.1 M tris, pH 7.8 | 16% PEG3350, 0.25 M ammonium acetate, 0.1 M tris, pH 8.2 | 27% PEG3350, 0.2 M potassium thiocyanate, 10% ethylene glycol, 0.1 M bis-tris propane, pH 7.5 | 19% PEG3350, 0.1 M ammonium acetate, 0.1 M tris, pH 7.8 | 16% PEG3350, 0.25 M ammonium acetate, 0.1 M tris, pH 7.8 |

[a] Values in brackets show the statistics for the highest resolution shells; [b] P/L/O indicate protein, ligand molecules of interest, and other (water and solvent molecules), respectively; [c] rms indicates root-mean-square.

**Supplementary table 1 *cont.*** Data collection and refinement statistics for MKK7 structures.

| **Complex** | **ΔN115-MKK7, wild type: CPT1-70-1** | **ΔN115-MKK7, wild type: K00007** | **ΔN115-MKK7, wild type: ASC69** | **ΔN75-MKK7, DD: TL10-105** | **ΔN75-MKK7, DD: SB1-G-23** |
|---|---|---|---|---|---|
| **PDB accession code** | **6YG3** | **6YG4** | **6YG5** | **6YG6** | **6YG7** |
| **Beamline** | Diamond, i03 | Diamond, i04 | Diamond, i04 | Diamond, i02 | Diamond, i03 |
| ***Data Collection*** | | | | | |
| Resolution[a] (Å) | 30.38-2.05 (2.16-2.05) | 38.46-2.30 (2.42-2.30) | 39.64-2.40 (2.53-2.40) | 47.33-2.15 (2.27-2.15) | 61.83-2.20 (2.32-2.20) |
| Spacegroup | $P\ 2_1 2_1 2_1$ | $P\ 2_1 2_1 2_1$ | $P\ 2_1 2_1 2_1$ | $P\ 2_1$ | $P\ 2_1$ |
| Cell dimensions | a=58.2, b=64.5, c=85.5 Å<br>α=β=γ= 90.0° | a=54.1, b=72.6, c=83.1 Å<br>α=β=γ= 90.0° | a=59.7, b=68.0, c=84.7 Å<br>α=β=γ= 90.0° | a=72.7, b=69.93, c=73.8 Å<br>α= γ= 90.0°, β=119.4° | a=72.1, b=69.2, c=72.3 Å<br>α= γ= 90.0°, β=117.9° |
| No. unique reflections[a] | 20,596 (2,930) | 15,049 (2,162) | 13,948 (2,013) | 35,192 (5,117) | 32,046 (4,662) |
| Completeness[a] (%) | 99.1 (98.4) | 99.6 (99.8) | 99.5 (99.6) | 99.9 (99.9) | 99.9 (99.9) |
| I/σI[a] | 10.7 (2.0) | 11.2 (2.1) | 12.5 (2.0) | 10.0 (2.2) | 10.2 (2.0) |
| $R_{merge}$[a] (%) | 0.098 (0.935) | 0.066 (0.745) | 0.061 (0.730) | 0.101 (0.906) | 0.070 (0.770) |
| CC (1/2) | 0.998 (0.494) | 0.998 (0.691) | 0.997 (0.766) | 0.996 (0.659) | 0.998 (0.740) |
| Redundancy[a] | 6.6 (6.6) | 4.6 (4.7) | 4.7 (4.7) | 5.3 (5.1) | 4.9 (4.9) |
| ***Refinement*** | | | | | |
| No. atoms in refinement (P/L/O)[b] | 2,122/ 34/ 153 | 2,064/ 16/ 58 | 2,060/ 26/ 13 | 4,536/ 92/ 179 | 4,438/ 92/ 84 |
| B factor (P/L/O)[b] (Å$^2$) | 44/ 45/ 51 | 65/ 56/ 70 | 77/ 100/ 72 | 63/ 47/ 56 | 66/ 48/ 56 |
| $R_{fact}$ (%) | 18.8 | 21.1 | 23.1 | 20.4 | 20.7 |
| $R_{free}$ (%) | 23.5 | 25.5 | 28.2 | 23.9 | 25.1 |
| rms deviation bond[c] (Å) | 0.015 | 0.012 | 0.008 | 0.012 | 0.014 |
| rms deviation angle[c] (°) | 1.5 | 1.3 | 1.2 | 1.3 | 1.5 |
| ***Molprobity Ramachandran*** | | | | | |
| Favour (%) | 98.47 | 95.97 | 92.80 | 97.65 | 97.23 |
| Allowed (%) | 0 | 0 | 0.8 | 0 | 0 |
| Crystallization condition | 19% PEG3350, 0.25 M ammonium acetate, 0.1 M tris, pH 8.2 | 19% PEG3350, 0.1 M ammonium acetate, 0.1 M tris, pH 8.2 | 16% PEG3350, 0.2 M ammonium acetate, 0.1 M tris, pH 7.8 | 19% PEG3350, 0.15 M ammonium acetate, 0.1 M tris, pH 8.2 | 25% PEG3350, 0.1 M ammonium acetate, 0.1 M tris, pH 8.5 |

[a] Values in brackets show the statistics for the highest resolution shells; [b] P/L/O indicate protein, ligand molecules of interest, and other (water and solvent molecules), respectively; [c] rms indicates root-mean-square.

**Supplementary table 2.** Tm shift values from screening MKK7 against an in-house library of 360 compounds

| No. | ΔTm WT | ΔTm C202S | compound ID | compound names | source | smile |
|---|---|---|---|---|---|---|
| 1 | 24.8 | 11.2 | K06214a | TL10-105 | in-house collection | CCN1CCN(CC1)Cc1ccc(cc1C(F)(F)F)NC(c1ccc(C)c(c1)Oc1ccnc(N[C@H]2CCN(C2)C(C=C)=O)n1)=O |
| 2 | 14.1 | 8.2 | K06669a | SB1-G-23 | in-house collection | CCN1CCN(CC1)Cc1ccc(cc1C(F)(F)F)NC(c1ccc(C)c(c1)Oc1ccnc(N[C@@H]2CCN(C2)C(C=C)=O)n1)=O |
| 3 | 13.2 | 0.6 | K05736a | Ibrutinib | SelleckChem | C=CC(N1C[C@H](N2N=C(C3=CC=C(OC4=CC=CC=C4)C=C3)C5=C(N)N=CN=C52)CCC1)=O |
| 4 | 12.4 | 10.8 | K05783a | OTSSP167 | SelleckChem | ClC1=CC(C2=CC=C(N=CC(C(C)=O)=C3N[C@H]4CC[C@H](CN(C)C)CC4)C3=N2)=CC(Cl)=C1O |
| 5 | 10.8 | 6.2 | K06774a | CPT1-70-1 | in-house collection | CN1CCN(CC1)c1ccc(cc1)Nc1nc(c2cc[nH]c2n1)Oc1ccc(cc1)NC(C[Cl])=O |
| 6 | 9.3 | 7.1 | K05737a | HYJ-2-002-1 | in-house collection | CCN1CCN(CC1)Cc1ccc(cc1C(F)(F)F)NC(c1ccc(C)c(CCc2cnc3c(cc[nH]3)c2OC)c1)=O |
| 7 | 8.8 | 6.2 | K03428 | HG-6-71-01 | in-house collection | COC(=C1CNC(=CC(NC(C(=CC=C(C2C(F)(F)F)CN(CCN3CC)CC3)C=2)=O)=CC2)C=2C)C(=C(N=C1)N1)C=C1 |
| 8 | 8.7 | 5.1 | K03434a | XMD15-46 | in-house collection | O=C(NC(=CC(C(F)(F)F)=C(CN(CCN(CC)C1)C1)C1)C=1)C(=CC(C#CC(=CN=C(NC=C1)C12)C=2)=C(C)C1)C=1 |
| 9 | 5.5 | 3.5 | K03427 | HG-6-64-01 | in-house collection | CC(=CC=C1C(NC(=CC=C(C2C(F)(F)F)CN(CCN3CC)CC3)C=2)=O)C(=C1)C=CC(=C(C(=C1NC2)C=2)OC)C=N1 |
| 10 | 3.4 | | K02079a | KH-HP04 | in-house collection | ClC1=C(Cl)C=CC=C1/C=C2NC(/C(C\2=O)=C\C3=CC(Br)=C(O)C(Br)=C3)=O |
| 11 | 3.2 | 2.3 | K00007 | K00007 | Calbiochem (EMD) | C1C=CC=C(C=1N1)C(C1=O)=CC(N=CN1)=C1 |
| 12 | 2.7 | 4.9 | K00609 | ASC69 | MolPort | C(=C(N1)C)C(=NC=1NC1C=CC(=CC=1)CC#N)NC(=NNC1C(C2)C2)C=1 |
| 13 | 2.6 | | K03010a | GSK1326255A | GSK | CCCN(CCC1C(C=CC(NC(N=C(NC(=CC(F)=CC2)C=2C(N)=O)C(C=CN2)=C23)=N3)=C2OC)=C2)CC1 |
| 14 | 2.6 | | K00980a | GSK 650394 | Tocris | N1C=C(C=C(C=1N1)C(=C1)C(=CC(=C(C1)C(O)=O)C(CCC2)C2)C=1)C(=CC=CC1)C=1 |
| 15 | 2.3 | | K02255a | 229_0248_0168 | BioFocus | N(C(=C1)C(C=C2OC(F)(F)F)=CC=C2)(N=C2NCCCO)C(=N1)C=C2 |
| 16 | 2.3 | | K00781a | 5361852 | ChemBridge | C(C(N=C1NC(=CC2)C=CC=2OCC)=CS1)(N(C1C2)C=CC=2)=C(N=1)C.[Br].[Br] |
| 17 | 2.2 | | K00777a | JNK Inhib IX | SelleckChem | O=C(C1=CC=CC2=C1C=CC=C2)NC3=C(C#N)C4=C(S3)CCCC4 |
| 18 | 2.2 | | K00772 | T0503-6573 | Enamine | FC(F)(F)C1C=CC(SC(=N2)C(=CC3C=CN=CC=3)C#N)=C2C=1 |
| 19 | 2.2 | | K00984a | BX-795 | Axon Ligands | C(=NC=C(C1NCCCNC(C(=CC=C2)S2)=O)I)(N=1)NC(=CC(=CC1)NC(N(CCC2)C2)=O)C=1 |
| 20 | 2.2 | | K02259a | 229_0236_0196 | BioFocus | N(C(=C1)C(C=C2OC)=CN=C2)(N=C2NCC3C=C(C(O4)=CC=3)OC4)C(=N1)C=C2 |
| 21 | 2.1 | | K00524 | 294_0046_0333 | BioFocus | C(=C(C1)C(=CC2)C=CC=2N)(N=C2NC(C=C3[Cl])=CC=C3)N(N=1)C=C2 |
| 22 | 2.1 | 3.3 | K00040 | TBB (218697) | Calbiochem (EMD) | C(=C(C(=C(C1N=N2)[Br])[Br])[Br])(C=1N2)[Br] |
| 23 | 2.1 | | K02245a | 229_0131_0068 | BioFocus | N(C(=C1)C(C=C(C(O2)=C3)OC2)=C3)(N=C2NCC3=CC(=C(C(=C3)OC)OC)OC)C(=N1)C=C2 |
| 24 | 2.1 | | K02250a | 229_0033_0135 | BioFocus | N(C(=C1)C(=CC2N(C)C)C=CC=2)(N=C2NCCOC)C(=N1)C=C2 |
| 25 | 2.0 | | K02128a | 032_0036_0715 | BioFocus | C(=CC1C(C=C2)=CC=C2CC(C)C)(N=CN=1)NC1=CC(=C(C=C1)OC)OC |
| 26 | 2.0 | | K02227a | 190_0027_0168 | BioFocus | C(N(C1C(C=C2OC(F)(F)F)=CC=C2)C2)(=NC=1)C(=NC=2)NC(=CC1)C=CC=1O |
| 27 | 2.0 | | K02240a | 179_0223_0312 | BioFocus | N(C(=C1C(C=C2C(F)(F)F)=CC=C2)N2)(N=C1)C(=CC=2)NCC(=CC1[Cl])C=CC=1 |
| 28 | 1.9 | | K00488 | 229_0153_0279 | BioFocus | N(C(=C1)C(C=C2[Cl])=CC=C2)(N=C2NC(CO)CC)C(=N1)C=C2 |
| 29 | 1.9 | | K02257a | 229_0131_0280 | BioFocus | N(C(=C1)C(C=C2)=CC=C2OC)(N=C2NCC3=CC(=C(C(=C3)OC)OC)OC)C(=N1)C=C2 |
| 30 | 1.9 | | K02234a | 190_0141_0079 | BioFocus | C(N(C1C(=CC(=C(C2)OC)OC)C=2)C2)(=NC=1)C(=NC=2)NC(C=C1OC(C(F)F)(F)F)=CC=C1 |
| 31 | 1.9 | | K00523 | 294_0046_0068 | BioFocus | C(=C(C1)C(C=C(C(O2)=C3)OC2)=C3)(N=C2NC(C=C3[Cl])=CC=C3)N(N=1)C=C2 |
| 32 | 1.9 | | K02272a | 229_0146_0160 | BioFocus | N(C(=C1)C(C=C2)=CC=C2[Cl])(N=C2NCC(SC3)=CC=3)C(=N1)C=C2 |
| 33 | 1.8 | | K02231a | 190_0235_0284 | BioFocus | C(N(C1C(=CC2O)C=CC=2)C2)(=NC=1)C(=NC=2)NC(C=C1C(F)(F)F)=CC=C1 |
| 34 | 1.8 | | K00582a | AM-807/14961157 | Specs | C(=C(N1)N)(SC=1NC1=CC(=C(C=C1)[Cl])[Cl])C(C(C=CC1F)=CC=1)=O |
| 35 | 1.8 | | K00575a | AK-777/36504023 | Specs | C(=C(N1)N)(SC=1NC(C=C1)=CC=C1F)C(C(C=C1)=CC=C1[Cl])=O |
| 36 | 1.8 | | K02288a | 382_0087_0284 | BioFocus | C1(C(=CC2C(=CC3O)C=CC=3)C(=NC=2)N)C=C(C(=C(C=1)OC)OC)OC |
| 37 | 1.8 | | K00515 | 229_0254_0284 | BioFocus | N(C(=C1)C(=CC2O)C=CC=2)(N=C2NC(C(C)C)CO)C(=N1)C=C2 |
| 38 | 1.8 | | K00027 | Oxindole I | Calbiochem (EMD) | C(=C(C(N1)=O)C(=C1C=CC1)C=1)C(=CC=C1)N1 |
| 39 | 1.8 | | K00046 | Genistein | Calbiochem (EMD) | C1=C(C=C(C(=C1O1)C(C(=C1)C(=CC=C(C1)O)C=1)=O)O)O |
| 40 | 1.7 | | K02220a | 184_0219_0168 | BioFocus | N(C(C1NC(=CC2)C=CC=2OC)=N2)(C(=CN=1)C(C=C1OC(F)(F)F)=CC=C1)C=C2 |
| 41 | 1.7 | | K02129 | 032_0049_0005 | BioFocus | C(=CC1C(C=C2N)=CC=C2)(N=CN=1)NC(=CC1)C=CC=1[Cl] |
| 42 | 1.7 | | K02256a | 229_0131_0204 | BioFocus | N(C(=C1)C(C=C(C(N2)=C3)C=C2)=C3)(N=C2NCC3=CC(=C(C(=C3)OC)OC)OC)C(=N1)C=C2 |
| 43 | 1.7 | | K00517 | 229_0254_0313 | BioFocus | N(C(=C1)C(C=C2F)=CC=C2)(N=C2NC(C(C)C)CO)C(=N1)C=C2 |
| 44 | 1.7 | | K00060a | SB 218078 | Tocris | C1=CC=CC(=C1N1C2C3)C(=C1C1N4C(O2)C3)C(=C(C=1C(=C4C1)C=CC=1)C1=O)C(N1)=O |
| 45 | 1.7 | | K02246a | 229_0236_0004 | BioFocus | N(C(=C1)C(=CC2)C=CC=2N(C)C)(N=C2NCC3C=C(C(O4)=CC=3)OC4)C(=N1)C=C2 |
| 46 | 1.7 | | K00572a | AK-777/09836058 | Specs | C(=C(N1)N)(SC=1NC(C=CC1)=CC=1)C(C(=CC=C1F)C=C1)=O |
| 47 | 1.7 | | K02226a | 190_0021_4139 | BioFocus | C(N(C1C(C=C2)=CC=C2C)C2)(=NC=1)C(=NC=2)NC1=CC(=C(C(=C1)OC)OC)OC |
| 48 | 1.7 | | K02416a | 081_0283_0078 | BioFocus | C(C(=NC1)C2=CC=1)(C(=CC1NC(C=C3)=CC=C3C(C)C)C=NC=1)=CC=C2 |
| 49 | 1.6 | 0.6 | K00828 | PI3Kg inhib II (528108) | Calbiochem (EMD) | C(=C(C1)OC2(F)F)(C=C(C=1)C=C(SC(N1)=O)C1=O)O2 |
| 50 | 1.6 | | K00565a | Pfmrk Inhibitor (538140) | Calbiochem (EMD) | C(C(=C1)[Br])=CC(=C1C(C1=O)=CC(C(C=CC2F)=CC=2)=O)N1 |
| 51 | 1.6 | | K02236a | 190_0141_4145 | BioFocus | C(N(C1C(=CC2C(=O)C)C=CC=2)C2)(=NC=1)C(=NC=2)NC(C=C1OC(C(F)F)(F)F)=CC=C1 |
| 52 | 1.6 | | K00052 | Lavendustin A | Tocris | OC1=CC=C(O)C=C1CN(CC3=C(O)C=CC=C3)C2=CC=C(O)C(C(O)=O)=C2 |
| 53 | 1.6 | | K02438a | 382_0341_0280 | BioFocus | C(=CC1C(C=C2)=CC=C2OC)(C(=NC=1)N)C(=CC1)C=CC=1OCOC |
| 54 | 1.5 | | K02265a | 229_0131_4038 | BioFocus | N(C(=C1)C(=C(F)C2)C=CC=2)(N=C2NCC3=CC(=C(C(=C3)OC)OC)OC)C(=N1)C=C2 |
| 55 | 1.5 | | K00478 | 229_0033_0279 | BioFocus | N(C(=C1)C(C=C2[Cl])=CC=C2)(N=C2NCCOC)C(=N1)C=C2 |
| 56 | 1.5 | | K02124a | 032_0100_0085 | BioFocus | C(OC(C=C1)=CC=C1C(C=C1NC(C=C2)=CC=C2N(CCO)CC)=NC=N1)(F)(F)F |
| 57 | 1.5 | | K02214a | 174_0213_0339 | BioFocus | C(=NC1C(=CC2)C=CC=2F)(N(CC2)CCN2CCO)C(=NC=1)N |
| 58 | 1.5 | | K00518 | 229_0254_4145 | BioFocus | N(C(=C1)C(=CC2C(=O)C)C=CC=2)(N=C2NC(C(C)C)CO)C(=N1)C=C2 |
| 59 | 1.5 | | K00519 | 229_4051_0139 | BioFocus | N(C(=C1)C(=CC2C(=O)O)C=CC=2)(N=C2NCC(C3)C3)C(=N1)C=C2 |
| 60 | 1.5 | | K00507 | 229_0242_4145 | BioFocus | N(C(=C1)C(=CC2C(=O)C)C=CC=2)(N=C2NC(CC3)CCO3)C(=N1)C=C2 |
| 61 | 1.5 | | K02244a | 229_0131_0005 | BioFocus | N(C(=C1)C(=CC2N)C=CC=2)(N=C2NCC3=CC(=C(C(=C3)OC)OC)OC)C(=N1)C=C2 |
| 62 | 1.4 | | K00776a | Wee1/Chk1 inhibitor | Calbiochem (EMD) | C1=CC=C(C=C1)C2=CC3=C(C4=C(N3)C=CC(=C4)O)C5=C2C(=O)NC5=O.O |
| 63 | 1.4 | | K02122a | 032_0118_0071 | BioFocus | C(=CC1C(C=N2)=CC=C2)(N(CC2)CCN2CC(N(C(C=C2)=CC=C2)C)=O)N=CN=1 |
| 64 | 1.4 | | K02217a | 174_1002_0061 | BioFocus | C(N=C1C(=CC2)C=CC=2)(N(CC2)CCC2C(=O)O)=C(N=C1)N |
| 65 | 1.4 | | K02285a | 382_0005_4038 | BioFocus | C(=CC1C(=C(F)C2)C=CC=2)(C(=NC=1)N)C(=CC1N)C=CC=1 |

| | | | | | |
|---|---|---|---|---|---|
| 66 | 1.4 | K02239a | 179_0236_4139 | BioFocus | N(C(=C1C(=CC2)C=CC=2C)N2)(N=C1)C(=CC=2)NCC1C=C(C(O2)=CC=1)OC2 |
| 67 | 1.4 | K02160a | 093_0220_0168 | BioFocus | C(N(C1=C2)N=C2)(=CC(=N1)C(C=C1OC(F)(F)F)=CC=C1)N(CC1)CCN1C |
| 68 | 1.4 | K00510 | 229_0254_0086 | BioFocus | N(C(=C1)C(=CC2)C=CC=2C#N)(N=C2NC(C(C)C)CO)C(=N1)C=C2 |
| 69 | 1.4 | K00584a | AM-807/41062770 | Specs | C(=C(N1)N)(C(NC(=NO2)C=C2C)=O)SC=1NC(C=C1)=CC=C1F |
| 70 | 1.4 | K00585a | AM-807/41930348 | Specs | C(=C(N1)N)(SC=1NC1C=C(C(=CC=1)F)[Cl])C(C(=CC=C1OC)C=C1)=O |
| 71 | 1.4 | K02228a | 190_0254_0085 | BioFocus | C(N(C1C(C=C2)=CC=C2OC(F)(F)F)C2)(=NC=1)C(=NC=2)NC(C(C)C)CO |
| 72 | 1.3 | K00827 | PI3Kg inhibitor (528106) | Calbiochem (EMD) | N1=CC=NC(=C1C1)C=C(C=1)C=C(SC(N1)=O)C1=O |
| 73 | 1.3 | K02253a | 229_0244_0139 | BioFocus | N(C(=C1)C(=CC2C(=O)O)C=CC=2)(N=C2NCC(=CC3)C=CC=3OC)C(=N1)C=C2 |
| 74 | 1.3 | K00932a | Lapatinib | Sequoia Research Products | C(C(=N1)NC2=CC(=C(OCC(C=C3F)=CC=C3)C=C2)[Cl])(C=C2C(OC3CNCCS(=O)(=O)C)=CC=3)=C(N=C1)C=C2 |
| 75 | 1.3 | K00586 | AM-807/41930350 | Specs | C(=C(N1)N)(SC=1NC1C=C(C(=CC=1)F)[Cl])C(C(=CC1OC)C=CC=1)=O |
| 76 | 1.3 | K02213a | 225_1008_0280 | BioFocus | N(C(NC(=O)C(=CC1)C=CN=1)=C1C)(C(=N1)C1)C=C(N=1)C(=CC1)C=CC=1OC |
| 77 | 1.3 | K00576a | AM-807/12426021 | Specs | C(=C(N1)N)(SC=1NC(=CC=C1OC)C=C1)C(C(C=CC1OC)=CC=1)=O |
| 78 | 1.3 | K00768a | T0512-1192 | Enamine | FC(F)(F)C1C=CC(N=C(S2)C(=CC3=CC=CN3)C#N)=C2C=1 |
| 79 | 1.3 | K02736a | Scorzodihydrostilbene A | in-house collection | O[C@H]1[C@H](O)[C@@H](O)[C@H](OC2=CC=C(O)C(CCC3=CC=C(O)C(OC)=C3)=C2C(C)=O)O[C@@H]1CO |
| 80 | 1.3 | K00513 | 229_0254_0196 | BioFocus | N(C(=C1)C(C=C2OC)=CN=C2)(N=C2NC(C(C)C)CO)C(=N1)C=C2 |
| 81 | 1.3 | K02267a | 229_0231_0062 | BioFocus | N(C(=C1)C(C(=C2C3)C=CC=3)=CC=C2)(N=C2NCCC(=CN3)C=CC=3)C(=N1)C=C2 |
| 82 | 1.3 | K02241a | 179_0236_4027 | BioFocus | N(C(=C1C(=C([Cl])C2)C=CC=2)N2)(N=C1)C(=CC=2)NCC1C=C(C(O2)=CC=1)OC2 |
| 83 | 1.2 | K02218a | 174_0221_0192 | BioFocus | C(N=C1C(=CC2C(=O)NCCO)C=CC=2)(N(CC2)CCN2CCN(C)C)=C(N=C1)N |
| 84 | 1.2 | K00483 | 229_0153_0081 | BioFocus | N(C(=C1)C(=CC2NC(=O)C)C=CC=2)(N=C2NC(CO)CC)C(=N1)C=C2 |
| 85 | 1.2 | K00244b | GSK inhibitor XIII | Calbiochem (EMD) | CC1=CC(NC2=NC(C3=CC=CC=C3)=NC4=C2CCCC4)=NN1 |
| 86 | 1.2 | K02437a | 382_0327_0087 | BioFocus | C(=CC1C2=CC(=C(C(=C2)OC)OC)OC)(C(=NC=1)N)C(C=C1)=CC=C1C(=O)NCCN(C)C |
| 87 | 1.2 | K02264a | 229_0131_4027 | BioFocus | N(C(=C1)C(=C([Cl])C2)C=CC=2)(N=C2NCC3=CC(=C(C(=C3)OC)OC)OC)C(=N1)C=C2 |
| 88 | 1.2 | K00491 | 229_0153_0311 | BioFocus | N(C(=C1)C(=CN2)C=CC=2OC)(N=C2NC(CO)CC)C(=N1)C=C2 |
| 89 | 1.2 | K00063 | Tyrphostin A23 | in-house collection | OC1=C(O)C=C(\\C=C(/C#N)C#N)C=C1 |
| 90 | 1.2 | K02225a | 190_4059_0083 | BioFocus | N(C(=N1)C2NC(=C(OC(=CC3)C=CC=3)C3)C=CC=3)(C(=C1)C(=C(OC)C1)C=CC=1)C=CN=2 |
| 91 | 1.2 | K02130a | 032_0045_0280 | BioFocus | C(=CC1C(C=C2)=CC=C2OC)(N=CN=1)NC1C=C(C(N2)=CC=1)C=C2 |
| 92 | 1.2 | K00152b | 229_0242_0279 | BioFocus | N(C(=C1)C(C=C2[Cl])=CC=C2)(N=C2NC(CC3)CCO3)C(=N1)C=C2 |
| 93 | 1.2 | K02441a | 378_0021_0134 | BioFocus | C(=CC1C(NC2=CC(=C(C(=C2)OC)OC)OC)=O)(N(CC2)CCN2CCN(CC)CC)N=CN=1 |
| 94 | 1.2 | K02221a | 184_0219_0176 | BioFocus | N(C(C1NC(=CC2)C=CC=2OC)=N2)(C(=CN=1)C(=CC1C(=O)NCCN(C)C)C=CC=1)C=C2 |
| 95 | 1.1 | K02125a | 032_0106_0068 | BioFocus | C(=CC1C(C=C(C(O2)=C3)OC2)=C3)(N=CN=1)NC(=CC1N(C)C)C=CC=1 |
| 96 | 1.1 | K00239b | SU11652 | Calbiochem (EMD) | C(C=C1)(=CC(=C1N1)C(C1=O)=CC(=C(C(=C1C)C(=O)NCCN(CC)CC)C)N1)[Cl] |
| 97 | 1.1 | K02242a | 179_0134_0286 | BioFocus | N(C(=C1C(C=C(C2)C(C)C)=C(C=2)OC)N2)(C(N(CC3)CCN3CCN(CC)CC)=CC=2)N=C1 |
| 98 | 1.1 | K02258a | 229_0131_0285 | BioFocus | N(C(=C1)C(C=C2)=CC=C2CO)(N=C2NCC3=CC(=C(C(=C3)OC)OC)OC)C(=N1)C=C2 |
| 99 | 1.1 | K00578a | AM-807/12739008 | Specs | C(=C(N1)N)(SC=1NC(=CC1C(F)(F)F)C=CC=1)C(=O)C(=CC=C1)C=C1 |
| 100 | 1.1 | K03424a | ALW-II-41-27 | in-house collection | CCN(CCN1CC(=C(C(F)(F)F)C=C(NC(C(=CC=C(C)C2NC(C(=CC(C(=CC=C3)S3)=CN3)C=3)=O)C=2)=O)C2)C=2)CC 1 |
| 101 | 1.1 | K00531 | 5387887 | ChemBridge | N(C(N1)=O)(C(C(C1=O)=CNC(C=C1N(C)C)=CC=C1)=O)C(C=C1)=CC=C1F |
| 102 | 1.1 | K02159a | 093_0215_0063 | BioFocus | C(N(C1=C2)N=C2)(=CC(=N1)C(=C(OC(=CC1)C=CC=1)C1)C=CC=1)N(CCN1C)CCC1 |
| 103 | 1.1 | K02224a | 190_6937_0087 | BioFocus | C(N(C1C2C=C(C(=C(C=2)OC)OC)OC)C2)(=NC=1)C(=NC=2)NC(C=C1)=CC=C1C(C)C |
| 104 | 1.1 | K01737a | KH-CARB2 | in-house collection | ClC(C=CC1=C2N(C)C3=C1C(C#N)C(C)(C)NC3=O)=C2Cl |
| 105 | 1.1 | K02121a | 032_0134_0060 | BioFocus | C(=CC1C(C=C2C(F)(F)F)=CC(C(F)(F)F)=C2)(N(CC2)CCN2CCN(CC)CC)N=CN=1 |
| 106 | 1.1 | K00844a | STOCK6S-33492 | InterBioScreen | C(N=C1N)(=C(C(=N1)N1)N=C1)NC(C=C1)=CC=C1[Cl] |
| 107 | 1.0 | K02119a | 032_0113_0058 | BioFocus | C(=CC1C(C=C2C(C=C3)=CC=C3)=CC=C2)(N(CC2)CCN2CC(N(CC2)CC2)=O)N=CN=1 |
| 108 | 1.0 | K02716a | 7K-316S | Key Organics | C(=CC=CC1C(=NN=C(C2OC)C(=CC=CC3)C=3)N=2)C=1 |
| 109 | 1.0 | K00496 | 229_0242_0081 | BioFocus | N(C(=C1)C(=CC2NC(=O)C)C=CC=2)(N=C2NC(CC3)CCO3)C(=N1)C=C2 |
| 110 | 1.0 | K02268a | 229_0237_0062 | BioFocus | N(C(=C1)C(C(=C2C3)C=CC=3)=CC=C2)(N=C2NCC(=CC3)C=CN=3)C(=N1)C=C2 |
| 111 | 1.0 | K00712a | F0676-0387 | Life Chemicals | N(C(=N1)C(=CC2F)C=CC=2)(C(=N1)C1)N=C(C=1)SCC(NC(=CC1)C=CC=1[Cl])=O |
| 112 | 1.0 | K02237a | 190_0021_0004 | BioFocus | C(N(C1C(=CC2)C=CC=2N(C)C)C2)(=NC=1)C(=NC=2)NC1=CC(=C(C(=C1)OC)OC)OC |
| 113 | 1.0 | K00573a | AK-777/09836063 | Specs | C(=C(N1)N)(SC=1NC(=CC1)C=CC=1F)C(=O)C(C=C1)=CC=C1 |
| 114 | 1.0 | K02238a | 179_0236_0313 | BioFocus | N(C(=C1C(C=C2F)=CC=C2)N2)(N=C1)C(=CC=2)NCC1C=C(C(O2)=CC=1)OC2 |
| 115 | 1.0 | K00606b | ASC65 | in-house collection | N#CCC1C=CC(C=C1)=Nc1nc(Nc2cc([nH]n2)C2CC2)c2ccccc2n1 |
| 116 | 1.0 | K02262a | 229_0131_0311 | BioFocus | N(C(=C1)C(=CN2)C=CC=2OC)(N=C2NCC3=CC(=C(C(=C3)OC)OC)OC)C(=N1)C=C2 |
| 117 | 1.0 | K02219a | 174_0213_0289 | BioFocus | C(N=C1C(=CC2C=CC(=O)O)C=CC=2)(N(CC2)CCN2CCO)=C(N=C1)N |
| 118 | 1.0 | K00581 | AM-807/14961123 | Specs | C(=C(N1)N)(SC=1NC(C(F)=C1)=CC=C1)C(C(=CC1)C=CC=1[Br])=O |
| 119 | 1.0 | K02261a | 229_0131_0294 | BioFocus | N(C(=C1)C(=C(CO)C2)C=CC=2)(N=C2NCC3=CC(=C(C(=C3)OC)OC)OC)C(=N1)C=C2 |
| 120 | 1.0 | K00089 | piceatannol | Sigma Aldrich | OC1=CC(\C=C\C2=CC(O)=C(O)C=C2)=CC(O)=C1 |
| 121 | 0.9 | K00564a | Syk Inhibitor | Calbiochem (EMD) | C(=CC1)(C=C(C=12)C(C(N2)=O)=CC(=C1)C(=C2N1C)C=CC=C2)S(=O)(=O)N |
| 122 | 0.9 | K00230 | Debromohymenialdisine | Calbiochem (EMD) | C1(NC(C(N=1)=O)=C(CCN1)C(=C2C1=O)C=CN2)N |
| 123 | 0.9 | K00521 | 229_4051_0284 | BioFocus | N(C(=C1)C(=CC2O)C=CC=2)(N=C2NCC(C3)C3)C(=N1)C=C2 |
| 124 | 0.9 | K02120a | 032_0118_0079 | BioFocus | C(=CC1C(=CC(=C(C2)OC)OC)C=2)(N(CC2)CCN2CC(N(C(=CC2)C=CC=2)C)=O)N=CN=1 |
| 125 | 0.9 | K02927 | GW806742X | GSK | O=S(C1=CC=CC(NC2=NC=CC(N(C)C3=CC=C(NC(NC4=CC=C(OC(F)(F)F)C=C4)=O)C=C3)=N2)=C1)(N)=O |
| 126 | 0.9 | K00225c | JAK Inhibitor I | Calbiochem (EMD) | CC(C)(C)C1=NC2=C(N1)C3=C(C=C(C=C3)F)C4=C2C=CNC4=O |
| 127 | 0.9 | K00853a | STOCK6S-36822 | InterBioScreen | C(N=C1N)(=C(C(=N1)N1)N=C1)NC(C=C1F)=CC=C1 |
| 128 | 0.9 | K03184a | PF4708671 | Tocris | C1=NC=NC(=C1CC)N(CCN(C1)CC(NC(=C2C3)C=CC=3C(F)(F)F)=N2)C1 |
| 129 | 0.8 | K02271a | 229_0144_0087 | BioFocus | N(C(C1=CC(=C(C(=C1)OC)OC)OC)=C1)(N=C2NCC(OC3)=CC=3)C(=N1)C=C2 |
| 130 | 0.8 | K03001b | GSK978744A | in-house collection | C[C@@H](Oc1cc(sc1C(=O)N)n2cnc3ccc(OC[C@@H](O)CO)cc23)c4ccccc4Cl |
| 131 | 0.8 | K00835a | STOCK6S-32440 | InterBioScreen | C(C(=C1N2)N=C2)(=NC(=N1)N)N(CC1)CCN1 |
| 132 | 0.8 | K00119 | 190_0027_4139 | BioFocus | C(N(C1C(C=C2)=CC=C2C)C2)(=NC=1)C(=NC=2)NC(=CC1)C=CC=1O |
| 133 | 0.8 | K02243a | 229_0033_0004 | BioFocus | N(C(=C1)C(C=C2)=CC=C2N(C)C)(N=C2NCCOC)C(=N1)C=C2 |
| 134 | 0.8 | K02222 | 184_0088_0057 | BioFocus | N(C(C1NC(=CC2)C=CC=2N(CC2)CCO2)=N2)(C(=CN=1)C(=CC(=C(C1)C=C3)C=C3)C=1)C=C2 |
| 135 | 0.8 | K00597b | Sorafenib | LC Labs | C(=C(C(=CC1)[Cl])C(F)(F)F)C=1NC(NC(=CC=C(C1)OC(=CC(=NC2)C(NC)=O)C=2)C=1)=O |

| | | | | | |
|---|---|---|---|---|---|
| 136 | 0.8 | K02248a | 229_0131_0078 | BioFocus | N(C(C(C(=NC1)C2=CC=1)=CC=C2)=C1)(N=C2NCC3=CC(=C(C(=C3)OC)OC)OC)C(=N1)C=C2 |
| 137 | 0.8 | K00580a | AM-807/14146364 | Specs | C(=C(N1)N)(SC=1NC(=CC1)C=CC=1OC)C(C(=CC=C1[Cl])C=C1)=O |
| 138 | 0.8 | K00490 | 229_0153_0291 | BioFocus | N(C(=C1)C(=CC2CO)C=CC=2)(N=C2NC(CO)CC)C(=N1)C=C2 |
| 139 | 0.8 | K02247a | 229_0033_0078 | BioFocus | N(C(C(C(=NC1)C2=CC=1)=CC=C2)=C1)(N=C2NCCOC)C(=N1)C=C2 |
| 140 | 0.7 | K00238b | IKK-2 Inhibitor VI | Calbiochem (EMD) | C(=CC=C1C(=CC(=C2NC(=O)N)C(=O)N)S2)C=C1 |
| 141 | 0.7 | K02215a | 174_1002_0060 | BioFocus | C(=NC1C(=CC2C(F)(F)F)C=C(C(F)(F)F)C=2)(N(CC2)CCC2C(=O)O)C(=NC=1)N |
| 142 | 0.7 | K00041 | BPDQ (4-[(3-Bromophenyl)amino]-6,7-diaminoquinazoline) | in-house collection | C1=CC(=CC(=C1)Br)NC2=NC=NC3=CC(=C(C=C32)N)N |
| 143 | 0.7 | K00703a | F0676-0186 | Life Chemicals | N(C(=N1)C(=CC2)C=CC=2[Cl])(C(=N1)C1)N=C(C=1)SCC(NCC(OC1)CC1)=O |
| 144 | 0.7 | K00065 | Tyrphostin B48 | in-house collection | C(=C(C(=CC1CC(C(NC(=CC=CC2)C=2)=O)C#N)O)O)C=1 |
| 145 | 0.7 | K00839a | STOCK6S-42112 | InterBioScreen | C(N=C1N)(=C(C(=N1)N1)N=C1)NC(C=C1[Cl])=CC=C1 |
| 146 | 0.7 | K02533a | D481-2007 | ChemDiv | N(N=C1C(C=C2)=CC=C2[Cl])(C(=N1)N)C(C(=NC1)C=CC=1)=O |
| 147 | 0.7 | K02134a | 033_4009_0078 | BioFocus | N(C(N(CC1)CCO1)=C1)=C(C(C(=NC2)C3=CC=2)=CC=C3)C=C1 |
| 148 | 0.7 | K02260 | 229_6307_0279 | BioFocus | N(C(=C1)C(C=C2[Cl])=CC=C2)(N=C2NCC(=CC3)C=CC=3S(=O)(=O)N)C(=N1)C=C2 |
| 149 | 0.7 | K00090 | SL-327 | SelleckChem | C(C(=C(N)Sc1ccc(cc1)N)c1ccccc1C(F)(F)F)#N |
| 150 | 0.6 | K00579a | AM-807/12740052 | Specs | C(=C(N1)N)(SC=1NC(=CC1C(F)(F)F)C=CC=1)C(C(C=C1)=CC=C1OC)=O |
| 151 | 0.6 | K00505 | 229_0242_0339 | BioFocus | N(C(=C1)C(C=C2)=CC=C2F)(N=C2NC(CC3)CCO3)C(=N1)C=C2 |
| 152 | 0.6 | K00621a | ATM/ATR inhibitor | Calbiochem (EMD) | C(=CC=C1C(C(NC(C([Cl])([Cl])[Cl])NC(NC(C=CC(=C2[N+]([O-])=O)F)=C2)=S)=O)C(C=CC=C2)=C2)C=C1 |
| 153 | 0.6 | K00720a | F0676-0429 | Life Chemicals | N(C(=N1)C(=CC2F)C=CC=2)(C(=N1)C1)N=C(C=1)SCC(NC(=NC1)SC=1)=O |
| 154 | 0.6 | K00057 | Daphnetin (7,8-Dihydroxycoumarin) | Sigma Aldrich | O=C1C=CC2=CC=C(O)C(O)=C2O1 |
| 155 | 0.6 | K00039 | PD 169316 | Calbiochem (EMD) | FC(C=C1)=CC=C1C2=C(C3=CC=NC=C3)NC(C4=CC=C([N+]([O-])=O)C=C4)=N2 |
| 156 | 0.6 | K01734a | C713-0187 | ChemDiv | N(C(=N1)C(C=C2)=CC=C2F)(C(=N1)C1)N=C(C=1)NCCCN(CCC)CCC |
| 157 | 0.6 | K00798a | T5694244 | Enamine | C(=NN1)(C(NC(C=C2)=CC=C2F)=O)C(=C1C=C1)C=C1 |
| 158 | 0.6 | K00616a | Aurora/Cdk Inhibitor | Calbiochem (EMD) | C(=CC1)(C=CC=1NC1=NN(C(=N1)N)C(=O)C(C(=CC=C1)F)=C1F)S(=O)(=O)N |
| 159 | 0.6 | K00548a | 5214367 | ChemBridge | C(NC1=CC(=C(C=C1)[Cl])[Cl])(NC(=CC1)C=CC=1OCCCCC)=O |
| 160 | 0.5 | K02291a | 378_4051_0107 | BioFocus | C(=CC1C(NCC(C2)C2)=O)(N(CC2)CCN2C(=CC(=C(C2)OC)OC)C=2)N=CN=1 |
| 161 | 0.5 | K00699a | F0676-0126 | Life Chemicals | N(C(=N1)C(C=C2)=CC=C2[Cl])(C(=N1)C1)N=C(C=1)SCC(N(CC1)CCO1)=O |
| 162 | 0.5 | K00080 | HA-1004 (HCl) | Sigma Aldrich | O=S(C1=CC=CC2=CN=CC=C21)(NCCNC(N)=N)=O.Cl.Cl |
| 163 | 0.5 | K00722a | F0676-0458 | Life Chemicals | N(C(=N1)C(=CC2F)C=CC=2)(C(=N1)C1)N=C(C=1)SCC(=NC1)C=CC=1 |
| 164 | 0.5 | K00024 | Cdk4 Inhibitor C228125 | Toronto Research Chemicals | C1=CC=C2C(=C1)C3=C4C(=C5C6=C(C=C(C=C6)Br)NC5=C3N2)C(=O)NC4=O |
| 165 | 0.5 | K02216a | 174_0220_0087 | BioFocus | C(N=C1C2=CC(=C(C(=C2)OC)OC)OC)(N(CC2)CCN2C)=C(N=C1)N |
| 166 | 0.5 | K00731a | F0676-0862 | Life Chemicals | N(C(=N1)C(=CC(=C(C2)OC)OC)C=2)(C(=N1)C1)N=C(C=1)SCC(NC(=CC1[Cl])C=CC=1)=O |
| 167 | 0.5 | K00713a | F0676-0389 | Life Chemicals | N(C(=N1)C(=CC2F)C=CC=2)(C(=N1)C1)N=C(C=1)SCC(NC(C(=C1)F)=CC=C1F)=O |
| 168 | 0.5 | K00769a | T0518-8465 | Enamine | CC(=O)NC1C=CC(C=C(C#N)C2=NC(C=CC=C3)=C3S2)=CC=1 |
| 169 | 0.5 | K00837a | STOCK1N-70806 | InterBioScreen | N1=C(C(=C(N=C1N)NCCC1=CC(=C(C=C1)OC)OC)N1)NC=1 |
| 170 | 0.5 | K00841a | STOCK1N-70809 | InterBioScreen | N1=C(C(=C(N=C1N)NCC(C=C1)=CC=C1)N1)NC=1 |
| 171 | 0.5 | K02127a | 032_0131_0135 | BioFocus | C(C(=C1)OC)(=C(C=C1CNC(=CC1C(C=C2N(C)C)=CC=C2)N=CN=1)OC)OC |
| 172 | 0.5 | K00574a | AK-777/09836064 | Specs | C(=C(N1)N)(SC=1NC(=CC1)C=CC=1F)C(C(C=C1)=CC=C1F)=O |
| 173 | 0.5 | K00642a | 5225597 | ChemBridge | C(C(N1)=O)(C(=C1C1)C=C(C=1)C)=NNC(C(C=C1O)=CC=C1)=O |
| 174 | 0.5 | K00710a | F0676-0384 | Life Chemicals | N(C(=N1)C(=CC2F)C=CC=2)(C(=N1)C1)N=C(C=1)SCC(NC(=CC1F)C=CC=1)=O |
| 175 | 0.5 | K00994a | Dovitinib | LC Labs | C(=CC=C1)(C(=C1C1)C(=C(C1=O)C(=NC(=C1C2)C=C(C=2)N(CCN(C2)C)C2)N1)N)F |
| 176 | 0.5 | K00847a | STOCK6S-41767 | InterBioScreen | N1=C(C(=C(N=C1N)NCC(=CC1[Cl])C=CC=1)N1)NC=1 |
| 177 | 0.4 | K00625a | VX-680 | Tocris | CN1CCN(C2=NC(SC3=CC=C(C=C3)NC(C4CC4)=O)=NC(NC5=NNC(C)=C5)=C2)CC1 |
| 178 | 0.4 | K03442a | 9043251 | ChemBridge | C(C(NC1=CC(=C(N2)C=C1)C=N2)=O)(C(=C1)OC)=CC(=C1)[Cl] |
| 179 | 0.4 | K00701a | F0676-0156 | Life Chemicals | N(C(=N1)C(C=C2)=CC=C2[Cl])(C(=N1)C1)N=C(C=1)SCC(NC(=CC1OC)C=CC=1)=O |
| 180 | 0.4 | K03453a | STOCK1S-33760 | InterBioScreen | C(C(=N1)S2)(C(=C2C2N)N=CN=2)=C(C(=C1N(CC1)CCO1)C1)CCC1 |
| 181 | 0.4 | K00933a | Nilotinib | Sequoia Research Products | C(=CC(=CC1)C(NC(=CC(=CC2N(C=NC3C)C=3)C(F)(F)F)C=2)=O)(C=1C)NC(=NC(=CC1)C(=CN=CC2)C=2)N=1 |
| 182 | 0.4 | K00044 | ML 3163 (no. 475800) | Calbiochem (EMD) | C(=CC(=CC1)F)C=1C(NC(=N1)SCC(=CC=C(C2)S(C)=C)C=2)=C1C(=CC=NC1)C=1 |
| 183 | 0.4 | K00555a | 6626103 | ChemBridge | N(C(N1)=O)(C(C(C1=O)=CNC(C=C1)=CC=C1F)=O)C |
| 184 | 0.4 | K00709a | F0676-0370 | Life Chemicals | N(C(=N1)C(=CC2F)C=CC=2)(C(=N1)C1)N=C(C=1)SCC(=O)N |
| 185 | 0.4 | K02071a | KH-CARB3A | in-house collection | ClC(C=CC1=C2N(C)C3=C1C(C#N)C4(CCCCC4)NC3=O)=C2Cl |
| 186 | 0.4 | K00498 | 229_0242_0139 | BioFocus | N(C(=C1)C(=CC2C(=O)O)C=CC=2)(N=C2NC(CC3)CCO3)C(=N1)C=C2 |
| 187 | 0.4 | K00700a | F0676-0146 | Life Chemicals | N(C(=N1)C(C=C2)=CC=C2[Cl])(C(=N1)C1)N=C(C=1)SCC(NC(=CC1F)C=CC=1)=O |
| 188 | 0.4 | K00721a | F0676-0454 | Life Chemicals | N(C(=N1)C(=CC2F)C=CC=2)(C(=N1)C1)N=C(C=1)SCC(C=N1)=CC=C1 |
| 189 | 0.4 | K00050 | Lavendustin C | Sigma Aldrich | Oc1ccc(O)c(CNc2ccc(O)c(c2)C(=O)O)c1 |
| 190 | 0.4 | K00786a | 7974871 | ChemBridge | C(=NC(C1=C2)=CC(=C2C)C)(C(=CC(=CC2)C=CN=2)C#N)N1 |
| 191 | 0.4 | K00704a | F0676-0191 | Life Chemicals | N(C(=N1)C(=CC2)C=CC=2[Cl])(C(=N1)C1)N=C(C=1)SCC(NC(=NC1)SC=1)=O |
| 192 | 0.4 | K03452a | STOCK1S-37479 | InterBioScreen | C(C(=N1)S2)(C(=C2C2N)N=CN=2)=C(C(=C1N(CC1)CCO1)C1)CC(O1)(C)C |
| 193 | 0.4 | K00937d | TAE684 | SelleckChem | CN1CCN(C2CCN(C3=CC=C(NC4=NC=C(Cl)C(NC5=CC=CC=C5S(C(C)C)(=O)=O)=N4)C(OC)=C3)CC2)CC1 |
| 194 | 0.4 | K00094 | 040_0232_0074 | BioFocus | C(C(N=C1)=CN=C1NCC(=CC1)C=CC=1)(=CS1)C=C1 |
| 195 | 0.4 | K00849a | STOCK6S-40137 | InterBioScreen | C(N=C1N)(=C(C(=N1)N1)N=C1)NC(C(OC)=C1)=CC=C1 |
| 196 | 0.4 | K01938b | 9041990 | ChemBridge | N(C(C(=C1N2)N=C2)=NC=N1)(CC1C)CC(O1)C |
| 197 | 0.3 | K00072 | Cdk1 Inhibitor III | Calbiochem (EMD) | NC1=NC(NC2=CC=C(S(N)(=O)=O)C=C2)=NN1C(NC3=C(F)C=CC=C3F)=S |
| 198 | 0.3 | K02223 | 190_6937_0083 | BioFocus | N(C(=N1)C2NC(C=C3)=CC=C3C(C)C)(C(=C1)C(=C(OC)C1)C=CC=1)C=CN=2 |
| 199 | 0.3 | K00852a | STOCK6S-36498 | InterBioScreen | N1=C(C(=C(N=C1N)NCC(C=C1)=CC=C1OC)N1)NC=1 |
| 200 | 0.3 | K00767a | T0507-3237 | Enamine | CC(=C(C#N)C1=NC(C=CC=C2)=C2S1)C1C=CC([Cl])=CC=1 |
| 201 | 0.3 | K00757a | T5674742 | Enamine | N#CC(=CC1C=CC=NC=1)C1=NC(C=CC=C2)=C2S1 |
| 202 | 0.3 | K00617a | Dasatinib | SelleckChem | CC1=NC(NC2=NC=C(C(NC3=C(C=CC=C3Cl)C)=O)S2)=CC(N4CCN(CC4)CCO)=N1 |
| 203 | 0.3 | K00702a | F0676-0180 | Life Chemicals | CCOC(CNC(CSc1ccc2nnc(c3ccc(cc3)[Cl])n2n1)=O)=O |

| | | | | | |
|---|---|---|---|---|---|
| 204 | 0.3 | K00806a | O6-Benzylguanine | AXXORA | C(=CC1)(C=CC=1)COC(N=C1N)=C2C(=N1)NC=N2 |
| 205 | 0.3 | K00075 | HA-1077 (Fasudil; BRD7868) | Sigma Aldrich | O=S(C1=CC=CC2=CN=CC=C21)(N3CCNCCC3)=O |
| 206 | 0.3 | K00062 | Alsterpaullone | Sigma Aldrich | O=C1CC2=C(C3=CC=CC=C3N1)NC4=C2C=C([N+]([O-])=O)C=C4 |
| 207 | 0.3 | K00048 | AG 1879 (PP2) | SelleckChem | CC(C)(C)n1c2c(c(c3ccc(cc3)[Cl])n1)c(N)ncn2 |
| 208 | 0.3 | K00037 | SB 202190 | Calbiochem (EMD) | C(=C(C=CC1C(=C(NC2C(=CC=C(C3)O)C=3)C(=CC=NC3)C=3)N=2)F)C=1.[Cl] |
| 209 | 0.3 | K00566a | ERK Inhibitor | Calbiochem (EMD) | C(=C(C1)OCCC)C=C(C=1)C=C(C(N(C1=O)CCN)=O)S1 |
| 210 | 0.3 | K00846a | STOCK6S-37451 | InterBioScreen | N1=C(C(=C(N=C1N)NCC1=CC(=C(C=C1)[Cl])[Cl])N1)NC=1 |
| 211 | 0.2 | K00729a | F0676-0846 | Life Chemicals | N(C(=N1)C(=CC(=C(C2)OC)OC)C=2)(C(=N1)C1)N=C(C=1)SCC(=O)N |
| 212 | 0.2 | K03423 | ALW-II-38-3 | in-house collection | CC(N=CN1C(=CC(C(NC(=CC=C(C)C2NC(C(=CC=N3)O3)=O)C=2)=O)=CC2C(F)(F)F)C=2)=C1 |
| 213 | 0.2 | K02703a | Reversine | Calbiochem (Merck) | N1=C(N=C(C(=C1N1)N=C1)NC(CCCC1)C1)NC(=CC=C(C1)N(CCOC2)C2)C=1 |
| 214 | 0.2 | K00058 | Wedelolactone | Calbiochem (EMD) | C(=CC(=C1)OC)(C(=C1O1)C(=C(C1=O)C(=C1C2)C=C(C=2O)O)O1)O |
| 215 | 0.2 | K00209 | MEK1/2 Inhibitor | Calbiochem (EMD) | C1=CC(=C(C=C1)C(C#N)=C(N)SC(C=CC1N)=CC=1)C(F)(F)F |
| 216 | 0.2 | ATP | | | |
| 217 | 0.2 | K00714a | F0676-0414 | Life Chemicals | N(C(=N1)C(=CC2F)C=CC=2)(C(=N1)C1)N=C(C=1)SCC(NC(=CC1NC(=O)C)C=CC=1)=O |
| 218 | 0.2 | K02487a | HA-156 | Toronto Research Chemicals | N(CCN1S(C(=C(C2=C(C3)[Cl])C=CN=C2)C=3)(=O)=O)CC1.[Cl].[Cl] |
| 219 | 0.2 | K00240a | Gefitinib | LC Labs | COc1cc2c(cc1OCCCN1CCOCC1)c(Nc1ccc(c(c1)[Cl])F)ncn2 |
| 220 | 0.2 | K01786a | KH-CARB11 | in-house collection | ClC(C=CC1=C2N(C)C3=C1C(C#N)C4(CCN(CC)CC4)NC3=O)=C2Cl |
| 221 | 0.1 | K00760a | T0518-4225 | Enamine | CC1=CC=C(C=C(C#N)C2=NC(C=CC=C3)=C3S2)O1 |
| 222 | 0.1 | K03448a | 7535026 | ChemBridge | C(C(=C1C(=O)N)N)(=C(S1)N1)C(=C(C=1C1)CCC1)C(OC1)=CC=1 |
| 223 | 0.1 | K00705a | F0676-0216 | Life Chemicals | N(C(=N1)C(C=C2)=CC=C2[Cl])(C(=N1)C1)N=C(C=1)SCC(C=N1)=CC=C1 |
| 224 | 0.1 | K00730a | F0676-0859 | Life Chemicals | N(C(=N1)C(=CC(=C(C2)OC)OC)C=2)(C(=N1)C1)N=C(C=1)SCC(NC(C(F)=C1)=CC=C1)=O |
| 225 | 0.1 | K00931a | ART 17513474 | Asinex | C(=C(C(=CC1)OC=1)N1)(C(=C1N=C1)C=C1)CN(CCN1C)CC1 |
| 226 | 0.1 | K00789a | T0502-1077 | Enamine | N(=C(C(=CC(C=C1)=CC=N1)C#N)S1)C(=C1)C(C=C1)=CC=C1 |
| 227 | 0.1 | K00528 | 5981310 | ChemBridge | N(C(N1)=O)(C(C(C1=O)=CNC1=CC(=C(C=C1)C)[Cl])=O)C(C=C1)=CC=C1 |
| 228 | 0.1 | K00708a | F0676-0364 | Life Chemicals | N(C(=N1)C(=CC2F)C=CC=2)(C(=N1)C1)N=C(C=1)SCC(N(CC1)CCO1)=O |
| 229 | 0.1 | K00930a | ART 17513448 | Asinex | C(=C(C(=CC1)OC=1)N1)(C(=C1N=C1)C=C1)CN(CCO1)CC1 |
| 230 | 0.1 | K00130 | 225_1008_0160 | BioFocus | N(C(NC(=O)C(C=C1)=CC=N1)=C1C)(C(=N1)C1)C=C(N=1)C(=CC1)C=CC=1[Cl] |
| 231 | 0.1 | K00569d | SB-220025 | GSK | C1CNCCC1N2C=NC(=C2C3=NC(=NC=C3)N)C4=CC=C(C=C4)F |
| 232 | 0.1 | K02149a | 093_0256_0204 | BioFocus | C(N(C1=C2)N=C2)(=CC(=N1)C(C=C(C(N1)=C2)C=C1)=C2)N(C(CO)C1)CC1 |
| 233 | 0.1 | K00719a | F0676-0424 | Life Chemicals | N(C(=N1)C(=CC2F)C=CC=2)(C(=N1)C1)N=C(C=1)SCC(NCC(OC1)CC1)=O |
| 234 | 0.1 | K02400a | 9051337 | ChemBridge | N1NC(=C(C=C2NC(CC(=C(OC)C3)C=CC=3)=O)C=1)C=C2 |
| 235 | 0.1 | K00547a | 5809868 | ChemBridge | C(C(N(C(N1)=O)C)=O)(C1=O)=CC(NC1)=CC=1 |
| 236 | 0.1 | K00976a | Cdk/CKI Inhibitor, (R)-DRF053 | Calbiochem (EMD) | N(C(=C(C(=NC1NC(C)CO)N2C(C)C)N=C2)N=1)C(=CC=C(C1)C(=NC=CC2)C=2)C=1 |
| 237 | 0.1 | K00477a | Imatinib | LC Labs | Cc1ccc(cc1Nc1nccc(c2cccnc2)n1)NC(c1ccc(CN2CCN(C)CC2)cc1)=O |
| 238 | 0.1 | K00938a | ML-7 | Sigma Aldrich | C(CN(CCN1)S(C(=C(C(=CC2)C3I)C=CC=3)C=2)(=O)=O)C1 |
| 239 | 0.1 | K02249a | 229_0236_0078 | BioFocus | N(C(C(C(=NC1)C2=CC=1)=CC=C2)=C1)(N=C2NCC3C=C(C(O4)=CC=3)OC4)C(=N1)C=C2 |
| 240 | 0.1 | K00784a | 6339039 | ChemBridge | C(=N1)(N(C(C1=CC1)=CC=1)C)C(=CC(C=CN1)=CC=1)C#N |
| 241 | 0.1 | K00707a | F0676-0361 | Life Chemicals | N(C(=N1)C(=CC2F)C=CC=2)(C(=N1)C1)N=C(C=1)SCC(N(CC1)CC1)=O |
| 242 | 0.1 | K03863a | PD 166285 | Tocris | CN(C1=NC(NC4=CC=C(OCCN(CC)CC)C=C4)=NC=C1C=C2C3=C(Cl)C=CC=C3Cl)C2=O.Cl.Cl |
| 243 | 0.1 | K02401a | 9051854 | ChemBridge | C(C(NC1=CC(=C(N2)C=C1)C=N2)=O)(C(=C1OC)OC)=CC=C1 |
| 244 | 0.1 | K00464 | ST051029 | TimTec | C(C(N1)=O)(C(N=C1)=C1)=C([Cl])C=C1 |
| 245 | 0.0 | K00788a | T0501-6793 | Enamine | C(=N1)(C(=CC(C=C2)=CC=N2)C#N)SC(=C1CC1)CC1C |
| 246 | 0.0 | K00851a | STOCK6S-32524 | InterBioScreen | N1=C(C(=C(N=C1N)NCC(C([Cl])=C1)=CC=C1)N1)NC=1 |
| 247 | 0.0 | K00724a | F0676-0462 | Life Chemicals | N(C(=N1)C(=CC2F)C=CC=2)(C(=N1)C1)N=C(SC(C(=O)OCC)CC)C=1 |
| 248 | 0.0 | K01756a | KH-DTCMA | in-house collection | ClC(C=CC1=C2N(C)C3=C1C(CNC(C)=O)C(C)(C)NC3=O)=C2Cl |
| 249 | 0.0 | K00247 | Chk2 inhibitor II | Calbiochem (EMD) | NC(=O)C(C=C(C12)N=C(N2)C(C=CC(=C2)OC3C=CC(=CC=3)[Cl])=C2)=CC=1 |
| 250 | 0.0 | K00549a | 5214400 | ChemBridge | C(NC1C=C(C(=CC=1)[Cl])[Cl])(NC(=CC1)C=CC=1C(=O)OCC)=O |
| 251 | 0.0 | K00706a | F0676-0226 | Life Chemicals | N(C(=N1)C(C=C2)=CC=C2[Cl])(C(=N1)C1)N=C(SC(C(=O)OC)C)C=1 |
| 252 | 0.0 | K00711a | F0676-0386 | Life Chemicals | N(C(=N1)C(=CC2F)C=CC=2)(C(=N1)C1)N=C(C=1)SCC(NC(=CC1[Cl])C=CC=1)=O |
| 253 | 0.0 | K00739a | F0676-0943 | Life Chemicals | N(C(=N1)C(=CC(=C(C2)OC)OC)C=2)(C(=N1)C1)N=C(C=1)SCC(=O)O |
| 254 | 0.0 | K00233 | Chelerythrine Chloride | AXXORA | C1C=C(C(=C(C=1OC)OC)C1)C(C([N+]=1C)=C(C1C=C(C2O3)OC3)C=2)=CC=1 |
| 255 | 0.0 | K00398 | STOCK2S-67628 | InterBioScreen | C1(=C(C(=C(S1)N1)C(=C(C=1CC(C)C)C1)CCC1)N1)C(NC=1)=O |
| 256 | 0.0 | K00076 | Indirubin-3monoxime | Calbiochem (EMD) | InChI=1S/C16H11N3O2/c20-16-13(9-5-1-3-7-11(9)18-16)15-14(19-21)10-6-2-4-8-12(10)17-15/h1-8,17,21H,(H,18,20) |
| 257 | 0.0 | K00077d | K252a | Tocris | [H][C@](C[C@]8(O)[C@@](OC)=O)(O[C@]78C)N5C3=C2N7C1=CC=CC=C1C2=C(CNC6=O)C6=C3C4=CC=CC=C45 |
| 258 | 0.0 | K00100 | 229_0224_0142 | BioFocus | N(C(=C1)C(OC2)=CC=2)(N=C2NCC(C=C3)=CC=C3F)C(=N1)C=C2 |
| 259 | 0.0 | K00778a | PIM1 Inhib II | Calbiochem (EMD) | C1=C(NC(C(=C1C(=CC=CC1)C=1)C#N)=O)C(=CC(=CC1)[Br])C=1O |
| 260 | 0.0 | K00833a | STOCK4S-62175 | InterBioScreen | N1C(=C(C(=NC=1N)SCC(=C(C1)[Cl])C=CC=1[Cl])N1)NC=1 |
| 261 | 0.0 | K00959a | Pazopanib | Sequoia Research Products | C(=C(C(=CC1NC(=NC(=CC2)N(C(=CC(C(C3)=C(N4C)C)=N4)C=3)C)N=2)S(=O)(N)=O)C)C=1 |
| 262 | 0.0 | K02251a | 229_0131_0167 | BioFocus | N(C(=C1)C(=CC(=C(C2)C)C)C=2)(N=C2NCC3=CC(=C(C(=C3)OC)OC)OC)C(=N1)C=C2 |
| 263 | 0.0 | K02252a | 229_0236_0167 | BioFocus | N(C(=C1)C(=CC(=C(C2)C)C)C=2)(N=C2NCC3C=C(C(O4)=CC=3)OC4)C(=N1)C=C2 |
| 264 | 0.0 | K02254a | 229_0244_0167 | BioFocus | N(C(=C1)C(=CC(=C(C2)C)C)C=2)(N=C2NCC(=CC3)C=CC=3OC)C(=N1)C=C2 |
| 265 | 0.0 | K02232a | 190_0144_0168 | BioFocus | N(C(=N1)C2NCC(OC3)=CC=3)(C(=C1)C(C=C1OC(F)(F)F)=CC=C1)C=CN=2 |
| 266 | 0.0 | K02233a | 190_0141_0313 | BioFocus | C(N(C1C(=CC2F)C=CC=2)C2)(=NC=1)C(=NC=2)NC(C=C1OC(C(F)F)(F)F)=CC=C1 |
| 267 | 0.0 | K02235a | 190_0141_0347 | BioFocus | C(N(C1C(=CC2C(=O)N)C=CC=2)C2)(=NC=1)C(=NC=2)NC(C=C1OC(C(F)F)(F)F)=CC=C1 |
| 268 | 0.0 | K02229a | 190_0235_0081 | BioFocus | C(N(C1C(=CC2NC(=O)C)C=CC=2)C2)(=NC=1)C(=NC=2)NC(C=C1C(F)(F)F)=CC=C1 |
| 269 | 0.0 | K02263a | 229_0236_0312 | BioFocus | N(C(=C1)C(C=C2C(F)(F)F)=CC=C2)(N=C2NCC3C=C(C(O4)=CC=3)OC4)C(=N1)C=C2 |
| 270 | 0.0 | K00525 | 294_0046_4140 | BioFocus | C(=C(C1)C(=CC2OC)C=CC=2)(N=C2NC(C=C3[Cl])=CC=C3)N(N=1)C=C2 |
| 271 | 0.0 | K00514 | 229_0254_0279 | BioFocus | N(C(=C1)C(C=C2[Cl])=CC=C2)(N=C2NC(C(C)C)CO)C(=N1)C=C2 |

| | | | | | |
|---|---|---|---|---|---|
| 272 | 0.0 | K00486 | 229_0153_0164 | BioFocus | N(C(=C1)C(=CC(=C(C2)F)[Cl])C=2)(N=C2NC(CO)CC)C(=N1)C=C2 |
| 273 | 0.0 | K00516 | 229_0254_0312 | BioFocus | N(C(=C1)C(C=C2C(F)(F)F)=CC=C2)(N=C2NC(C(C)C)CO)C(=N1)C=C2 |
| 274 | 0.0 | K00571a | AG-690/11763552 | Specs | C(=NC(=C1C(=CC=C2)C=C2)C(C=C2)=CC=C2[Br])(C(=CN2)C(=C2C2)C=CC=2)N1 |
| 275 | 0.0 | K00011 | TX-1918 | Calbiochem (EMD) | C(=C(C(=C(C1)C)O)C)C=1C=C(C(C=C1)=O)C1=O |
| 276 | 0.0 | K00520 | 229_4051_0168 | BioFocus | N(C(=C1)C(C=C2OC(F)(F)F)=CC=C2)(N=C2NCC(C3)C3)C(=N1)C=C2 |
| 277 | 0.0 | K02282a | 382_0341_0312 | BioFocus | C(C(C=C1C2C=C(C(=NC=2)N)C(=CC2)C=CC=2OCOC)=CC=C1)(F)(F)F |
| 278 | 0.0 | K02421a | 229_0236_0062 | BioFocus | N(C(=C1)C(C(=C2C3)C=CC=3)=CC=C2)(N=C2NCC3C=C(C(O4)=CC=3)OC4)C(=N1)C=C2 |
| 279 | 0.0 | K00499 | 229_0242_0160 | BioFocus | N(C(=C1)C(C=C2)=CC=C2[Cl])(N=C2NC(CC3)CCO3)C(=N1)C=C2 |
| 280 | 0.0 | K02200a | 040_0115_0164 | BioFocus | N(C(C(=CC(=C(C1)F)[Cl])C=1)=C1)=CC(=N1)NCCC(N=C1)=CC=C1 |
| 281 | 0.0 | K00577a | AM-807/12426164 | Specs | C(=C(N1)N)(SC=1NC(=CC1[Cl])C=CC=1)C(C(=CC1)C=CC=1F)=O |
| 282 | 0.0 | K02420a | 229_0131_0063 | BioFocus | N(C(=C1)C(=C(OC(=CC2)C=CC=2)C2)C=CC=2)(N=C2NCC3=CC(=C(C(=C3)OC)OC)OC)C(=N1)C=C2 |
| 283 | 0.0 | K00056 | Emodin | Calbiochem (EMD) | C1=C(C=C(C(=C1C1=O)C(C(=C1C1)C(=CC=1O)O)=O)O)C |
| 284 | 0.0 | K00203 | BIM 4 | AXXORA | C(=C(C(C(C1=CC2)=CC=2)=CN1)C(N1)=O)(C1=O)C1C(C=CC=C2)=C2NC=1 |
| 285 | 0.0 | K00755a | T5675105 | Enamine | N#CC(=CC1=CC=CO1)C1=NC(C=CC=C2)=C2S1 |
| 286 | 0.0 | K02468a | 7517783 | ChemBridge | C(NC(C(C=C1F)=CC=C1)=O)(=C(N(CC1)CCO1)C1)C=C(C(F)(F)F)C=1 |
| 287 | 0.0 | K00047c | 1-Na-PP1 | Calbiochem (EMD) | NC1=C(C(C2=C(C=CC=C3)C3=CC=C2)=NN4C(C)(C)C)C4=NC=N1 |
| 288 | 0.0 | K00473 | ST4022620 | TimTec | C(C(N1)=O)(C(=C1C1)C=CC=1)=NC(C=C1)=CC=C1C(=O)O |
| 289 | 0.0 | K00983a | AZD 7762 | SelleckChem | FC1=CC=CC(C2=CC(NC(N)=O)=C(C(N[C@@H]3CNCCC3)=O)S2)=C1 |
| 290 | 0.0 | K00752a | T0509-1692 | Enamine | N#CC(=C1CCCC1)C1=NC(C=CC=C2)=C2S1 |
| 291 | 0.0 | K00779a | Flt3 Inhib III | Calbiochem (EMD) | C(=CC=CC1C(=CN=C2NC(=CC=C(C3)OCCN(CCC4)C4)C=3)S2)C=1 |
| 292 | 0.0 | K00862a | LY 364947 | Calbiochem (EMD) | C(=CC=C(C1C2C(C(=NN3)C(=CC=CC4)N=4)=C3)N=CC=2)C=1 |
| 293 | 0.0 | K03865a | SCH-51344 | Tocris | CC1=NN=C(N3)C1=C(NCCOCCO)C2=C3C=CC(OC)=C2 |
| 294 | 0.0 | K00956a | Axitinib | SelleckChem | O=C(NC)C1=C(SC2=CC=C(C(/C=C/C4=CC=CC=N4)=NN3)C3=C2)C=CC=C1 |
| 295 | 0.0 | K00794a | T5531146 | Enamine | C(=NN1)(C(=C1C=C1)C=C1)C(NCC(OC1)=CC=1)=O |
| 296 | 0.0 | K00613a | Aminopurvalanol | Calbiochem (EMD) | N1C(=NC(=C2C=1NC1=CC(=CC(=C1)N)[Cl])N(C=N2)C(C)C)NC(C(C)C)CO |
| 297 | -0.1 | K02395a | 9046662 | ChemBridge | C(NC1=CC(=C(N2)C=C1)C=N2)(C(C=C1F)=CC=C1)=O |
| 298 | -0.1 | K00742a | F0676-0950 | Life Chemicals | N(C(=N1)C(=CC(=C(C2)OC)OC)C=2)(C(=N1)C1)N=C(SC(C(=O)C)C(=O)C)C=1 |
| 299 | -0.1 | K02659b | SKF-86002-A2 | GSK | [Cl].FC(=CC=C1C(N2)=C(N3C=2SCC3)C(C=CN=C2)=C2)C=C1 |
| 300 | -0.1 | K04080a | BIX02188 | Boehringer Ingelheim | CN(C)Cc1cccc(c1)NC(=C1C(Nc2cc(ccc12)C(N)=O)=O)c1ccccc1 |
| 301 | -0.1 | K00819a | IKK-3 Inhibitor IX | Calbiochem (EMD) | C1=C(C(=CC(=C1N1C(SC(=C2OCC(=CC=CC3)C=3S(=O)(=O)C)C#N)=C2)N=C1)OC)OC |
| 302 | -0.1 | K00761a | T5679507 | Enamine | CN1C=C(C=C(C#N)C2=NC(C=CC=C3)=C3S2)C=N1 |
| 303 | -0.1 | K00799a | T5730157 | Enamine | C(=NN1)(C(NC(=NC2)SC=2)=O)C(=C1C=C1)C=C1 |
| 304 | -0.1 | K01741a | KH-CARB9 | in-house collection | ClC(C=CC1=C2N(C)C3=C1C(C#N)C4(CCOCC4)NC3=O)=C2Cl |
| 305 | -0.1 | K00019 | 2-(Morpholin-4-yl)-benzo[h]chromen-4-one | Calbiochem (EMD) | C(=CC=CC1)(C=1C(=C(C1)C2=O)OC(=C2)N(CCOC2)C2)C=1 |
| 306 | -0.1 | K01739a | KH-CARB7 | in-house collection | ClC(C=CC1=C2N(C)C3=C1C(C#N)C4(CCCC4)NC3=O)=C2Cl |
| 307 | -0.1 | K03447a | 5919559 | ChemBridge | C1(=C(C(C(=N2)S1)=C(C(=C2N(CC1)CCO1)C1)CCC1)N)C(=O)N |
| 308 | -0.1 | K00715a | F0676-0415 | Life Chemicals | N(C(=N1)C(=CC2F)C=CC=2)(C(=N1)C1)N=C(C=1)SCC(NC(C=C1)=CC=C1NC(=O)C)=O |
| 309 | -0.1 | K00717a | F0676-0418 | Life Chemicals | N(C(=N1)C(=CC2F)C=CC=2)(C(=N1)C1)N=C(C=1)SCC(NCC(=O)OCC)=O |
| 310 | -0.1 | K00067 | ST638 | Calbiochem (EMD) | OC1=C(C=C(C=C(C(N)=O)C#N)C=C1OCC)CSC2=CC=CC=C2 |
| 311 | -0.1 | K00725a | F0676-0464 | Life Chemicals | N(C(=N1)C(=CC2F)C=CC=2)(C(=N1)C1)N=C(SC(C(=O)OC)C)C=1 |
| 312 | -0.1 | K02615b | JAK3 Inhibitor IV (no. 1367) | Tocris | C1C=CC(=C2C=1)C=CC(=C2)C(CCN(CC1=CC=CC=C1)C(C)C)=O.[H][Cl] |
| 313 | -0.1 | K02466a | 7497813 | ChemBridge | C(NC(C(=CN1)C=CC=1)=O)(=C(N(CC1)CCO1)C1)C=C(C(F)(F)F)C=1 |
| 314 | -0.1 | K02845b | SB-223133 | GSK | NC(N=CC=C1C(=C2C(C=CC(F)=C3)=C3)N(C=N2)C(CCC(O)C2)C2)=N1 |
| 315 | -0.1 | K00064 | Tyrphostin B42 | SelleckChem | C1=CC=C(C=C1)CNC(=O)C(=CC2=CC(=C(C=C2)O)O)C#N |
| 316 | -0.1 | K00738a | F0676-0939 | Life Chemicals | N(C(=N1)C(=CC(=C(C2)OC)OC)C=2)(C(=N1)C1)N=C(SC(C(=O)OC)CC)C=1 |
| 317 | -0.1 | K00069 | Cdk2 Inhibitor II | Calbiochem (EMD) | C1=CC(=CC=C1NNC2=C3C=C(C=CC3=NC2=O)Br)S(=O)(=O)N |
| 318 | -0.1 | K02396a | 9048213 | ChemBridge | C(NC1=CC(=C(N2)C=C1)C=N2)(C(C=C(C(=C1)OC)OC)=C1)=O |
| 319 | -0.1 | K00003 | H-1152 | AXXORA | N(CC(N1S(C(=C(C(=CC2)C3)C(=CN=3)C)C=2)(=O)=O)C)CCC1.[Cl].[Cl] |
| 320 | -0.2 | K00231 | (-)-Arctigenin | Calbiochem (EMD) | C1C(=CC(=C(C=1)OC)OC)CC(CO1)C(C1=O)CC1C=CC(=C(C=1)OC)OC |
| 321 | -0.2 | K00718a | F0676-0420 | Life Chemicals | N(C(=N1)C(=CC2F)C=CC=2)(C(=N1)C1)N=C(C=1)SCC(NCC(C=C1)=CC=C1F)=O |
| 322 | -0.2 | K00676a | F2563-0290 | Life Chemicals | COC(c1ccccc1NC(CSc1nnc(c2ccccc2)n1n1cccc1)=O)=O |
| 323 | -0.2 | K00740a | F0676-0945 | Life Chemicals | N(C(=N1)C(=CC(=C(C2)OC)OC)C=2)(C(=N1)C1)N=C(SC(C(=O)O)CC)C=1 |
| 324 | -0.2 | K00554a | 5174681 | ChemBridge | N(C(N1)=O)(C(C(C1=O)=NNC(C=C1)=CC=C1[Br])=O)C |
| 325 | -0.2 | K00222 | Compound 52 | Calbiochem (EMD) | N(C(=C(C(=NC1NCCO)N2C(C)C)N=C2)N=1)C(=CC(=CC1)[Cl])C=1 |
| 326 | -0.2 | K00005 | (Z)-3-[4-(Dimethylamino)benzylidenyl]indolin-2-one | Calbiochem (EMD) | C1C=CC=C(C=1N1)C(C1=O)=CC(=CC=C(C1)N(C)C)C=1 |
| 327 | -0.2 | K04055a | SK1-I (BML-EI411-0005) | Enzo | CCCCCC1=CC=C(/C=C/[C@H](O)[C@H](NC)CO)C=C1.Cl |
| 328 | -0.2 | K00746a | VEGFR 2/3 Inhibitor | Calbiochem (EMD) | C(C=C1)=CC(=C1N1)C(=C1)C(=C(C1=O)C(=C2)C=C(C(=C2OC)OC)OC)C(N1)=O |
| 329 | -0.2 | K02399a | 9051276 | ChemBridge | N1NC(=C(C=C2NC(=O)CC(C=C3)=CC=C3F)C=1)C=C2 |
| 330 | -0.3 | K03183a | TCS 2002 | Tocris | C1=CC(=CC(=C1O1)C(=C1)C(=CC=C(C1)S(C)=O)C=1)C(=NN=C1C)O1 |
| 331 | -0.3 | K00276 | BAS 00382581 | Asinex | S(C(=CC=C1N=C(C(N2)=O)C(=C2C2)C=CC=2)C=C1)(=O)(=O)N |
| 332 | -0.3 | K00753a | T0507-3274 | Enamine | N#CC(=C1CCCN1)C1=NC(C=CC=C2)=C2S1 |
| 333 | -0.3 | K00785a | 7818069 | ChemBridge | N(C(=C1F)N(CC2)CCO2)=C(N=C1)NC(=CC1)C=CC=1OC |
| 334 | -0.3 | K03955a | Pazopanib | LC Labs | O=S(=O)(N)c1c(ccc(c1)Nc2nccc(n2)N(c4ccc3c(nn(c3C)C)c4)C)C |
| 335 | -0.3 | K02499a | 7522605 | ChemBridge | C(OC1C(=CC2[Cl])C=CC=2)(C(NC(=C(N(CC2)CCO2)C2)C=C(C(F)(F)F)C=2)=O)=CC=1 |
| 336 | -0.3 | K00758a | T0501-4046 | Enamine | CN(C=CC=C1)C1=C(C#N)C1=NC(C=CC=C2)=C2S1 |
| 337 | -0.3 | K00747a | PD173074 | Calbiochem (EMD) | N(C(=N1)NCCCCN(CC)CC)=CC(=C12)C=C(C(=N2)NC(=O)NC(C)(C)C)C(=CC(=C1)OC)C=C1OC |
| 338 | -0.3 | K00737a | F0676-0937 | Life Chemicals | N(C(=N1)C(=CC(=C(C2)OC)OC)C=2)(C(=N1)C1)N=C(SC(C(=O)OCC)C)C=1 |
| 339 | -0.3 | K00793a | T5302202 | Enamine | C(=NN1)(C(=C1C=C1)C=C1)C(NCC(OC1)CC1)=O |
| 340 | -0.3 | K00733a | F0676-0910 | Life Chemicals | N(C(=N1)C(=CC(=C(C2)OC)OC)C=2)(C(=N1)C1)N=C(C=1)SCC(NC(=NO1)C=C1C)=O |

| | | | | | |
|---|---|---|---|---|---|
| 341 | -0.3 | K00736a | F0676-0936 | Life Chemicals | N(C(=N1)C(=CC(=C(C2)OC)OC)C=2)(C(=N1)C1)N=C(C=1)SCC(=O)OCC |
| 342 | -0.3 | K00744a | PI 3-K Inhibitor IV | Calbiochem (EMD) | C(S1)=CC(=C1C(=NC1C(C=C(C2)O)=CC=2)N2CCOCC2)N=1 |
| 343 | -0.3 | K03964a | Bosutinib isomer | LC Labs | CN(C)Cc1cccc(c1)NC(=C1C(Nc2cc(ccc12)C(N(C)C)=O)=O)c1ccccc1 |
| 344 | -0.3 | K00727a | F0676-0840 | Life Chemicals | N(C(=N1)C(=CC(=C(C2)OC)OC)C=2)(C(=N1)C1)N=C(C=1)SCC(N(CC1)CCO1)=O |
| 345 | -0.3 | K00999a | WZ-4-49-8 | in-house collection | O=S(C1=CC=CC=C1NC2=C3C(NN=C3)=NC(NC4=CC=C(N5CCC(N6CCCCC6)CC5)C=C4OCC)=N2)(C(C)C)=O |
| 346 | -0.4 | K00745a | BAY 61-3606 | SelleckChem | O=C(C1=CC=CN=C1NC2=NC(C3=CC=C(OC)C(OC)=C3)=CC4=NC=CN24)N |
| 347 | -0.4 | K00068 | JNK Inhibitor II | Calbiochem (EMD) | C1=CC=CC(=C1C1=O)C(C(=C1C1)C(=CC=1)N1)=N1 |
| 348 | -0.4 | K00759a | T5674810 | Enamine | CN(C=CC1)C=1C=C(C#N)C1=NC(C=CC=C2)=C2S1 |
| 349 | -0.5 | K04081a | BIX01289 | Boehringer Ingelheim | CN(C)Cc1cccc(c1)NC(=C1C(Nc2cc(ccc12)C(N(C)C)=O)=O)c1ccccc1 |
| 350 | -0.5 | K03740a | GW656282X | GSK | Cc1ccc(cc1Nc1ccnc(Nc2cccc(c2)C(N)=O)n1)O |
| 351 | -0.5 | K00570b | SB 239063 | Sigma Aldrich | C(=C(N=C1)C2=CC=C(C=C2)F)(N1C1CCC(CC1)O)C1=CC=NC(=N1)OC |
| 352 | -0.5 | K00298 | 1910-5183 | ChemDiv | C(C(=C1C2)C=C([N+]([O-])=O)C=2)(C(N1)=O)=NC(C=C1)=CC=C1C(=O)OCC |
| 353 | -0.5 | K00472 | ST2029251 | TimTec | C(=C(S1)N2)(C(=C1C1)CC(C(CC)(C)C)C1)C(NC=2)=O |
| 354 | -0.5 | K00795a | T5543714 | Enamine | C(=NN1)(C(=C1C=C1)C=C1)C(NCC(F)(F)F)=O |
| 355 | -0.6 | K00774a | T0505-3534 | Enamine | O=C(C1CC1)C(=C1NC(C=CC=C2)=C2S1)C#N |
| 356 | -0.6 | K00648a | 5228205 | ChemBridge | C(=C(C1=O)C2O)(C(C(=C1C1)C=CC=1)=O)C(=CC=2)NCCC(=O)OC |
| 357 | -0.7 | K00728a | F0676-0841 | Life Chemicals | N(C(=N1)C(=CC(=C(C2)OC)OC)C=2)(C(=N1)C1)N=C(C=1)SCC(N(CC1)CCC1C)=O |
| 358 | -0.8 | K02721a | AG-227/37195019 | Specs | N(=C(C1C(C=C2)=CC=C2)N(CC)CC)C(=NN=1)C(=CC1)C=CC=1[Cl] |
| 359 | -1.0 | K03953a | Temsirolimus | Sigma Aldrich | O=C([C@H](C)\C=C([C@@H](O)[C@@H](OC)C([C@@H]4C)=O)/C)C[C@]([C@H](C)C[C@H]2C[C@@H](OC)[C@H](OC(C(CO)(C)CO)=O)CC2)([H])OC([C@@]1([H])N(C(C([C@@]3(O)[C@H](C)CC[C@](C[C@@H](/C(C)=C/C=C/C=C/[C@H](C4)C)OC)([H])O3)=O)=O)CCCC1)=O |
| 360 | -1.7 | K03886a | FTY720 | Sigma Aldrich | CCCCCCCCc1ccc(CCC(CO)(CO)N)cc1 |

**Supplementary method.**

**Sources and chemical syntheses of inhibitors**

| **Inhibitors** | **Sources** |
|---|---|
| ibrutinib | SelleckChem (Cat no. S2680) |
| OTSSP167 | SelleckChem (Cat no. S7159) |
| CPT1-70-1 | Reference# (Tan et al., 2017) and patent WO 2016130920 |
| K00007 | Calbiochem (Cat no. 175580) |
| ASC69 | MolPort (Cat no. MolPort-039-333-128);<br><br>Synthesis also described in patent WO2010031056A2 |
| HG-6-64-01 | Reference# (Miao et al., 2015),(Tan et al., 2015) |
| HG-6-71-01 | Reference# (Kim et al., 2013) |
| HYJ-2-002-1 | Reference# (Tan et al., 2015) |
| XMD15-46 | Reference# (Deng et al., 2010) |
| TL10-105 | Synthesis described below |
| SB1-G-23 | Synthesis described below |

### Chemical syntheses of TL10-105 and SB1-G-23

Unless otherwise noted, reagents and solvents were obtained from commercial suppliers and were used without further purification. 1H NMR spectra were recorded on 600 or 400 MHz (Varian AS600 or Bruker A400), and chemical shifts are reported in parts per million (ppm, $\delta$) downfield from tetramethylsilane (TMS). Coupling constants (*J*) are reported in Hz. Spin multiplicities are described as s (singlet), br (broad singlet), d (doublet), t (triplet), q (quartet), and m (multiplet). Mass spectra were obtained on a Waters Micromass ZQ instrument. Preparative HPLC was performed on a Waters Sunfire C18 column (19 x 50 mm, 5µM) using a gradient of 15-95% methanol in water containing 0.05% trifluoroacetic acid (TFA) over 22 min (28 min run time) at a flow rate of 20 mL/min. Purities of assayed compounds were in all cases greater than 95%, as determined by reverse-phase HPLC analysis.

### Synthesis of (R)-3-((2-((1-acryloylpyrrolidin-3-yl)amino)pyrimidin-4-yl)oxy)-N-(4-((4-ethylpiperazin-1-yl)methyl)-3-(trifluoromethyl)phenyl)-4-methylbenzamide (SB1-G-23)

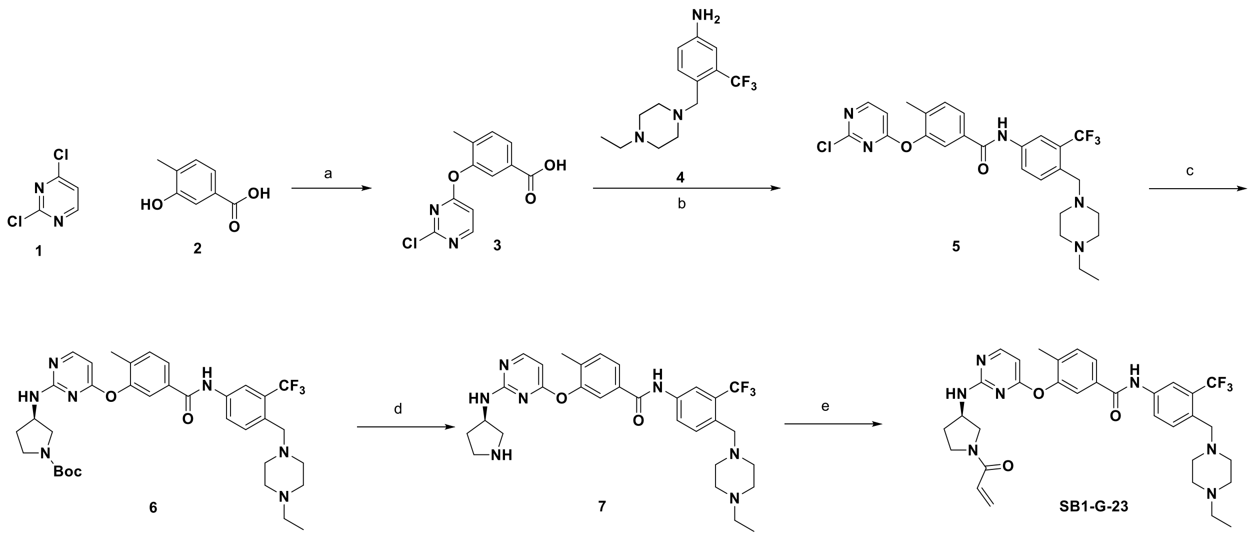


Reagents and conditions: (a) $Cs_2CO_3$, *i*-PrOH, 55 °C, 3h, 87%; (b) oxalyl dichloride, *N*,*N*-dimethylformamide, $CH_2Cl_2$, rt, 2h;then diisopropylethylamine, $CH_2Cl_2$, rt, overnight, 90%; (c) tert-butyl (R)-3-aminopyrrolidine-1-carboxylate, diisopropylethylamine, *n*-butanol, 110 °C, 59%; (d) TFA, $CH_2Cl_2$, 94%; (e) acryloyl chloride, diisopropylethylamine, $CH_3CN$, rt, 4 h, 17%.

### 3-(2-chloropyrimidin-4-yloxy)-4-methylbenzoic acid (3)

To a solution of 2,4-dichloropyrimidine (**1**) (10 g, 67.13 mmol), 3-hydroxy-4-methylbenzoic acid (**2**) (10.22 g, 67.13 mmol) and $Cs_2CO_3$ (43.75 g, 134.26 mmol) in *i*-PrOH (100 mL) was stirred at 55°C for 3h. The solvent was removed under reduced pressure and water (100 mL) was added. Adjust PH of the mixture to one with 1N HCl. Filtered and dried by acetone to give **3** (15.5 g, yield: 87 %) as a white solid. LCMS (ESI) *m/z*: 265 $[M+H]^+$.

**3-(2-chloropyrimidin-4-yloxy)-*N*-(4-((4-ethylpiperazin-1-yl)methyl)-3-(trifluoromethyl)phenyl)-4-methylbenzamide (5)**

To a solution of 3-(2-chloropyrimidin-4-yloxy)-4-methylbenzoic acid (**3**) (6.5 g, 24.56 mmol) in $CH_2Cl_2$ (100 mL) was added *N*,*N*-dimethylformamide (DMF) (1 mL) and oxalyl dichloride (9.36 g, 73.68 mmol). The mixture was stirred at r.t for 2h. The solvent was removed under reduced pressure and $CH_2Cl_2$ (100 mL) was added. The mixture was added diisopropylethylamine (DIEA) (6.39 g, 49.44 mmol) and 4-((4-ethylpiperazin-1-yl)methyl)-3-(trifluoromethyl)aniline (**4**) (7.1 g, 24.72 mmol) and stirred at r.t for 3h. The solvent was removed under reduced pressure and the product was purified by flash column ($MeOH/CH_2Cl_2$ = 5% ~ 15%) to give **5** (12 g, yield: 90 %) as a brown solid. LCMS (ESI) *m/z*: 534 $[M + H]^+$.

**tert-butyl (R)-3-((4-(5-((4-((4-ethylpiperazin-1-yl)methyl)-3-(trifluoromethyl)phenyl)carbamoyl)-2-methylphenoxy)pyrimidin-2-yl)amino)pyrrolidine-1-carboxylate (6)**

The mixture of **5** (600 mg, 1.12 mmol), tert-butyl (R)-3-aminopyrrolidine-1-carboxylate (550 mg, 2.95 mmol), DIEA (2 mL) in *n*-BuOH (15 mL) was stirred at 110°C overnight, after the reaction was completed the mixture was concentrated to remove the solvent, added saturated $Na_2CO_3$ solution (30 mL),then extracted with ethyl acetate (100 mL × 3), the organic phase was washed with brine (50 mL × 2), dried over $Na_2SO_4$, filtered, concentrated and purified by silica gel ($MeOH/CH_2Cl_2$ = 1/10) to obtain **6** (light brown solid, 450 mg, yield 59%). LCMS (ESI) *m/z*: 684 $[M + H]^+$.

**(R)-N-(4-((4-ethylpiperazin-1-yl)methyl)-3-(trifluoromethyl)phenyl)-4-methyl-3-((2-(pyrrolidin-3-ylamino)pyrimidin-4-yl)oxy)benzamide (7)**

The mixture of **6** (180 mg, 0.263 mmol), TFA (2 mL) in $CH_2Cl_2$ (5 mL) was stirred at r.t. for 4 h, concentrated to remove the solvent to get **7** (light brown solid, 120 mg, yield 94%). LCMS (ESI) *m/z*: 584 $[M + H]^+$.

**(R)-3-((2-((1-acryloylpyrrolidin-3-yl)amino)pyrimidin-4-yl)oxy)-N-(4-((4-ethylpiperazin-1-yl)methyl)-3-(trifluoromethyl)phenyl)-4-methylbenzamide (SB1-G-23)**

To the mixture of **7** (60 mg, 0.103 mmol), DIPEA (1 mL) in $CH_3CN$ (10 mL) was added acryloyl chloride (12 mg, 0.133 mmol), then stirred at r.t for 4 h, concentrated to remove the solvent, the residue was purified by prep-HPLC (C18 column, $CH_3CN/H_2O$, containing 0.05%$NH_4HCO_3$) to obtain **SB1-G-23** (off-white solid, 11 mg, yield 17%). LCMS (ESI) *m/z*: 638 $[M + H]^+$; $^1H$ NMR (400 MHz, DMSO-$d_6$) δ 10.24 (s, 1H), 8.22 (d, *J* = 7.6 Hz, 1H), 8.15 (s, 1H), 8.01 (d, *J* = 10.8 Hz, 1H), 7.83 (d, *J* = 11.2 Hz, 2H), 7.78 (s, 1H), 7.68 (d, *J* = 12.0 Hz, 1H), 7.47 (d, *J* = 10.0 Hz, 1 H), 7.20 (bs,

1H), 6.00-6.51 (m, 3H), 5.50-5.61 (m, 1H), 4.16 (bs, 1H), 3.28-3.65 (m, 7H), 2.30-2.40 (m, 8H) 2.28 (q, $J$ = 9.6 Hz, 2H), 2.20 (s, 3H), 0.98 (t, $J$ = 9.6 Hz, 3H).

**Synthesis of (S)-3-((2-((1-acryloylpyrrolidin-3-yl)amino)pyrimidin-4-yl)oxy)-N-(4-((4-ethylpiperazin-1-yl)methyl)-3-(trifluoromethyl)phenyl)-4-methylbenzamide (TL10-105)**

TL10-105 was synthesized by similar procedure as synthesis of SB1-G-23 with tert-butyl (S)-3-aminopyrrolidine-1-carboxylate as starting material in step c. LCMS (ESI) *m/z*: 638 $[M + H]^+$; $^1H$ NMR (600 MHz, DMSO-$d_6$, TFA salt) δ 10.41 (s, 1H), 9.34 (br, 1H), 8.25 (d, $J$ = 8.4 Hz, 1H), 8.20 (s, 1H), 8.10 (d, $J$ = 7.8 Hz, 1H), 7.85 (d, $J$ = 7.8 Hz, 2H), 7.80 (s, 1H), 7.71 (d, $J$ = 8.4 Hz, 1H), 7.50 (d, $J$ = 8.4 Hz, 1H), 6.52 (dd, $J$ = 16.8, 10.8 Hz,1H), 6.33 (d, $J$ = 6.0 Hz, 1H), 6.28 (br, 1H), 5.63 (d, $J$ = 6.0 Hz, 1H), 4.33 (m, 7H), 3.69 (s, 2H), 3.46 (m, 2H), 3.15 (m, 2H), 2.88–3.04 (m, 4H) 2.41 (q, $J$ = 7.2 Hz, 2H), 2.20 (s, 3H), 1.22 (t, $J$ = 7.2 Hz, 3H).

### References


Deng, X., Lim, S.M., Zhang, J., and Gray, N.S. (2010). Broad spectrum alkynyl inhibitors of T315I Bcr-Abl. Bioorg Med Chem Lett *20*, 4196-4200.

Kim, H.G., Tan, L., Weisberg, E.L., Liu, F., Canning, P., Choi, H.G., Ezell, S.A., Wu, H., Zhao, Z., Wang, J.*, et al.* (2013). Discovery of a potent and selective DDR1 receptor tyrosine kinase inhibitor. ACS Chem Biol *8*, 2145-2150.

Miao, B., Ji, Z., Tan, L., Taylor, M., Zhang, J., Choi, H.G., Frederick, D.T., Kumar, R., Wargo, J.A., Flaherty, K.T.*, et al.* (2015). EPHA2 is a mediator of vemurafenib resistance and a novel therapeutic target in melanoma. Cancer Discov *5*, 274-287.

Tan, L., Gurbani, D., Weisberg, E.L., Hunter, J.C., Li, L., Jones, D.S., Ficarro, S.B., Mowafy, S., Tam, C.P., Rao, S.*, et al.* (2017). Structure-guided development of covalent TAK1 inhibitors. Bioorg Med Chem *25*, 838-846.

Tan, L., Nomanbhoy, T., Gurbani, D., Patricelli, M., Hunter, J., Geng, J., Herhaus, L., Zhang, J., Pauls, E., Ham, Y.*, et al.* (2015). Discovery of type II inhibitors of TGFbeta-activated kinase 1 (TAK1) and mitogen-activated protein kinase kinase kinase kinase 2 (MAP4K2). J Med Chem *58*, 183-196.

### Supplementary table 3

| KINOMEscan<br><br>Kinases | TL10-105 (1µM)<br>% control<br><br>This study | HYJ-2-002-1 (1µM)<br>% control<br><br>This study | CPT-1-70-1 (1µM)<br>% control<br><br>This study | ibrutinib (1µM)<br><br>% control http://lincs.hms.harvard.edu/ | OTSSP167 (10µM)<br><br>% control http://lincs.hms.harvard.edu/ | XMD15-46 (10µM)<br><br>% control from Deng et al., 2010 | HG-6-71-01 (1µM)<br><br>% control from Kim et al., 2013 | HG6-64-1 (1µM)<br><br>% control from Tan et al., 2015 | HG6-64-1 (10µM)<br><br>% control from Tan et al., 2015 |
|---|---|---|---|---|---|---|---|---|---|
| MKK7 | 0.9 | 8.2 | 8.1 | 1.6 | 0.3 | 5.4 | 20 | 39 | 1.4 |
| AAK1 | 82 | 100 | 18 | 92 | 2.3 | 59 | 100 | 67 | 26 |
| ABL1(E255K)-phosphorylated | 1.6 | 0.55 | 90 | 65 | 0.9 | 0.2 | 1.3 | 0.85 | 1 |
| ABL1(F317I)-nonphosphorylated | 1.3 | 0 | 100 | 30 | 14 | 8.8 | 2.4 | 19 | 0.9 |
| ABL1(F317I)-phosphorylated | 7.6 | 34 | 100 | 39 | 3.2 | 4.9 | 32 | 22 | 19 |
| ABL1(F317L)-nonphosphorylated | 1.6 | 0 | 100 | 16 | 94 | 0 | 4.2 | 0 | 0 |
| ABL1(F317L)-phosphorylated | 3.3 | 2 | 99 | 48 | 7.3 | 0.85 | 0.8 | 3.4 | 2.2 |
| ABL1(H396P)-nonphosphorylated | 0 | 0 | 7.2 | 12 | 5.6 | 0.05 | 0.8 | 0 | 0.1 |
| ABL1(H396P)-phosphorylated | 2.3 | 0.1 | 100 | 87 | 0.2 | 0.05 | 0.45 | 0.8 | 0.1 |
| ABL1(M351T)-phosphorylated | 2.9 | 0.85 | 86 | 74 | 1.2 | 1.1 | 0.4 | 4.4 | 2.2 |
| ABL1(Q252H)-nonphosphorylated | 0.8 | 0.05 | 52 | 5.1 | 15 | 1.6 | 2.6 | 1.4 | 0.1 |
| ABL1(Q252H)-phosphorylated | 2.4 | 1 | 100 | 76 | 0.9 | 0.2 | 1.6 | 1.2 | 1.2 |
| ABL1(T315I)-nonphosphorylated | 5 | 9 | 46 | 96 | 12 | 2.4 | 2.5 | 11 | 0 |
| ABL1(T315I)-phosphorylated | 62 | 0.75 | 19 | 70 | 1.5 | 0.9 | 1.2 | 5.4 | 1.8 |
| ABL1(Y253F)-phosphorylated | 1 | 0.7 | 100 | 62 | 0.6 | 0.2 | 1.1 | 0.75 | 0.3 |
| ABL1-nonphosphorylated | 1 | 0.05 | 25 | 7.2 | 2.6 | 0.1 | 2.8 | 0 | 0 |
| ABL1-phosphorylated | 2 | 0.1 | 51 | 54 | 0.2 | 0.05 | 0.45 | 0.7 | 0.1 |
| ABL2 | 5 | 0 | 60 | 38 | 0.2 | 0 | 0.55 | 0.7 | 0 |
| ACVR1 | 100 | 100 | 100 | 100 | 0.7 | 100 | 100 | 100 | 100 |
| ACVR1B | 92 | 100 | 100 | 100 | 31 | 63 | 100 | 95 | 98 |
| ACVR2A | 94 | 100 | 100 | 97 | 100 | 100 | 100 | 100 | 100 |
| ACVR2B | 93 | 100 | 100 | 90 | 70 | 23 | 86 | 91 | 62 |
| ACVRL1 | 85 | 100 | 100 | 100 | 38 | 100 | 100 | 100 | 100 |
| ADCK3 | 100 | 100 | 100 | 100 | 0.1 | 100 | 100 | 100 | 100 |
| ADCK4 | 96 | 100 | 91 | 96 | 85 | 82 | 100 | 81 | 100 |
| AKT1 | 100 | 100 | 100 | 100 | 3.7 | 41 | 100 | 100 | 100 |
| AKT2 | 98 | 95 | 100 | 75 | 1.2 | 4.6 | 93 | 86 | 48 |

| | | | | | | | | | |
|---|---|---|---|---|---|---|---|---|---|
| AKT3 | 94 | 100 | 100 | 100 | 48 | 82 | 100 | 100 | 100 |
| ALK | 82 | 75 | 38 | 51 | 6.6 | 52 | 44 | 80 | 51 |
| ALK(C1156Y) | 100 | n/a | 52 | 74 | 8 | n/a | n/a | n/a | n/a |
| ALK(L1196M) | 78 | n/a | 100 | 82 | 4.8 | n/a | n/a | n/a | n/a |
| AMPK-alpha1 | 92 | 87 | 32 | 75 | 0.5 | 0.8 | 89 | 25 | 20 |
| AMPK-alpha2 | 86 | 100 | 41 | 100 | 1.4 | 7.2 | 97 | 61 | 39 |
| ANKK1 | 100 | 9.4 | 93 | 71 | 2.3 | 7.1 | 5.2 | 6 | 5.7 |
| NUAK1 | 83 | 65 | 9.3 | 100 | 2.5 | 2.5 | 85 | 100 | 100 |
| ASK1 | 94 | 96 | 100 | 87 | 1 | 24 | 91 | 100 | 35 |
| ASK2 | 87 | 100 | 100 | 100 | 0.4 | 11 | 16 | 87 | 64 |
| AURKA | 92 | 100 | 37 | 49 | 0 | 97 | 86 | 99 | 97 |
| AURKB | 98 | 43 | 61 | 91 | 0.1 | 4.6 | 54 | 65 | 16 |
| AURKC | 100 | 80 | 51 | 77 | 34 | 11 | 71 | 34 | 1.5 |
| AXL | 78 | 21 | 4.8 | 65 | 16 | 0.1 | 60 | 55 | 4.6 |
| BIKE | 96 | 96 | 18 | 84 | 3.4 | 30 | 88 | 25 | 3 |
| BLK | 0.5 | 0.1 | 1.6 | 0.1 | 0.7 | 0 | 0 | 0.25 | 0.2 |
| BMPR1A | 96 | 100 | 100 | 100 | 46 | 53 | 100 | 100 | 100 |
| BMPR1B | 93 | 100 | 100 | 75 | 0.1 | 72 | 96 | 95 | 76 |
| BMPR2 | 100 | 98 | 100 | 89 | 0.8 | 61 | 84 | 87 | 14 |
| BMX | 54 | 8.7 | 9.2 | 14 | 2.6 | 2.6 | 5 | 1.6 | 6.6 |
| BRAF | 82 | 3.7 | 87 | 53 | 3.6 | 0.45 | 11 | 17 | 0.6 |
| BRAF(V600E) | 41 | 1.2 | 100 | 50 | 1.5 | 0.1 | 4 | 7.4 | 0.1 |
| BRK | 100 | n/a | n/a | 1 | 42 | n/a | n/a | 64 | 9.6 |
| BRSK1 | 100 | 97 | 100 | 100 | 35 | 100 | 100 | 83 | 100 |
| BRSK2 | 100 | 91 | 97 | 76 | 24 | 90 | 62 | 93 | 100 |
| BTK | 98 | 8.1 | 7 | 0 | 7.6 | 1.2 | 8.8 | 10 | 0.3 |
| BUB1 | 88 | n/a | 85 | 100 | 13 | n/a | 100 | n/a | n/a |
| CAMK1 | 29 | 80 | 57 | 77 | 100 | 20 | 95 | 94 | 30 |
| CAMK1D | 28 | 94 | 56 | 84 | 60 | 68 | 94 | 100 | 100 |
| CAMK1G | 60 | 94 | 50 | 73 | 23 | 92 | 87 | 96 | 100 |
| CAMK2A | 95 | 100 | 95 | 100 | 81 | 100 | 83 | 92 | 100 |
| CAMK2B | 100 | 74 | 100 | 95 | 100 | 100 | 93 | 93 | 100 |
| CAMK2D | 88 | 100 | 100 | 79 | 1.6 | 100 | 100 | 100 | 100 |
| CAMK2G | 89 | 100 | 100 | 81 | 3 | 92 | 100 | 98 | 100 |
| CAMK4 | 73 | 100 | 100 | 100 | 1 | 100 | 100 | 100 | 100 |
| CAMKK1 | 81 | 100 | 84 | 90 | 0.7 | 69 | 100 | 96 | 100 |
| CAMKK2 | 69 | 100 | 68 | 73 | 3.5 | 27 | 96 | 100 | 100 |
| CASK | 81 | 100 | 100 | 59 | 25 | 100 | 72 | 96 | 77 |
| CDC2L1 | 36 | 88 | 100 | 96 | 0.6 | 0 | 98 | 61 | 8.5 |
| CDC2L2 | 37 | 76 | 100 | 81 | 3.4 | 0 | 78 | 70 | 5.4 |
| CDC2L5 | 65 | 100 | 91 | 89 | 52 | 13 | 94 | 100 | 100 |
| CDK11 | 0 | 14 | 100 | 88 | 0.9 | 4.2 | 32 | 27 | 4.2 |
| CDK2 | 77 | 100 | 100 | 100 | 2.8 | 13 | 100 | 90 | 96 |

| | | | | | | | | | |
|---|---|---|---|---|---|---|---|---|---|
| CDK3 | 56 | 99 | 100 | 95 | 1.4 | 32 | 95 | 100 | 100 |
| CDK4-cyclinD1 | 95 | 80 | 81 | 96 | 10 | 29 | 100 | 100 | 74 |
| CDK4-cyclinD3 | 100 | 100 | 100 | 84 | 0.7 | 65 | 72 | 77 | 73 |
| CDK5 | 72 | 100 | 100 | 90 | 1.9 | 24 | 100 | 100 | 100 |
| CDK7 | 49 | 44 | 100 | 86 | 0.3 | 4 | 97 | 72 | 41 |
| CDK8 | 2.8 | 9.2 | 100 | 76 | 1.2 | 18 | 32 | 25 | 21 |
| CDK9 | 72 | 100 | 90 | 90 | 7.5 | 31 | 100 | 92 | 96 |
| CDKL1 | 77 | 94 | 66 | 72 | 2.6 | 89 | 72 | 100 | 61 |
| CDKL2 | 19 | 2.3 | 11 | 100 | 4 | 4.8 | 2 | 4.2 | 1.6 |
| CDKL3 | 24 | 34 | 100 | 100 | 0.2 | 1.9 | 71 | 50 | 4 |
| CDKL5 | 74 | 100 | 100 | 97 | 0.9 | 28 | 58 | 100 | 78 |
| CHEK1 | 100 | 100 | 92 | 100 | 37 | 88 | 100 | 100 | 28 |
| CHEK2 | 94 | 63 | 38 | 96 | 100 | 22 | 100 | 82 | 66 |
| CIT | 61 | 6.7 | 26 | 65 | 3.1 | 0 | 18 | 4.6 | 0 |
| CLK1 | 85 | 57 | 5.8 | 100 | 15 | 6.8 | 83 | 59 | 14 |
| CLK2 | 98 | 85 | 22 | 99 | 100 | 100 | 90 | 88 | 71 |
| CLK3 | 96 | 88 | 82 | 90 | 21 | 94 | 100 | 90 | 100 |
| CLK4 | 82 | 69 | 6.5 | 97 | 3.4 | 32 | 86 | 98 | 100 |
| CSF1R | 2.8 | 0.45 | 12 | 75 | 11 | 0 | 0.55 | 0.75 | 0 |
| CSF1R-autoinhibited | 95 | n/a | 1.4 | 58 | 3.2 | n/a | 64 | n/a | n/a |
| CSK | 58 | 0.8 | 93 | 0.5 | 23 | 0.1 | 1 | 2 | 0.8 |
| CSNK1A1 | 88 | 100 | 55 | 87 | 2.8 | 89 | 66 | 93 | 87 |
| CSNK1A1L | 96 | 100 | 80 | 99 | 1 | 75 | 100 | 95 | 100 |
| CSNK1D | 100 | 73 | 43 | 97 | 9.4 | 22 | 89 | 84 | 38 |
| CSNK1E | 90 | 75 | 18 | 42 | 0.2 | 26 | 95 | 80 | 45 |
| CSNK1G1 | 91 | 93 | 18 | 86 | 78 | 100 | 89 | 87 | 98 |
| CSNK1G2 | 97 | 45 | 0.7 | 92 | 2.2 | 14 | 90 | 94 | 39 |
| CSNK1G3 | 92 | 65 | 2.6 | 100 | 3 | 22 | 99 | 71 | 48 |
| CSNK2A1 | 100 | 94 | 35 | 57 | 0.9 | 100 | 31 | 100 | 90 |
| CSNK2A2 | 93 | 100 | 28 | 52 | 0.1 | 78 | 69 | 77 | 95 |
| CTK | 74 | 91 | 100 | 40 | 22 | 45 | 8 | 60 | 100 |
| DAPK1 | 90 | 98 | 86 | 100 | 45 | 94 | 82 | 77 | 100 |
| DAPK2 | 99 | 92 | 88 | 92 | 100 | 100 | 69 | 99 | 100 |
| DAPK3 | 100 | 100 | 59 | 91 | 100 | 96 | 96 | 87 | 99 |
| DCAMKL1 | 99 | 100 | 100 | 62 | 4.5 | 100 | 75 | 100 | 93 |
| DCAMKL2 | 75 | 100 | 92 | 90 | 22 | 30 | 100 | 87 | 100 |
| DCAMKL3 | 100 | 84 | 56 | 82 | 0.4 | 100 | 100 | 100 | 3.2 |
| DDR1 | 0.4 | 0 | 79 | 90 | 2.6 | 0 | 0.15 | 0.05 | 0.2 |
| DDR2 | 12 | 0.05 | 100 | 83 | 22 | 2.8 | 9.9 | 0.2 | 9 |
| DLK | 90 | 86 | 78 | 93 | 85 | 3.8 | 100 | 74 | 22 |
| DMPK | 98 | 100 | 41 | 100 | 7.2 | 27 | 96 | 100 | 57 |
| DMPK2 | 96 | 92 | 67 | 97 | 1.2 | 35 | 100 | 76 | 16 |

| | | | | | | | | | |
|---|---|---|---|---|---|---|---|---|---|
| DRAK1 | 88 | 82 | 14 | 100 | 9.6 | 24 | 100 | 52 | 1.3 |
| DRAK2 | 98 | 57 | 90 | 100 | 100 | 12 | 92 | 16 | 1 |
| DYRK1A | 100 | 62 | 100 | 83 | 0 | 100 | 74 | 100 | 63 |
| DYRK1B | 91 | 76 | 79 | 93 | 5.8 | 15 | 82 | 73 | 27 |
| DYRK2 | 85 | 96 | 100 | 82 | 0.4 | 51 | 82 | 68 | 78 |
| EGFR | 68 | 5 | 63 | 1.8 | 13 | 0.3 | 4 | 7.6 | 0.4 |
| EGFR(E746-A750del) | 66 | 1 | 17 | 5.8 | 8.4 | 0.75 | 4.4 | 0.7 | 3.8 |
| EGFR(G719C) | 59 | 0.75 | 8 | 1.2 | 36 | 0 | 1.3 | 0.45 | 0.1 |
| EGFR(G719S) | 81 | 1.8 | 34 | 0.9 | 46 | 0.15 | 1.8 | 0.9 | 0 |
| EGFR(L747-E749del, A750P) | 52 | 2.2 | 17 | 4.2 | 4 | 0.25 | 2.6 | 3 | 1.1 |
| EGFR(L747-S752del, P753S) | 38 | 2.2 | 20 | 3.6 | 6.6 | 0.8 | 3.5 | 0.5 | 0.6 |
| EGFR(L747-T751del,Sins) | 43 | 3.4 | 14 | 1.1 | 25 | 0.4 | 0.85 | 1.6 | 0 |
| EGFR(L858R) | 84 | 3.2 | 22 | 2.4 | 1 | 0.25 | 1.6 | 4 | 0.1 |
| EGFR(L858R,T790M) | 94 | 100 | 0.35 | 6.4 | 1.8 | 100 | 72 | 100 | 77 |
| EGFR(L861Q) | 66 | 0.85 | 18 | 0.5 | 2.2 | 0 | 0 | 0.25 | 0 |
| EGFR(S752-I759del) | 90 | 0 | 11 | 1.8 | 56 | 0.75 | 2 | 1.4 | 0 |
| EGFR(T790M) | 90 | 31 | 0.8 | 0.2 | 1 | 13 | 39 | 68 | 9.9 |
| EIF2AK1 | 91 | 100 | 100 | 81 | 68 | 72 | 67 | 100 | 78 |
| EPHA1 | 87 | 50 | 79 | 61 | 0.3 | 0.6 | 72 | 16 | 2 |
| EPHA2 | 70 | 0.6 | 96 | 86 | 1.8 | 0.35 | 2 | 1.2 | 1.7 |
| EPHA3 | 45 | 15 | 100 | 100 | 40 | 0.75 | 19 | 5.7 | 4.8 |
| EPHA4 | 85 | 0.55 | 94 | 94 | 0 | 0.25 | 10 | 3.7 | 0 |
| EPHA5 | 87 | 8.4 | 100 | 96 | 1.1 | 0.85 | 9.8 | 21 | 3.6 |
| EPHA6 | 98 | 18 | 100 | 74 | 1.4 | 0.15 | 36 | 29 | 0 |
| EPHA7 | 97 | 79 | 62 | 78 | 0.6 | 0.1 | 100 | 78 | 16 |
| EPHA8 | 12 | 0.8 | 53 | 52 | 0.1 | 0 | 1.4 | 0.35 | 0.2 |
| EPHB1 | 100 | 3 | 84 | 86 | 0.2 | 0.35 | 10 | 11 | 0.5 |
| EPHB2 | 91 | 4.4 | 100 | 100 | 0 | 0.6 | 5.3 | 5.8 | 0 |
| EPHB3 | 99 | 35 | 100 | 88 | 6.6 | 2.7 | 19 | 63 | 4.7 |
| EPHB4 | 95 | 37 | 100 | 86 | 1.4 | 0.8 | 23 | 21 | 2.7 |
| EPHB6 | 76 | 21 | 44 | 14 | 2.8 | 0.85 | 12 | 2.8 | 1.8 |
| ERBB2 | 95 | 0 | 57 | 0 | 40 | 0.3 | 6.2 | 19 | 0 |
| ERBB3 | 98 | 100 | 53 | 0.1 | 1.3 | 89 | 63 | 100 | 100 |
| ERBB4 | 98 | 1.2 | 4.6 | 0 | 24 | 0.1 | 0.75 | 3 | 0.1 |
| ERK1 | 100 | 100 | 100 | 94 | 100 | 96 | 100 | 100 | 100 |
| ERK2 | 87 | 100 | 100 | 90 | 100 | 100 | 100 | 100 | 100 |
| ERK3 | 99 | 100 | 1 | 100 | 100 | 100 | 100 | 91 | 100 |
| ERK4 | 91 | 100 | 96 | 100 | 100 | 26 | 100 | 77 | 100 |
| ERK5 | 87 | 100 | 87 | 89 | 1.4 | 36 | 100 | 83 | 73 |
| ERK8 | 97 | 85 | 86 | 91 | 1.5 | 2.6 | 88 | 87 | 67 |

| | | | | | | | | | |
|---|---|---|---|---|---|---|---|---|---|
| ERN1 | 96 | 100 | 69 | 74 | 16 | 54 | 12 | 100 | 72 |
| FAK | 99 | 91 | 85 | 100 | 0.4 | 59 | 98 | 84 | 67 |
| FER | 40 | 56 | 45 | 100 | 1.3 | 2.6 | 43 | 21 | 27 |
| FES | 5.3 | 13 | 73 | 88 | 1.2 | 0.1 | 8.2 | 6.8 | 0.2 |
| FGFR1 | 55 | 1.2 | 2.2 | 36 | 4.6 | 0.15 | 0.95 | 2.8 | 0.1 |
| FGFR2 | 77 | 5.7 | 22 | 46 | 11 | 0.5 | 12 | 5.4 | 3.7 |
| FGFR3 | 93 | 28 | 35 | 43 | 21 | 0.15 | 32 | 13 | 1.8 |
| FGFR3(G697C) | 84 | 29 | 34 | 28 | 29 | 0.15 | 42 | 23 | 2.5 |
| FGFR4 | 51 | 4.4 | 60 | 84 | 81 | 0 | 15 | 8.1 | 0.5 |
| FGR | 14 | 0.75 | 4.2 | 3.2 | 0.8 | 0.15 | 1 | 1 | 0.2 |
| FLT1 | 36 | 10 | 54 | 60 | 2.2 | 0.8 | 9.2 | 12 | 2.4 |
| FLT3 | 7 | 0.5 | 3 | 18 | 7.6 | 0.15 | 0.2 | 0.15 | 1 |
| FLT3(D835H) | 52 | 7.3 | 10 | 14 | 9.6 | 0.35 | 3.8 | 2.6 | 0.2 |
| FLT3(D835Y) | 48 | 38 | 0.35 | 16 | 1.6 | 0.85 | 17 | 6.8 | 2.8 |
| FLT3(ITD) | 24 | 2.3 | 1.6 | 43 | 2.1 | 0.15 | 5 | 1.4 | 0.4 |
| FLT3(K663Q) | 4.2 | 1.6 | 2.4 | 22 | 5.1 | 0 | 0.55 | 0.15 | 0.1 |
| FLT3(N841I) | 20 | 1.5 | 1.2 | 23 | 100 | 0 | 0.75 | 2.3 | 0.3 |
| FLT3(R834Q) | 59 | 44 | 51 | 64 | 100 | 7.2 | 4.7 | 7.4 | 50 |
| FLT3-autoinhibited | 79 | n/a | 38 | 67 | 2.8 | n/a | 76 | n/a | n/a |
| FLT4 | 62 | 3.8 | 81 | 35 | 0 | 0.1 | 11 | 26 | 0.2 |
| FRK | 11 | 1.4 | 100 | 14 | 2.4 | 0.15 | 2.2 | 3.6 | 0.6 |
| FYN | 39 | 2.2 | 90 | 7.8 | 2.2 | 0.25 | 6 | 5.2 | 1.2 |
| GAK | 52 | 4.8 | 7.9 | 88 | 32 | 2 | 3.9 | 7 | 2.2 |
| GCN2(Kin.Dom.2,S808G) | 99 | 14 | 13 | 100 | 35 | 0.25 | 2.8 | 11 | 0.6 |
| GRK1 | 79 | 100 | 82 | 100 | 0.2 | 88 | 71 | 100 | 96 |
| GRK4 | 92 | 100 | 100 | 100 | 50 | 14 | 97 | 40 | 8.2 |
| GRK7 | 96 | 100 | 100 | 69 | 3.1 | 100 | 98 | 92 | 77 |
| GSK3A | 72 | 78 | 100 | 100 | 31 | 100 | 100 | 100 | 100 |
| GSK3B | 96 | 88 | 98 | 72 | 5.8 | 90 | 78 | 100 | 99 |
| HASPIN | 100 | n/a | 100 | 75 | 56 | n/a | 23 | n/a | n/a |
| HCK | 5.5 | 0.55 | 85 | 0.6 | 0.8 | 0.15 | 0.45 | 0.5 | 0.5 |
| HIPK1 | 21 | 51 | 52 | 72 | 0.5 | 48 | 62 | 74 | 74 |
| HIPK2 | 16 | 84 | 90 | 91 | 0.3 | 46 | 56 | 86 | 41 |
| HIPK3 | 18 | 55 | 74 | 75 | 0.8 | 25 | 51 | 80 | 44 |
| HIPK4 | 36 | 2.6 | 74 | 91 | 7.8 | 1.9 | 22 | 16 | 5.5 |
| HPK1 | 32 | 3.4 | 13 | 100 | 5.4 | 0.05 | 2.8 | 0.55 | 0 |
| HUNK | 86 | 79 | 78 | 81 | 6.2 | 21 | 100 | 57 | 18 |
| ICK | 79 | 100 | 100 | 98 | 0.5 | 96 | 54 | 96 | 99 |
| IGF1R | 95 | 100 | 62 | 83 | 31 | 95 | 100 | 100 | 100 |
| IKK-alpha | 50 | 0.2 | 85 | 91 | 11 | 0.45 | 0.6 | 2 | 0 |
| IKK-beta | 90 | 0.3 | 100 | 72 | 2.2 | 0.1 | 1 | 5.7 | 0.2 |
| IKK-epsilon | 98 | 94 | 100 | 100 | 19 | 92 | 100 | 100 | 94 |

| | | | | | | | | | |
|---|---|---|---|---|---|---|---|---|---|
| INSR | 100 | 92 | 17 | 57 | 8.8 | 53 | 100 | 78 | 79 |
| INSRR | 81 | 100 | 33 | 90 | 23 | 68 | 85 | 100 | 100 |
| IRAK1 | 76 | 62 | 53 | 74 | 3 | 6.2 | 100 | 90 | 26 |
| IRAK3 | 93 | 100 | 89 | 100 | 100 | 100 | 100 | 76 | 65 |
| IRAK4 | 81 | 94 | 94 | 66 | 1.2 | 64 | 86 | 100 | 48 |
| ITK | 98 | 75 | 0.85 | 6.5 | 0.3 | 2.2 | 100 | 81 | 18 |
| JAK1(JH1domain-catalytic) | 62 | 41 | 77 | 100 | 5.9 | 2.9 | 66 | 76 | 39 |
| JAK1(JH2domain-pseudokinase) | 100 | 93 | 0.75 | 100 | 10 | 74 | 100 | 88 | 41 |
| JAK2(JH1domain-catalytic) | 84 | 57 | 1.2 | 77 | 4 | 3 | 58 | 32 | 12 |
| JAK3(JH1domain-catalytic) | 45 | 2.4 | 0 | 0 | 2.4 | 3.5 | 12 | 55 | 1.1 |
| JNK1 | 28 | 72 | 2.4 | 72 | 0.1 | 0.45 | 100 | 77 | 19 |
| JNK2 | 0.35 | 1.8 | 13 | 60 | 0.2 | 0 | 9.2 | 6.6 | 0.1 |
| JNK3 | 52 | 75 | 4.6 | 77 | 0.1 | 1.8 | 77 | 76 | 15 |
| KIT | 12 | 0 | 20 | 44 | 2 | 0 | 0.15 | 0.3 | 0 |
| KIT(A829P) | 12 | 61 | 75 | 56 | 100 | 24 | 3 | 8 | 64 |
| KIT(D816H) | 79 | 43 | 83 | 82 | 77 | 8.2 | 18 | 25 | 36 |
| KIT(D816V) | 66 | 9.2 | 60 | 77 | 1 | 0.65 | 22 | 8.8 | 0.7 |
| KIT(L576P) | 4.5 | 82 | 8.9 | 23 | 77 | 0 | 0.8 | 0.15 | 0 |
| KIT(V559D) | 7.5 | 0 | 15 | 25 | 3.6 | 0 | 0.1 | 0.05 | 0 |
| KIT(V559D,T670I) | 16 | 0.85 | 28 | 98 | 9.2 | 0 | 2.8 | 4.9 | 0.1 |
| KIT(V559D,V654A) | 51 | 28 | 100 | 96 | 47 | 1.4 | 24 | 7.1 | 2.6 |
| KIT-autoinhibited | 93 | n/a | 39 | 73 | 1.2 | n/a | 78 | n/a | n/a |
| LATS1 | 100 | 100 | 77 | 55 | 4.6 | 3.4 | 100 | 80 | 32 |
| LATS2 | 100 | 98 | 40 | 95 | 3.2 | 7 | 100 | 100 | 79 |
| LCK | 2.6 | 1.1 | 32 | 0.4 | 0.5 | 0.1 | 0.3 | 0.35 | 0.2 |
| LIMK1 | 96 | 26 | 90 | 30 | 73 | 1.2 | 50 | 25 | 7.7 |
| LIMK2 | 94 | 32 | 100 | 97 | 100 | 0.35 | 58 | 26 | 2.9 |
| LKB1 | 96 | 100 | 76 | 100 | 24 | 60 | 83 | 58 | 7.9 |
| LOK | 0.35 | 0.75 | 100 | 40 | 0.1 | 0 | 0 | 0 | 0 |
| LRRK2 | 94 | 94 | 7.5 | 85 | 99 | 6.2 | 97 | 53 | 12 |
| LRRK2(G2019S) | 97 | 83 | 2.6 | 92 | 100 | 8.8 | 89 | 71 | 18 |
| LTK | 84 | 89 | 100 | 82 | 5.2 | 2.5 | 74 | 93 | 62 |
| LYN | 6.5 | 0.15 | 49 | 11 | 0.5 | 0.05 | 0.4 | 0.45 | 0 |
| LZK | 100 | 78 | 73 | 84 | 56 | 7.8 | 98 | 82 | 20 |
| MAK | 66 | 100 | 100 | 100 | 69 | 61 | 80 | 89 | 100 |
| MAP3K1 | 75 | 83 | 100 | 48 | 37 | 96 | 27 | 100 | 71 |
| MAP3K15 | 70 | 100 | 100 | 92 | 25 | 72 | 11 | 100 | 100 |
| MAP3K2 | 72 | 26 | 37 | 80 | 0.3 | 0.1 | 22 | 23 | 0.7 |
| MAP3K3 | 58 | 1.6 | 16 | 63 | 1.8 | 1.4 | 4.8 | 2 | 1.4 |
| MAP3K4 | 100 | 100 | 66 | 76 | 30 | 62 | 95 | 98 | 66 |

| | | | | | | | | | |
|---|---|---|---|---|---|---|---|---|---|
| MAP4K2 | 1.8 | 4.6 | 25 | 79 | 0.4 | 0 | 5.2 | 3.2 | 0 |
| MAP4K3 | 93 | 40 | 23 | 91 | 11 | 1.8 | 33 | 20 | 14 |
| MAP4K4 | 2.6 | 11 | 49 | 93 | 18 | 1.8 | 0.75 | 1.8 | 4.2 |
| MAP4K5 | 59 | 16 | 25 | 96 | 13 | 1.6 | 2.4 | 7.2 | 3.4 |
| MAPKAPK2 | 84 | 100 | 100 | 96 | 3.6 | 100 | 100 | 100 | 100 |
| MAPKAPK5 | 100 | 100 | 98 | 88 | 15 | 98 | 100 | 85 | 90 |
| MARK1 | 96 | 100 | 2.1 | 84 | 84 | 100 | 72 | 100 | 94 |
| MARK2 | 84 | 64 | 10 | 100 | 57 | 83 | 100 | 59 | 68 |
| MARK3 | 100 | 100 | 9.2 | 60 | 5.2 | 42 | 100 | 100 | 38 |
| MARK4 | 100 | 94 | 1.8 | 81 | 31 | 85 | 59 | 81 | 100 |
| MAST1 | 88 | 99 | 100 | 67 | 91 | 100 | 34 | 96 | 100 |
| MEK1 | 100 | 100 | 51 | 18 | 0.1 | 82 | 94 | 90 | 74 |
| MEK2 | 100 | 22 | 19 | 20 | 0 | 37 | 78 | 74 | 58 |
| MEK3 | 100 | 100 | 26 | 76 | 1 | 67 | 59 | 62 | 68 |
| MEK4 | 94 | 100 | 35 | 100 | 3.2 | 25 | 91 | 100 | 76 |
| MEK5 | 100 | 0.85 | 18 | 0.2 | 0.2 | 0.4 | 1.3 | 1.4 | 0.2 |
| MEK6 | 88 | 91 | 55 | 100 | 67 | 88 | 100 | 100 | 79 |
| MELK | 85 | 48 | 16 | 75 | 5.5 | 6.1 | 73 | 72 | 20 |
| MERTK | 100 | 30 | 70 | 90 | 1.8 | 0.5 | 46 | 88 | 9.2 |
| MET | 100 | 59 | 28 | 98 | 28 | 9.2 | 82 | 100 | 40 |
| MET(M1250T) | 99 | 61 | 39 | 89 | 33 | 59 | 98 | 74 | 26 |
| MET(Y1235D) | 96 | 83 | 23 | 84 | 18 | 4.9 | 100 | 94 | 79 |
| MINK | 12 | 33 | 41 | 88 | 1.6 | 5.4 | 10 | 24 | 2.8 |
| MKNK1 | 98 | 84 | 100 | 100 | 38 | 1.2 | 51 | 96 | 68 |
| MKNK2 | 74 | 7 | 100 | 53 | 2.2 | 0.6 | 44 | 48 | 2.7 |
| MLCK | 96 | 89 | 100 | 100 | 2.2 | 18 | 94 | 100 | 100 |
| MLK1 | 94 | 100 | 9 | 100 | 4.2 | 1.7 | 100 | 4.7 | 1.4 |
| MLK2 | 100 | 93 | 56 | 91 | 52 | 9.5 | 55 | 53 | 20 |
| MLK3 | 97 | 53 | 91 | 96 | 0.6 | 3 | 62 | 23 | 3.6 |
| MRCKA | 96 | 100 | 100 | 100 | 46 | 73 | 100 | 79 | 15 |
| MRCKB | 85 | 100 | 100 | 100 | 6.4 | 67 | 100 | 85 | 65 |
| MST1 | 100 | 89 | 100 | 74 | 3.4 | 12 | 100 | 72 | 21 |
| MST1R | 100 | 100 | 100 | 100 | 41 | 76 | 100 | 81 | 100 |
| MST2 | 93 | 83 | 86 | 57 | 100 | 26 | 41 | 100 | 5.2 |
| MST3 | 88 | 76 | 93 | 98 | 2.3 | 3.6 | 87 | 75 | 20 |
| MST4 | 75 | 98 | 49 | 62 | 0.4 | 11 | 25 | 87 | 18 |
| MTOR | 100 | 93 | 66 | 89 | 100 | 88 | 100 | 100 | 100 |
| MUSK | 48 | 1.4 | 8.8 | 97 | 1 | 0 | 3.9 | 2 | 0 |
| MYLK | 100 | 100 | 100 | 99 | 5.5 | 56 | 100 | 81 | 90 |
| MYLK2 | 100 | 0.85 | 86 | 85 | 7.4 | 0.15 | 13 | 4.6 | 0.6 |
| MYLK4 | 91 | 100 | 92 | 100 | 6.8 | 40 | 93 | 88 | 23 |
| MYO3A | 14 | 59 | 88 | 54 | 56 | 15 | 41 | 84 | 28 |
| MYO3B | 70 | 49 | 100 | 100 | 100 | 0 | 49 | 84 | 40 |

| | | | | | | | | | |
|---|---|---|---|---|---|---|---|---|---|
| NDR1 | 99 | 100 | 37 | 70 | 4.5 | 33 | 48 | 89 | 72 |
| NDR2 | 91 | 100 | 61 | 72 | 14 | 11 | 93 | 79 | 35 |
| NEK1 | 100 | 82 | 100 | 100 | 95 | 50 | 63 | 83 | 89 |
| NEK10 | 88 | n/a | 14 | 63 | 14 | n/a | n/a | n/a | n/a |
| NEK11 | 99 | 37 | 95 | 96 | 100 | 18 | 51 | 87 | 32 |
| NEK2 | 94 | 95 | 100 | 91 | 50 | 100 | 75 | 100 | 100 |
| NEK3 | 99 | 100 | 68 | 96 | 100 | 49 | 47 | 79 | 77 |
| NEK4 | 74 | 57 | 100 | 84 | 100 | 14 | 43 | 100 | 33 |
| NEK5 | 93 | 21 | 82 | 100 | 85 | 0.65 | 48 | 47 | 4.3 |
| NEK6 | 82 | 100 | 100 | 91 | 89 | 73 | 95 | 100 | 100 |
| NEK7 | 100 | 100 | 100 | 94 | 96 | 50 | 94 | 92 | 64 |
| NEK9 | 94 | 38 | 100 | 88 | 88 | 12 | 100 | 86 | 8.4 |
| NIK | 95 | n/a | 100 | 72 | 15 | n/a | n/a | n/a | n/a |
| NIM1 | 98 | 100 | 100 | 100 | 70 | 98 | 100 | 99 | 100 |
| NLK | 49 | 28 | 100 | 100 | 36 | 13 | 28 | 18 | 17 |
| OSR1 | 62 | 93 | 96 | 78 | 36 | 60 | 100 | 100 | 74 |
| p38-alpha | 0 | 0.25 | 100 | 100 | 41 | 0 | 0.9 | 1.7 | 0 |
| p38-beta | 1.4 | 0.35 | 100 | 100 | 44 | 0 | 6.4 | 0.3 | 0 |
| p38-delta | 55 | 66 | 100 | 100 | 5.6 | 2.6 | 100 | 67 | 22 |
| p38-gamma | 1.3 | 54 | 100 | 71 | 21 | 5.7 | 47 | 97 | 40 |
| PAK1 | 87 | 100 | 64 | 64 | 100 | 62 | 92 | 100 | 100 |
| PAK2 | 90 | 99 | 64 | 92 | 100 | 76 | 80 | 93 | 100 |
| PAK3 | 78 | 100 | 50 | 39 | 100 | 1.1 | 80 | 66 | 6 |
| PAK4 | 100 | 99 | 12 | 89 | 100 | 87 | 100 | 100 | 86 |
| PAK6 | 93 | 100 | 74 | 96 | 100 | 85 | 100 | 100 | 54 |
| PAK7 | 98 | 92 | 13 | 100 | 2.4 | 64 | 100 | 70 | 100 |
| PCTK1 | 93 | 100 | 100 | 98 | 0 | 46 | 100 | 100 | 98 |
| PCTK2 | 56 | 100 | 100 | 77 | 13 | 3.7 | 100 | 85 | 61 |
| PCTK3 | 76 | 100 | 97 | 78 | 2.7 | 12 | 100 | 84 | 100 |
| PDGFRA | 43 | 0.65 | 64 | 34 | 0.3 | 0 | 9 | 2.6 | 0.2 |
| PDGFRB | 6.8 | 0 | 6.9 | 33 | 1.8 | 0 | 0 | 0.1 | 0.1 |
| PDPK1 | 89 | 72 | 64 | 71 | 10 | 60 | 99 | 100 | 65 |
| PFCDPK1(P.falciparum) | 1.8 | 0.85 | 100 | 0 | 2.4 | 0.2 | 2.6 | 4.3 | 0.2 |
| PFPK5(P.falciparum) | 97 | 100 | 100 | 100 | 100 | 93 | 100 | 100 | 93 |
| PFTAIRE2 | 100 | 94 | 100 | 99 | 1 | 2 | 43 | 93 | 36 |
| PFTK1 | 77 | 100 | 100 | 81 | 1 | 1.6 | 98 | 93 | 72 |
| PHKG1 | 95 | 100 | 100 | 100 | 11 | 100 | 100 | 85 | 100 |
| PHKG2 | 99 | 73 | 72 | 84 | 100 | 100 | 88 | 100 | 100 |
| PIK3C2B | 100 | 100 | 100 | 63 | 2.6 | 100 | 79 | 91 | 100 |
| PIK3C2G | 100 | 100 | 100 | 80 | 3 | 100 | 89 | 100 | 82 |
| PIK3CA | 100 | 100 | 100 | 61 | 5.4 | 93 | 100 | 100 | 100 |
| PIK3CA(C420R) | 100 | 100 | 100 | 72 | 3 | 95 | 68 | 100 | 100 |

| | | | | | | | | | |
|---|---|---|---|---|---|---|---|---|---|
| PIK3CA(E542K) | 100 | 100 | 100 | 72 | 4 | 96 | 97 | 100 | 100 |
| PIK3CA(E545A) | 100 | 100 | 79 | 65 | 2.2 | 93 | 66 | 100 | 92 |
| PIK3CA(E545K) | 100 | 100 | 84 | 76 | 4.7 | 98 | 62 | 100 | 100 |
| PIK3CA(H1047L) | 100 | 100 | 100 | 54 | 84 | 100 | 76 | 98 | 100 |
| PIK3CA(H1047Y) | 100 | 100 | 100 | 65 | 36 | 82 | 71 | 71 | 90 |
| PIK3CA(I800L) | 84 | 100 | 100 | 75 | 35 | 100 | 94 | 100 | 100 |
| PIK3CA(M1043I) | 100 | 100 | 100 | 65 | 42 | 100 | 91 | 100 | 84 |
| PIK3CA(Q546K) | 91 | 100 | 100 | 83 | 6 | 100 | 99 | 100 | 98 |
| PIK3CB | 100 | 100 | 100 | 59 | 24 | 85 | 68 | 100 | 100 |
| PIK3CD | 100 | 100 | 100 | 71 | 55 | 76 | 81 | 100 | 78 |
| PIK3CG | 100 | 100 | 100 | 73 | 0.4 | 100 | 100 | 100 | 96 |
| PIK4CB | 92 | 100 | 100 | 100 | 0.3 | 100 | 42 | 100 | 91 |
| PIM1 | 98 | 100 | 100 | 94 | 87 | 100 | 100 | 100 | 100 |
| PIM2 | 97 | 100 | 100 | 89 | 2.1 | 100 | 90 | 100 | 100 |
| PIM3 | 99 | 93 | 81 | 94 | 75 | 100 | 100 | 100 | 100 |
| PIP5K1A | 90 | 77 | 13 | 97 | 81 | 10 | 100 | 21 | 2.6 |
| PIP5K1C | 52 | 100 | 57 | 13 | 95 | 100 | 0.9 | 100 | 84 |
| PIP5K2B | 80 | 90 | 100 | 83 | 100 | 7.7 | 100 | 12 | 0.4 |
| PIP5K2C | 76 | 96 | 63 | 92 | 100 | 100 | 20 | 100 | 100 |
| PKAC-alpha | 93 | 77 | 69 | 80 | 0.4 | 27 | 71 | 84 | 100 |
| PKAC-beta | 99 | 98 | 100 | 97 | 1.7 | 11 | 62 | 76 | 100 |
| PKMYT1 | 89 | 96 | 94 | 100 | 40 | 48 | 92 | 68 | 71 |
| PKN1 | 94 | 81 | 79 | 71 | 81 | 28 | 100 | 85 | 16 |
| PKN2 | 99 | 78 | 100 | 100 | 27 | 2.2 | 58 | 35 | 26 |
| PKNB(M.tuberculosis) | 86 | 90 | 58 | 68 | 0.1 | 98 | 100 | 84 | 89 |
| PLK1 | 88 | 100 | 100 | 90 | 0.1 | 56 | 100 | 98 | 100 |
| PLK2 | 89 | 100 | 78 | 77 | 2.2 | 100 | 100 | 97 | 60 |
| PLK3 | 89 | 100 | 100 | 69 | 32 | 92 | 100 | 100 | 83 |
| PLK4 | 88 | 100 | 23 | 44 | 0 | 97 | 83 | 86 | 59 |
| PRKCD | 58 | 78 | 100 | 95 | 6.3 | 24 | 51 | 60 | 49 |
| PRKCE | 82 | 100 | 63 | 100 | 3.7 | 78 | 100 | 62 | 75 |
| PRKCH | 83 | 100 | 100 | 95 | 3.2 | 79 | 95 | 100 | 100 |
| PRKCI | 100 | 73 | 65 | 74 | 15 | 40 | 70 | 100 | 100 |
| PRKCQ | 72 | 82 | 100 | 100 | 100 | 36 | 100 | 83 | 31 |
| PRKD1 | 100 | 90 | 52 | 70 | 6.3 | 63 | 100 | 100 | 3.5 |
| PRKD2 | 89 | 42 | 70 | 100 | 6.4 | 13 | 43 | 81 | 50 |
| PRKD3 | 96 | 67 | 54 | 100 | 100 | 6.4 | 72 | 93 | 52 |
| PRKG1 | 97 | 100 | 100 | 100 | 5.4 | 100 | 100 | 86 | 100 |
| PRKG2 | 95 | 100 | 100 | 54 | 5.1 | 77 | 100 | 82 | 91 |
| PRKR | 85 | 75 | 93 | 95 | 61 | 54 | 92 | 97 | 80 |
| PRKX | 91 | 78 | 100 | 100 | 100 | 19 | 100 | 57 | 75 |
| PRP4 | 87 | 100 | 90 | 80 | 100 | 84 | 87 | 85 | 12 |

| | | | | | | | | | |
|---|---|---|---|---|---|---|---|---|---|
| PYK2 | 84 | 21 | 66 | 96 | 4 | 0.15 | 25 | 19 | 0.8 |
| QSK | 87 | 100 | 100 | 98 | 0.8 | 77 | 52 | 100 | 100 |
| RAF1 | 99 | 8.9 | 86 | 70 | 100 | 0.95 | 25 | 7.8 | 0.9 |
| RET | 29 | 0.15 | 25 | 18 | 0.1 | 0 | 0.25 | 0 | 0.1 |
| RET(M918T) | 27 | 0.95 | 13 | 18 | 0.1 | 0 | 0.2 | 0.05 | 0 |
| RET(V804L) | 97 | 1.4 | 11 | 91 | 0 | 0 | 11 | 3.4 | 0.1 |
| RET(V804M) | 65 | 1 | 2.2 | 90 | 0 | 0 | 2.2 | 0.8 | 0 |
| RIOK1 | 100 | 100 | 49 | 77 | 98 | 53 | 95 | 20 | 0.8 |
| RIOK2 | 86 | 35 | 5.2 | 95 | 24 | 2 | 28 | 61 | 19 |
| RIOK3 | 93 | 100 | 53 | 69 | 90 | 38 | 73 | 27 | 0.4 |
| RIPK1 | 56 | 8.2 | 0.15 | 100 | 100 | 0 | 28 | 15 | 0.2 |
| RIPK2 | 82 | 5.3 | 100 | 5 | 28 | 0.3 | 9.6 | 4.5 | 1.8 |
| RIPK4 | 90 | 100 | 100 | 92 | 0.1 | 8 | 37 | 100 | 56 |
| RIPK5 | 100 | 86 | 33 | 9.2 | 1.8 | 8.2 | 100 | 91 | 36 |
| ROCK1 | 100 | 98 | 29 | 98 | 100 | 60 | 100 | 91 | 43 |
| ROCK2 | 82 | 94 | 26 | 86 | 100 | 18 | 100 | 88 | 34 |
| ROS1 | 100 | 100 | 100 | 100 | 12 | 34 | 97 | 100 | 100 |
| RPS6KA4(Kin.Dom.1-N-terminal) | 100 | 99 | 100 | 100 | 1 | 3.6 | 70 | 72 | 50 |
| RPS6KA4(Kin.Dom.2-C-terminal) | 94 | 100 | 79 | 84 | 0 | 87 | 78 | 73 | 30 |
| RPS6KA5(Kin.Dom.1-N-terminal) | 79 | 76 | 100 | 100 | 0.4 | 20 | 69 | 62 | 34 |
| RPS6KA5(Kin.Dom.2-C-terminal) | 99 | 100 | 100 | 79 | 18 | 97 | 89 | 100 | 100 |
| RSK1(Kin.Dom.1-N-terminal) | 83 | 96 | 72 | 83 | 5.2 | 34 | 95 | 91 | 31 |
| RSK1(Kin.Dom.2-C-terminal) | 99 | 61 | 100 | 91 | 34 | 17 | 91 | 50 | 20 |
| RSK2(Kin.Dom.1-N-terminal) | 94 | 100 | 100 | 73 | 0.1 | 2.8 | 68 | 89 | 14 |
| RSK2(Kin.Dom.2-C-terminal) | 100 | n/a | 100 | 100 | 3.2 | n/a | 100 | n/a | n/a |
| RSK3(Kin.Dom.1-N-terminal) | 95 | 42 | 66 | 89 | 7.7 | 0.7 | 91 | 68 | 2.7 |
| RSK3(Kin.Dom.2-C-terminal) | 96 | 73 | 100 | 100 | 7.5 | 87 | 83 | 67 | 42 |
| RSK4(Kin.Dom.1-N-terminal) | 90 | 100 | 45 | 100 | 0.1 | 28 | 81 | 99 | 42 |
| RSK4(Kin.Dom.2-C-terminal) | 91 | 28 | 100 | 89 | 7.2 | 29 | 69 | 50 | 15 |
| S6K1 | 100 | 55 | 100 | 63 | 1.6 | 3.2 | 44 | 46 | 9 |
| SBK1 | 100 | 93 | 10 | 68 | 49 | 87 | 68 | 94 | 57 |
| SGK | 90 | n/a | 26 | 54 | 0 | n/a | 100 | n/a | n/a |
| SgK110 | 96 | 96 | 100 | 95 | 100 | 42 | 100 | 45 | 41 |
| SGK2 | 100 | n/a | 100 | 68 | 1.8 | n/a | n/a | n/a | n/a |
| SGK3 | 98 | 100 | 74 | 100 | 1.1 | 11 | 48 | 90 | 38 |
| SIK | 87 | 1.8 | 100 | 93 | 1.3 | 0.25 | 1.2 | 2.1 | 0.9 |
| SIK2 | 87 | 34 | 57 | 100 | 48 | 18 | 52 | 26 | 30 |
| SLK | 63 | 9.8 | 77 | 76 | 1.5 | 0.3 | 2.1 | 13 | 0.6 |
| SNARK | 100 | 100 | 8.5 | 78 | 0.4 | 84 | 54 | 76 | 95 |

| | | | | | | | | |
|---|---|---|---|---|---|---|---|---|
| SNRK | 83 | 100 | 55 | 60 | 23 | 60 | 43 | 100 | 95 |
| SRC | 16 | 0.2 | 6.6 | 4 | 0.1 | 0.05 | 0.3 | 2.8 | 0.2 |
| SRMS | 54 | 8.4 | 100 | 0 | 3.4 | 0.2 | 30 | 34 | 0.7 |
| SRPK1 | 84 | 83 | 44 | 60 | 15 | 100 | 93 | 28 | 1.8 |
| SRPK2 | 86 | 100 | 98 | 86 | 74 | 63 | 100 | 76 | 28 |
| SRPK3 | 80 | 90 | 100 | 83 | 87 | 11 | 98 | 37 | 13 |
| STK16 | 93 | 80 | 0.75 | 99 | 46 | 47 | 100 | 100 | 54 |
| STK33 | 98 | 66 | 10 | 80 | 13 | 6.8 | 58 | 78 | 31 |
| STK35 | 60 | 91 | 100 | 50 | 10 | 0.8 | 73 | 65 | 23 |
| STK36 | 39 | 2.2 | 78 | 65 | 0.1 | 0.15 | 0.85 | 1.4 | 0.3 |
| STK39 | 73 | 94 | 85 | 52 | 66 | 100 | 56 | 70 | 100 |
| SYK | 24 | 18 | 63 | 100 | 3 | 1 | 14 | 18 | 7.2 |
| TAK1 | 0.4 | 6.6 | 3.6 | 64 | 6 | 0.25 | 11 | 1.2 | 4.8 |
| TAOK1 | 14 | 34 | 96 | 84 | 0.5 | 0 | 46 | 22 | 0.8 |
| TAOK2 | 3.7 | 18 | 65 | 82 | 0.3 | 0 | 53 | 16 | 0.5 |
| TAOK3 | 2.8 | 0.45 | 71 | 81 | 0.2 | 0 | 5.2 | 0.75 | 0 |
| TBK1 | 100 | 100 | 43 | 91 | 5 | 84 | 90 | 88 | 55 |
| TEC | 93 | 3.8 | 10 | 5.7 | 8.2 | 0.1 | 18 | 16 | 0.5 |
| TESK1 | 96 | 81 | 83 | 86 | 43 | 2.4 | 94 | 100 | 15 |
| TGFBR1 | 100 | 100 | 100 | 87 | 100 | 100 | 100 | 100 | 100 |
| TGFBR2 | 84 | 41 | 100 | 98 | 0.6 | 1.4 | 94 | 52 | 5.7 |
| TIE1 | 11 | 4.3 | 39 | 66 | 100 | 1.1 | 3.6 | 0.95 | 4.6 |
| TIE2 | 47 | 0.15 | 100 | 68 | 5.1 | 0 | 0.35 | 0 | 0.1 |
| TLK1 | 100 | 74 | 100 | 100 | 23 | 93 | 94 | 79 | 100 |
| TLK2 | 100 | 97 | 100 | 99 | 3.2 | 100 | 100 | 100 | 93 |
| TNIK | 9.7 | 11 | 20 | 100 | 5.1 | 1.6 | 15 | 14 | 3.2 |
| TNK1 | 93 | 15 | 0.95 | 99 | 3.7 | 0.8 | 39 | 6.6 | 1.4 |
| TNK2 | 100 | 4.1 | 100 | 27 | 1.4 | 0.6 | 4.8 | 5.4 | 0.8 |
| TNNI3K | 81 | 6.6 | 100 | 73 | 9.7 | 0 | 7 | 5.9 | 1 |
| TRKA | 82 | 6.6 | 30 | 79 | 0.6 | 0.3 | 16 | 54 | 2.5 |
| TRKB | 80 | 4.2 | 28 | 62 | 1.8 | 2 | 2.3 | 9.4 | 5.2 |
| TRKC | 73 | 8 | 61 | 88 | 0.6 | 0.35 | 21 | 47 | 1 |
| TRPM6 | 100 | 100 | 48 | 31 | 100 | 64 | 16 | 100 | 100 |
| TSSK1B | 99 | 100 | 25 | 82 | 100 | 98 | 78 | 90 | 100 |
| TTK | 84 | 64 | 4.8 | 73 | 100 | 2 | 78 | 56 | 6.8 |
| TXK | 100 | 2.1 | 4.6 | 1.4 | 1.4 | 0.25 | 3.2 | 1.4 | 0.7 |
| TYK2(JH1domain-catalytic) | 91 | 53 | 7.4 | 80 | 2.4 | 6.8 | 60 | 70 | 12 |
| TYK2(JH2domain-pseudokinase) | 92 | 100 | 68 | 100 | 32 | 96 | 100 | 47 | 100 |
| TYRO3 | 100 | 40 | 100 | 41 | 100 | 38 | 66 | 76 | 100 |
| ULK1 | 94 | 89 | 32 | 71 | 3.8 | 38 | 52 | 68 | 8.5 |
| ULK2 | 100 | 88 | 29 | 91 | 0.2 | 61 | 81 | 81 | 35 |

| | | | | | | | | | |
|---|---|---|---|---|---|---|---|---|---|
| ULK3 | 78 | 53 | 19 | 73 | 0.1 | 0.25 | 71 | 55 | 1.5 |
| VEGFR2 | 45 | 6.4 | 40 | 52 | 0.3 | 0.35 | 30 | 14 | 2.6 |
| VRK2 | 77 | 100 | 100 | 92 | 9.2 | 90 | 75 | 100 | 92 |
| WEE1 | 97 | 100 | 88 | 93 | 100 | 100 | 99 | 100 | 100 |
| WEE2 | 95 | 21 | 100 | 100 | 97 | 20 | 72 | 74 | 4.8 |
| WNK1 | 86 | n/a | 100 | 76 | 89 | n/a | 81 | n/a | n/a |
| WNK3 | 84 | n/a | 100 | 55 | 72 | n/a | 100 | n/a | n/a |
| YANK1 | 86 | 100 | 100 | 45 | 19 | 98 | 68 | 100 | 100 |
| YANK2 | 90 | 100 | 100 | 77 | 13 | 100 | 82 | 100 | 100 |
| YANK3 | 93 | 100 | 100 | 84 | 100 | 83 | 89 | 85 | 100 |
| YES | 40 | 0.55 | 17 | 2 | 1 | 0.25 | 2.7 | 2.2 | 0.7 |
| YSK1 | 79 | 87 | 89 | 96 | 12 | 12 | 67 | 100 | 100 |
| YSK4 | 98 | 0.45 | 0.45 | 95 | 0.2 | 0.1 | 2.8 | 1.1 | 0.2 |
| ZAK | 3.1 | 3.9 | 71 | 34 | 3.8 | 0.25 | 0.6 | 1 | 0.8 |
| ZAP70 | 72 | 69 | 0.2 | 100 | 31 | 28 | 45 | 98 | 52 |
